# TypeSINE: Genome-Wide Detection of SINE Retrotransposon Polymorphisms Reveals Functional Variants Linked to Body Size Variation in Pigs

**DOI:** 10.1101/2025.05.21.653756

**Authors:** Yao Zheng, Naisu Yang, Cai Chen, Wencheng Zong, Hong Chen, Shasha Shi, Xiaoyan Wang, Eduard Murani, Henry Reyer, Bo Gao, Longchao Zhang, Klaus Wimmers, Chengyi Song

## Abstract

The genotyping of Short Interspersed Element (SINE) retrotransposon insertion polymorphisms (RIPs) from large-scale sequencing data remains technically challenging. To overcome this limitation, we developed TypeSINE to genotype SINE-RIPs from short-read sequencing data. Analyzing 362 porcine genomes, we identified 749,835 SINE-RIPs (>85% accuracy), including 65,917 common variants (5-95% frequency). These showed uniform genomic distribution (∼22/Mb), with 40% in introns. Population analyses based on SINE-RIPs revealed independent domestication of Asian and European pigs from local wild populations, followed by introgression. GWAS detected body size-associated regions using SINE-RIPs. We created pigRIPdb (82,227 curated SINE-RIPs) with browsing and visualization tools. The method’s cross-species applicability is supported by conserved SINE evolutionary patterns, enabling polymorphism discovery of SINEs in livestock and humans. TypeSINE represents an efficient, scalable solution for genome-wide SINE-RIP analysis, advancing population genomics research.

**Teaser:** An integrated SINE-RIP genotyping platform with companion database enables large-scale analysis of SINE evolution and population genetics in swine

## Introduction

Retrotransposons, a type of transposable element or "jumping gene," replicate via an RNA intermediate and are extensively distributed in mammalian genomes, comprising over 40% of their genomic sequences (*1, 2*). Retrotransposons are primarily classified into two categories: long terminal repeat (LTR) retrotransposons and non-LTR retrotransposons (*2*), which include LINEs (Long Interspersed Elements) and SINEs. SINEs are non-coding DNA sequences found abundantly within mammalian genomes (*1*). Typically ranging from 100 to 300 base pairs in length, SINEs often originate from small cellular RNAs, such as tRNAs or 7SL RNA, which have acquired the ability to retrotranspose. They frequently possess an RNA polymerase III promoter sequence at their 5’ end, enabling transcription (*3*). SINEs are dispersed throughout mammalian genomes and can constitute a significant portion of the total DNA. For instance, they account for over 10% of the human and pig genomes and comprise about 20% of the rabbit genome (*4-6*).

Unlike LINEs, SINEs do not encode the proteins necessary for their own reverse transcription and integration. Instead, they rely on proteins encoded by autonomous elements like LINEs for transposition (*7*). Once integrated into the genome, SINEs are subject to duplication and mutation over evolutionary time. Importantly, SINE integrations can impact gene expression and genome evolution. They may function as alternative promoters, enhancers, or repressors, influencing both transcriptional and post-transcriptional processes (*4, 8-15*). While their presence can enhance genetic diversity and adaptability, it can also lead to genomic instability and diseases if integrated into or near critical genes (*16-19*).

Understanding SINE genomic integration events is crucial for comprehending genome structure, function, and evolution. Studying SINE evolution also sheds light on the mechanisms of genome dynamics and regulatory processes. SINE retrotransposon insertion polymorphisms are valuable tools in phylogenetic studies for tracking evolutionary relationships (*20-24*) and lineage-specific events across species. A database providing approximately 45,000 reference and non-reference retrotransposon insertion polymorphisms (RIPs) is available (*25*) in human, documenting all published RIPs generated by Alu (the most abundant retrotransposon families of SINEs in human), LINE1, and SVA retrotransposons. Additionally, a rice transposon insertion polymorphism (RTRIP) database, derived from a comprehensive mobilome profile of 60,743 transposable element loci using resequencing data from 3,000 rice genomes from 89 countries across five varietal groups, is available (*26*). However, similar databases are not yet available for livestock. In addition, in the past two decades, numerous pipelines have been developed for identifying retrotransposon insertion polymorphisms in humans (*27, 28*), and these can generally be applied to livestock. However, only a few, such as TypeTE (*29*), Alu-detect (*30*), AluMine (*31*), and polyDetect (*32*), are specifically designed to discover polymorphic SINE insertions in genomes. Additionally, their genotyping performance has not been systematically assessed using large-scale datasets.

To address these gaps, we developed a specialized SINE-RIP mining program called TypeSINE, capable of directly identifying SINE-RIPs from raw reads in next-generation sequencing (NGS) genomic data. We evaluated its performance in large-scale SINE-RIP genotyping using NGS data from 362 pig individuals. The obtained SINE-RIPs were systematically annotated and utilized for population genetic and GWAS analyses, identifying potentially functional genomic regions associated with body size. Additionally, we created the pigRIPdb database, which houses 82,227 SINE-RIPs and offers various primary functions such as RIP data browsing, visualization, searching, annotation, and sharing, to aid in the functional and evolutionary analysis of SINE-RIPs.

## Results

### Overview of TypeSINE

We present a novel program, TypeSINE, designed to detect SINE retrotransposon insertion polymorphisms directly from raw next-generation sequencing (NGS) data. The mining process comprises two primary pipelines: Pipeline A, which identifies potential polymorphic SINE insertions present in the current reference genome, but might be missing in the tested genomes, and Pipeline B, which identifies potential polymorphic SINE insertions in tested animal genomes that are not present in the reference genome (Fig. 1).

**Fig 1.**
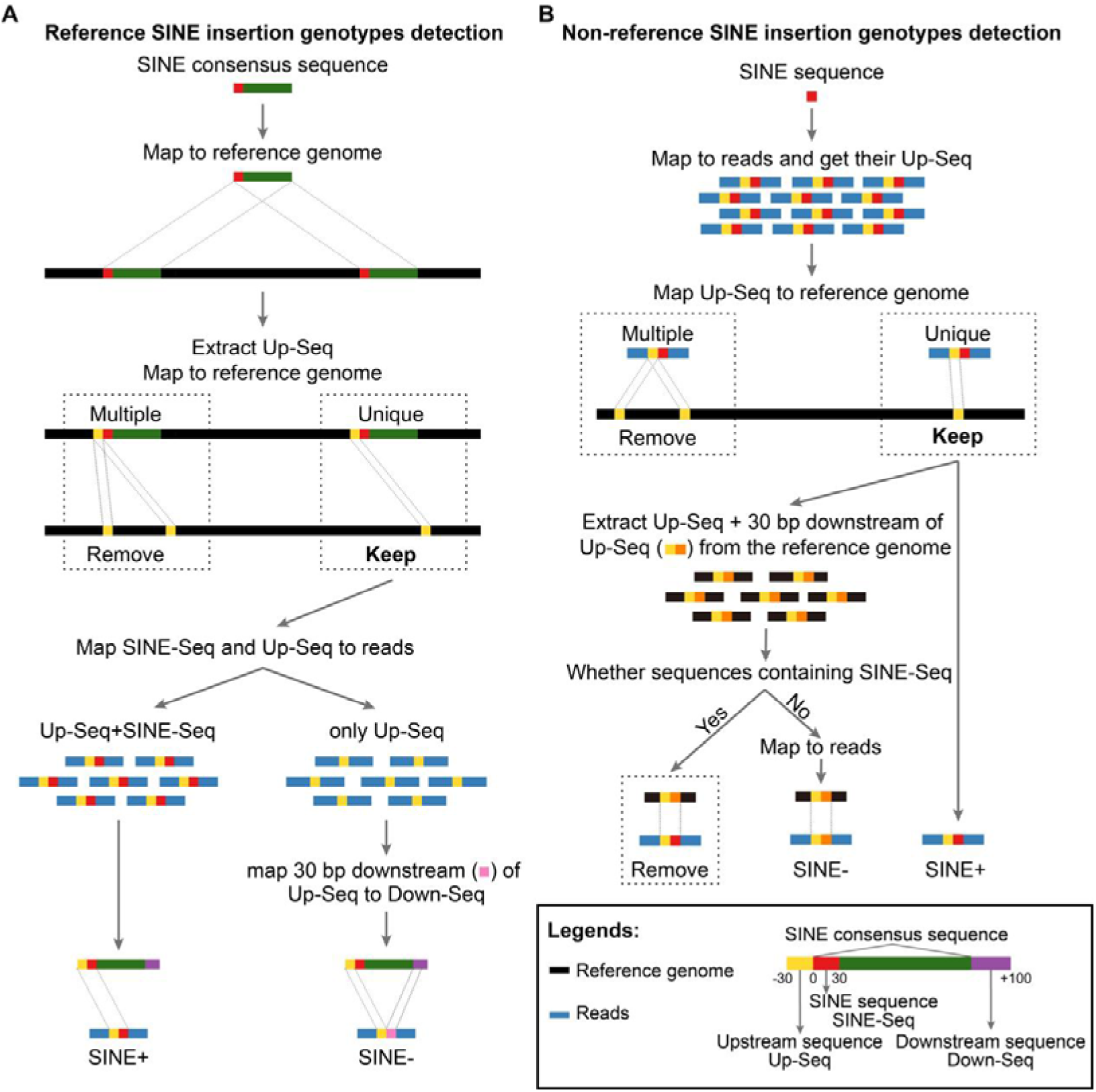
SINE RIP mining protocol. Illustration of the process for detecting SINE retrotransposon insertion polymorphisms (RIPs) using next-generation sequencing data. **(A)** Schematic diagram for reference SINE-RIPs detection. **(B)** Schematic diagram for non-reference SINE-RIPs detection.

Pipeline A assesses the presence or absence of SINE insertions at specific genomic sites in NGS data by utilizing SINE insertion coordinates annotated in the reference genome (Fig. 1A). Initially, the SINE consensus sequences (Table S1) were mapped to the reference genome by using the RepeatMasker program, then, 30 bp upstream sequence (Up-Seq) of each SINE insertion site were extracted and mapped to the reference genome, only unique coordinates were remained, and named as coordinates of unique Reference SINE insertions (uRefSINE+). Then, two sequence libraries are constructed based on the unique coordinates: one consists of 60 bp chimeric sequences, containing 30 bp from the 5’ end of the SINE sequence (SINE-Seq) and 30 bp from the upstream flank sequence (Up-Seq) at each insertion site, and the other consists solely of 30 bp from the upstream flank sequence (Up-Seq). These sequences are mapped to the raw sequencing data to determine whether reads contain the 60 bp chimeric sequence or only the 30 bp Up-sequence. Detection of the 60 bp chimeric sequence indicates the presence of a SINE insertion at that genomic site, consistent with the reference genome. Reads containing only the 30 bp Up-Seq are subjected to further analysis of their 30 bp downstream sequences. If this downstream sequence of reads can be mapped within a 100 bp region downstream of the corresponding SINE insertion site in the reference genome, the site is considered to be in an empty status of SINE (SINE^-^) relative to the reference genome (Fig. 1A).

Pipeline B employs a distinctive method to identify insertion polymorphisms of SINE retrotransposons that are missing from the reference genome (Fig. 1B). The first 30 base pairs of the 5’ end of SINE consensus sequences (SINE-Seq) from the most recent subfamilies (SINEA1-A3, table S1) are used as seed sequences to map reads containing SINE insertions. The 30 bp upstream genomic sequences (Up-Seq) of SINE-Seq within these reads are extracted and mapped to the reference genome, with only unique coordinates retained. Subsequently, 60 bp genomic sequences, including both 30 bp upstream and 30 bp downstream (Down-Seq) regions, are extracted from the reference genome. These sequences are aligned to SINE consensus sequences using BWA-mem2. Sequences containing SINE consensus (SINE-Seq) segments are discarded, while the remaining sequences are marked as SINE empty loci in the reference genome, indicating the presence of a novel polymorphic SINE insertion site, as inferred from the NGS data. These 60 bp SINE-empty genomic fragments are also mapped back to the NGS data to ascertain the allele status (SINE empty) at the corresponding positions. If these fragments are detected in the NGS data, it confirms that the reads share the same status (SINE empty) as the reference genome (Fig. 1B).

TypeSINE identifies SINE-RIPs from short reads data generated by next-generation sequencing technologies. Current pipeline can be used for individual genomes with more than 10x coverage, which generates sufficient and reliable RIPs, however, low coverage (less than10x coverage) might lose large number RIPs. Generally, we found that the TypeSINE program can detect all genotypes (SINE+/+, SINE+/-, SINE-/-) of individuals based on their raw sequencing reads for most mined RIPs, however, some RIPs were not detected in some individuals (designated as not available, NA) due to sequencing gaps. A comparative small-scale test (using sequencing data from 50 individual samples) was conducted for TypeSINE and MELT programs using identical computational resources. The total single-task processing times for TypeSINE and MELT were 99,597 and 93,106 minutes, respectively. Statistical analysis of the program outputs revealed that TypeSINE identified 191,218 SINE insertion polymorphic sites, where 53,932 sites showed SINE insertion allele frequencies between 0.05-0.95. In contrast, MELT identified 73,412 SINE insertion polymorphic sites, among which 41,930 sites exhibited SINE insertion allele frequencies between 0.05-0.95 (Table S2).

### Identification and evaluation of SINE-RIPs across 362 individuals of 79 breeds

Based on the TypeSINE protocol, totally, 817, 758 SINE-RIPs were identified from the genomes of wild boars and domestic pigs (Table S3 and S4), the RIPs were filtered if it was not detected in more than 75% of samples, finally, we obtained 749, 835 SINE-RIPs (Fig. 2A). However, majority of them (683,918) are rare RIPs, with SINE insertion allele (SINE^+^) frequencies (IAF) < 5% or > 95%, representing 91% of total RIPs, while the common RIPs (IAF ≥ 5 and ≤ 95%) only represent 9% of total RIPs (65, 917), as depicted in Fig. 2A and 2B. Furthermore, the old SINE-RIPs with almost fixed SINE insertion alleles (IAF > 0.95) account for 10.5% of total SINE-RIPs, while the new SINE-RIPs with very low SINE insertion alleles (IAF < 0.05) account for 80.6% of total SINE-RIPs, as depicted in Fig. 2C. The Venn diagrams revealed that most common SINE-RIPs (62, 189) are shared by wild and domestic pigs, only 3, 726 common SINE-RIPs are domestic pig specific, and two common SINE-RIPs are wild pig specific (Fig. 2D), while large number of rare SINE-RIPs (538, 318/71.7%) were mined from the genomes of domestic pigs, 84, 386 (11.2%) rare SINE-RIPs were shared by the genomes of wild boars and domestic pigs, and 61, 214 (8.1%) rare SINE-RIPs were only detected in the genomes of wild boars (Fig. 2E). PCR was used to evaluate the accuracy of SINE-RIP prediction by using 262 common SINE-RIPs, and more than 85% of them (234 RIPs) display polymorphisms (Fig. 2F and 2G, data S1), indicating the reliability of SINE mining protocol.

**Fig 2.**
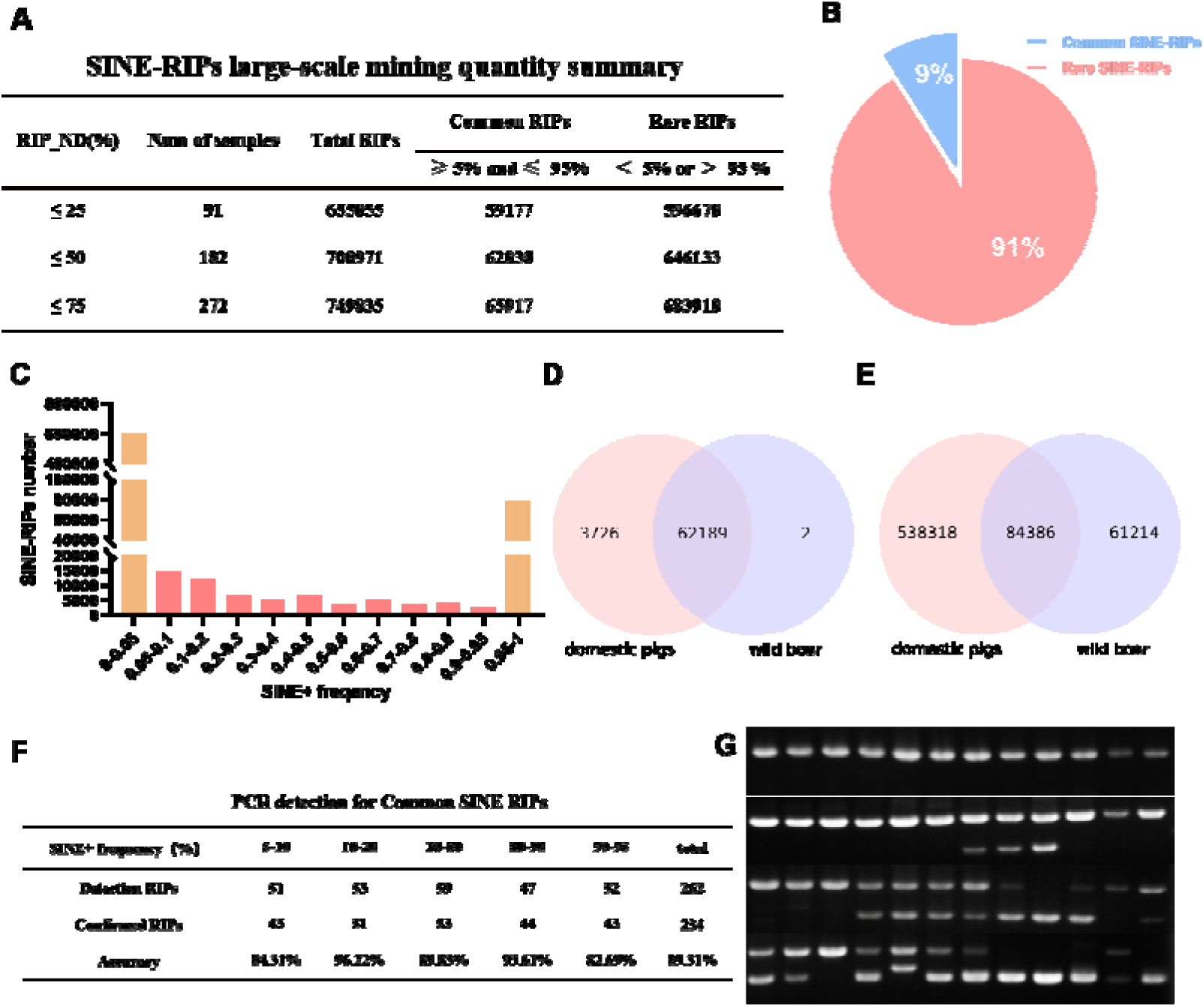
Identification and evaluation of SINE-RIPs across 362 individuals of pigs. **(A)** Summary of SINE-RIPs mined. **(B)** The percentages of common and rare SINE-RIPs. **(C)** The number of SINE-RIPs identified for different SINE+ (SINE insertion allele) frequencies. **(D)** Venn diagram of common SINE-RIPs. **(E)** Venn diagram of rare SINE-RIPs. **(F)** The summary of PCR detection results for 262 common SINE-RIPs. **(G)** Representative images of PCR detection for SINE-RIPs.

### Chromosomal distribution and Functional properties of SINE-RIPs

To evaluate the functional impact of SINE-RIPs, these RIPs were annotated using multiple pipelines (see Materials and Methods). Over 40% of the identified SINE-RIPs are located in intragenic regions. Specifically, 41.7% and 3.4% of SINE-RIPs are found within protein-coding genes and lncRNA genes, respectively, while 43.3% of common and 42.1% of rare SINE-RIPs reside in introns of these genes. In contrast, only 3.4% of common and 3.3% of rare SINE-RIP loci overlap with long non-coding RNAs (lncRNAs), 2% common SINE-RIPs and 2.4% rare SINE-RIPs are situated in the exons of protein-coding genes (Fig. 3A and 3B). Density plots and curves indicate that SINE-RIPs are widely distributed across the genome, with each genome containing 20-35 (mean 22.7) common SINE-RIPs and 240-300 (mean 258.5) rare SINE-RIPs per megabase (Mb) (see Fig. 3C and 3D). The sequences flanking genes exhibit a markedly distinct distribution of SINE-RIPs in proximal regions compared to distal regions. Both common and rare SINE-RIPs are significantly depleted immediately upstream and downstream of genes (within 2.5 kb), as depicted in Fig. 3E, indicating that proximal SINE-RIPs were under purifying selection. Within protein-coding gene regions, SINE elements exhibited directional insertion patterns. Forward-oriented SINE insertions constituted 42.7% (11,933) of the total common SINE-RIPs, while reverse-oriented insertions accounted for 57.3% (16,003). Spatial distribution analysis of SINE insertions relative to transcript coordinates revealed that reverse-oriented SINEs showed a tendency toward localization in the 5’ end of genes (Fig. 3F, data S2). The sequence logo revealed that the SINEs tend to integrate into 12 bp downstream of AT rich regions (Fig. 3G), and while the GC contents were significantly depleted in these regions (Fig. 3H).

**Fig 3.**
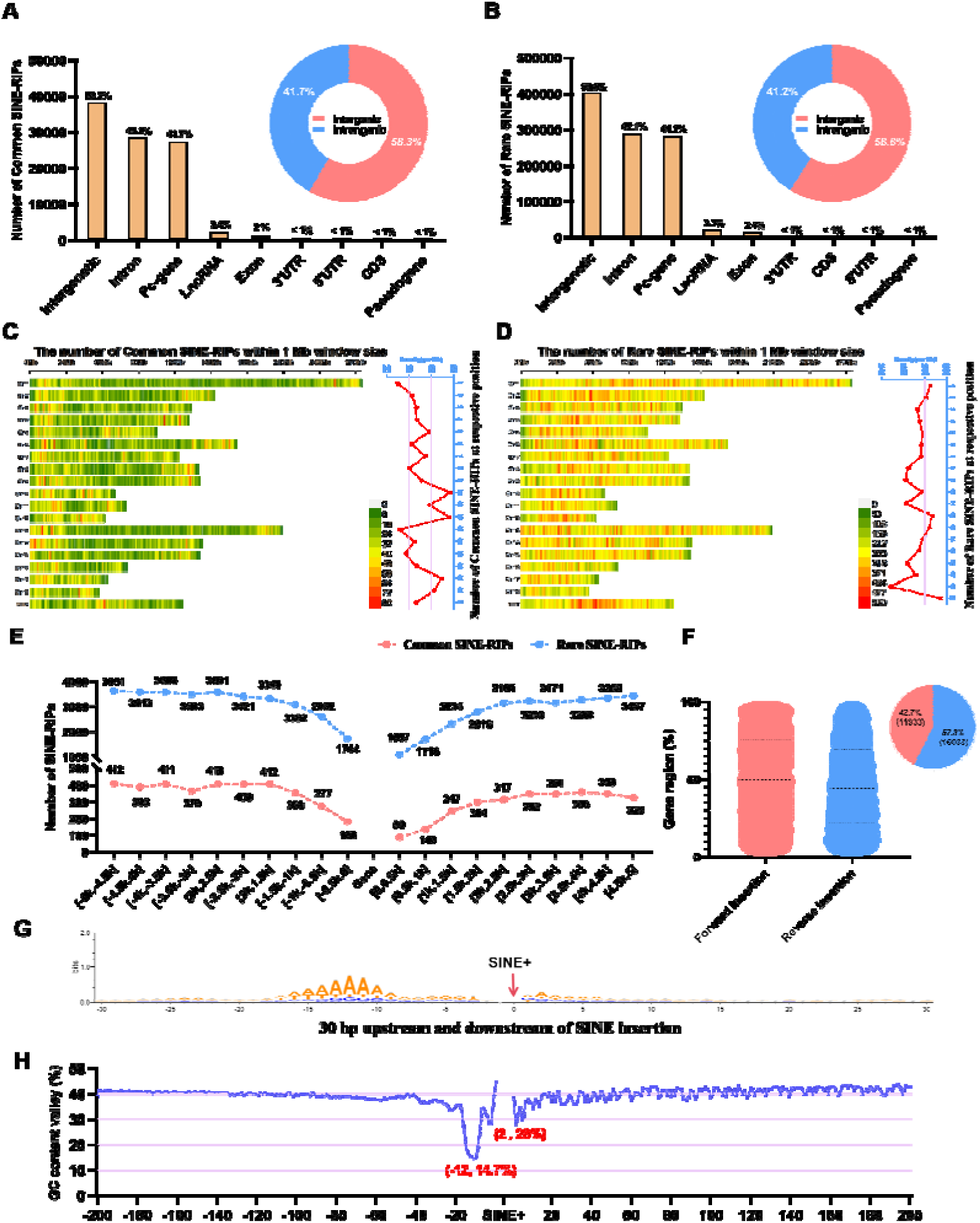
Chromosomal distribution and Functional properties of SINE-RIPs. **(A)** and **(B)** Histogram of the numbers of SINE-RIPs distributed in the genic regions and pie chart of the percentages of SINE-RIPs distributed intergenic and intragenic regions (A. for Common SINE-RIPs, and B. for Rare SINE-RIPs). **(C)** and **(D)** The number of Common SINE-RIPs within 1Mb window size and the density curve of its distribution on each chromosome (per Mb, C. for Common SINE-RIPs, and D. for Rare SINE-RIPs). **(E)** Common SINE-RIPs and rare SINE-RIPs landscape surrounding genes. The Gene on the X-axis is the location of the pc-gene, extending 5K bp upstream and downstream (each 500 bp is an interval), the Y axis is the number of SINE-RIPs. **(F)** Statistical analysis of SINE insertion orientation in common SINE-RIPs. The violin plot illustrates the regional distribution of SINE-RIPs within genes, with the Y-axis representing gene regions where "0%" indicates the transcription start site and "100%" denotes the transcription termination site. The pie chart presents the proportion of SINE insertions in the forward insertion (Red) versus the reverse insertion (Blue) relative to the transcriptional direction of the gene. **(G)** Integration sequence logo of Common SINE-RIPs. **(H)** The GC contents surrounding Common SINE-RIPs.

### Population genetic analysis based on common SINE-RIPs

The individuals were categorized into five major groups: European domestic pigs (ED), European wild boars (EW), Asian domestic pigs (AD), Asian wild boars (AW), and an Outgroup (OG) (Fig. 4A and B). Over 36% (23,979) of common SINE-RIPs were shared among all groups, while 22% (15,120) were shared by Sus species (EW, ED, AW, and AD). Rare SINE-RIPs unique to single groups were also identified, with EW, AD, and ED accounting for 2, 303, and 27, respectively (Fig. 4C, data S3). Principal component analysis (PCA) of common SINE-RIPs genotypes distinguished four Suidae species and separated Asian and European pigs (Fig. 4D-E, data S4). The Outgroup, including Babyrousa, Sus cebifrons, Sus verrucosus, and Porcula salvania, was clearly differentiated from Sus species (Fig. 4D). PC1 and PC2 (23.05% and 2.52%, respectively) revealed genetic divergence between Asian (AW and AD) and European (EW and ED) pigs (Fig. 4E). Notably, some Asian domestic pigs (e.g., Min pig, Qianshaohua pig, Yuedon black pig, and Putian pig) showed closer genetic affinity to Western pigs, likely due to human-mediated gene flow and introgression (Fig. 4E). Most Asian domestic pigs were genetically closer to Asian wild boars, while European domestic pigs aligned more closely with European wild boars (Fig. 4E). These findings, supported by phylogenetic and admixture analyses based on SINE-RIP genotypes (Fig. 4F-G, table S1), confirm that Asian and European domestic pigs originated from local wild boars and underwent independent domestication events. This aligns with previous SNP-based studies (*33, 34*). Genetic differentiation analysis further supports this, with lower Fst values between domestic pigs and wild boars from the same regions (Asian pairs: 0.049; European pairs: 0.04), indicating genetic connectivity. In contrast, higher Fst values were observed in trans-continental comparisons (European vs. Asian domestic pigs: 0.1648; European domestic pigs vs. Asian wild boars: 0.2115; Asian domestic pigs vs. European wild boars: 0.1914), reinforcing independent domestication events in Asia and Europe. European domestic pigs exhibited the highest differentiation with the Outgroup (0.2467), while Asian domestic pigs showed lower differentiation (0.1619), reflecting complex evolutionary relationships among populations (Fig. 4H-I).

**Fig 4.**
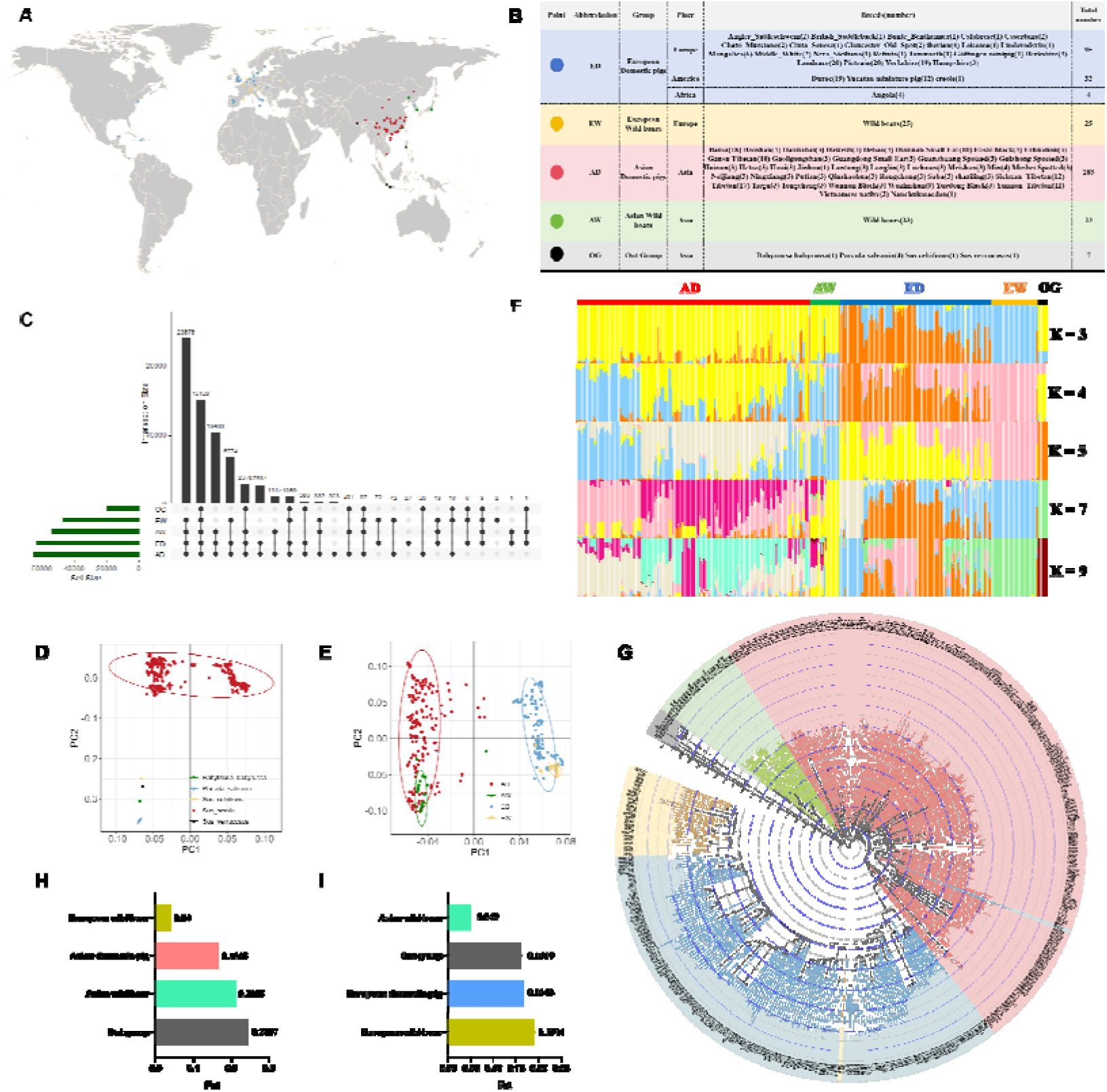
Population genetic analysis based on common SINE-RIPs. **(A)** Geographical distribution of the pig breeds included in this study and symbol description. **(B)** Table of ED (European domestic pigs), EW (European wild boars), AD (Asian domestic pigs), AW (Asian wild boars) and OG (Out group) information. **(C)** Venn diagram represents the distribution of common SINE-RIPs among different populations. **(D)** PCA plot displays the genetic relationship based on common SINE-RIPs among 369 individuals (Sus scrofa and out group). **(E)** PCA plot displays the genetic relationship based on SINE-RIPs among 362 individuals (Sus scrofa). **(F)** Genome wide admixture analyses inferred from Common SINE-RIPs (K = 3, 4, 5, 7 and 9) in 369 individuals (Sus scrofa and out group). **(G)** The phylogenetic tree was constructed based on Common SINE-RIPs for 369 individuals (Sus scrofa and out group). **(H)** The Fst results between European domestic pigs and other pig populations. **(I)** The Fst results between Asia domestic pigs and other pig populations. In figures (A), (B), (F), (E), (G), (H), and (I), different colors represent different populations, where blue represents European domestic pig populations, yellow represents European wild boar populations, red represents Asian domestic pig populations, green represents Asian wild boar populations, and black represents the outgroup.

The admixture analysis indicates that European domestic pigs and European wild boars are major ancestral components of European domestic pigs but minimal components of Asian domestic pigs. Conversely, Asian domestic pigs and Asian wild boars are major ancestral components of Asian domestic pigs but minimal components of European domestic pigs when K = 4 (Fig. 4F). This suggests that although Asian and European domestic pigs were domesticated independently, they experienced subsequent introgression processes.

Genetic differentiation among Chinese domestic pig populations revealed distinct patterns. At K=5, K=7, and K=9, Western Chinese populations split into Tibetan pigs and Yunnan-Sichuan pigs. At K=7, three more subgroups emerged: Southern, Southeastern-Central, and Eastern-Northern Chinese pigs. Notably, Tibetan pigs at K=4 and K=5 showed structural similarity to Asian wild boars (Fig. S1). Phylogenetic analysis of 369 individuals highlighted a branching pattern: Western Tibetan and Northern/Eastern pigs formed a transitional clade leading to Central Chinese pigs, which extended to Southern and Southeastern pigs (Fig 4G, fig. S2 and data. S5), aligning with population structure findings (Fig. S1).

### Association analysis of SINE-RIPs with body size

In this study, a genome-wide association study (GWAS) based on a mixed linear model (MLM) was performed to investigate the associations between common SINE-RIP genotypes and body size traits, including body weight, body circumference, body height, and body length. A total of 23 SINE-RIPs were significantly associated with body size. Specifically, 15 SINE-RIPs correlated with body weight (Fig. 5A and table S5), seven with body length (Fig. 5B and table S6), 17 with body height (Fig. 5C and table S7), and six with body circumference (Fig. 5D and table S8). The genotype frequencies of most of these RIPs showed significant differences across different categories of body size traits (Fig. S3 – S6). A total of 9, 5, 6, and 3 protein-coding genes (20 in total) contain SINE-RIPs associated with body weight, body length, body height, and body circumference, respectively (Fig. S3 – S6). Most of these RIPs were located in the introns of these genes, with six RIPs specifically situated in the first introns of genes (*PLAG1, LYN, PPARG, XPNPEP3, APPL2, and DNPEP*), which could influence gene expression, as first introns often contain regulatory elements (Table S9). Furthermore, three SINE-RIPs, including SRIP-17676 located in the first intron of *PLAG1*, SRIP-17680 within the first intron of *LYN*, and SRIP-30333 located in the fourth intron of *FRMD3*, (Fig. S3 and S4) showed significant associations with both of body weight and body length (*P* < 1 × 10^-5). These intronic SINE-RIPs showed strong statistical significance (*P* < 0.001) in genotype frequencies across different groups, as demonstrated by the Chi-square test, indicating potential regulatory roles in body size determination.

**Fig 5.**
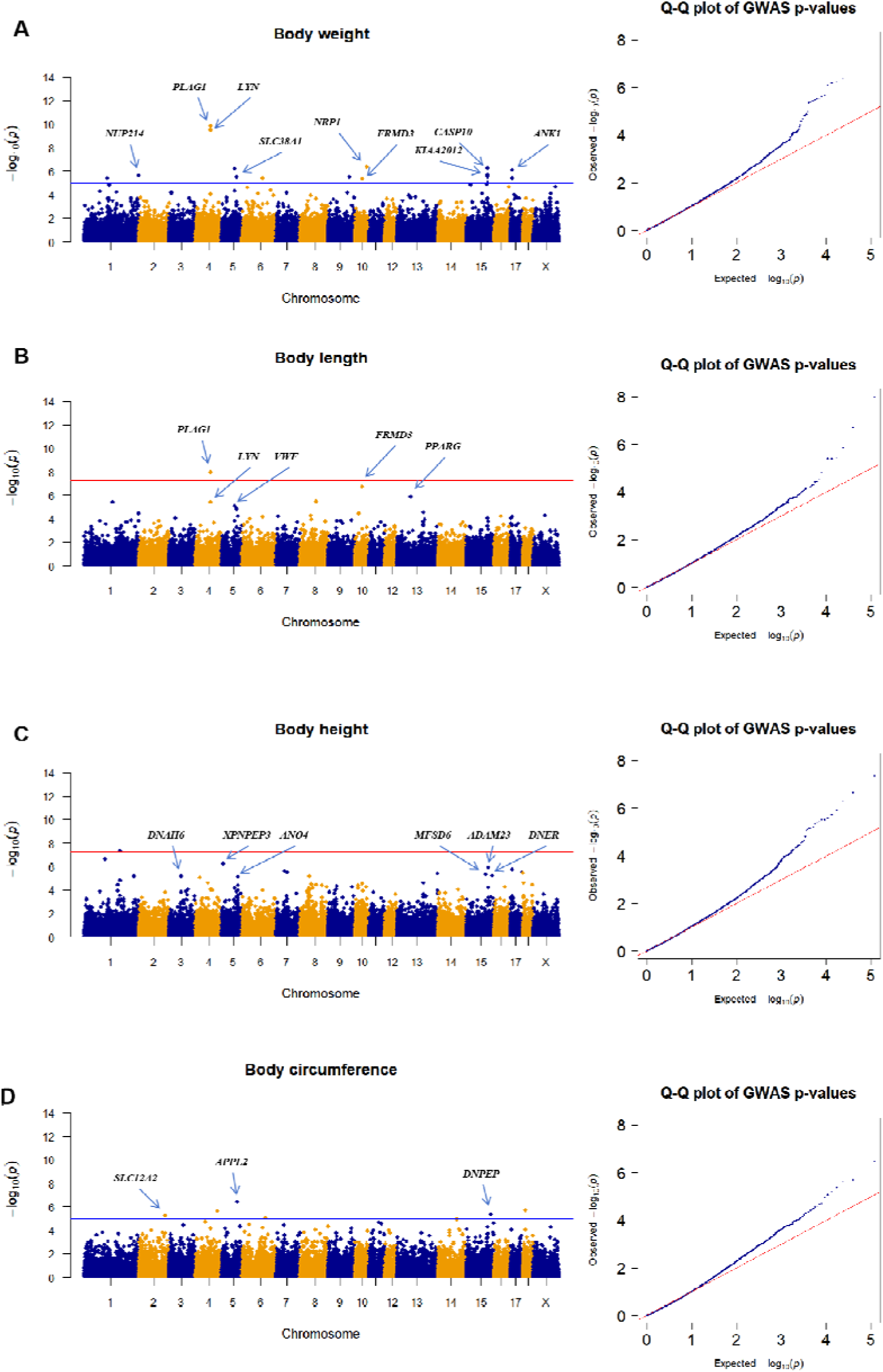
Manhattan plots and Quantile-Quantile (QQ) plots for four phenotypes of pig breed biological characteristics. Manhattan plots (left) and Quantile-Quantile plots (right) were plotted for body weight **(A)**, body length **(B)**, body height **(C)**, body circumference **(D)**. For Manhattan plots, the X-axis represents the chromosome 1-X, and the y-axis represents the −log_10_ (*P*) the P-values were corrected by the *P* < 1e-5 as cutoff, which was defined by blue line, and *P* < 3e-8 defined by red line, and arrow-marked points indicate significant SINE-RIPs residing within annotated genes.. For QQ plots, the X-axis represents the expected−log_10_(*P*), and the y-axis represents the observed −log_10_(*P*).

In addition, the analysis of intersections between common SINE-RIPs genomic regions (10 kb upstream and 10 kb downstream) and previously reported quantitative trait loci (QTL) associated SNPs (QTL-SNPs) obtained from Animal QTLdb (https://www.animalgenome.org/cgi-bin/QTLdb/SS/index) was conducted to identify potential associations of SINE-RIPs with body size traits. The overlapping SINE-RIPs constitute a small portion of the total common SINE-RIPs, accounting for less than 7%, ranging from 1.14% to 6.73%. In contrast, the overlapping SNPs represent approximately 40% (37.49-44.49%) of the total QTL-SNPs for each trait type, including meat and carcass traits, reproduction traits, health, production traits, and reproduction traits, as illustrated in fig. 6A and data S6. Additionally, 1,497 common SINE-RIPs genomic regions (2% of the total) overlap with the QTL-SNPs for ten growth traits. About half of these overlap with the QTL-SNPs for average daily gain, while 13%, 8%, 6%, 6%, and 4% overlap with the QTL-SNPs for body length, body circumference, body weight, days to 100 kg, and body height, respectively. Meanwhile, 13% overlap with QTL-SNPs for other growth trait types (Fig. 6B and 6C).

**Fig 6.**
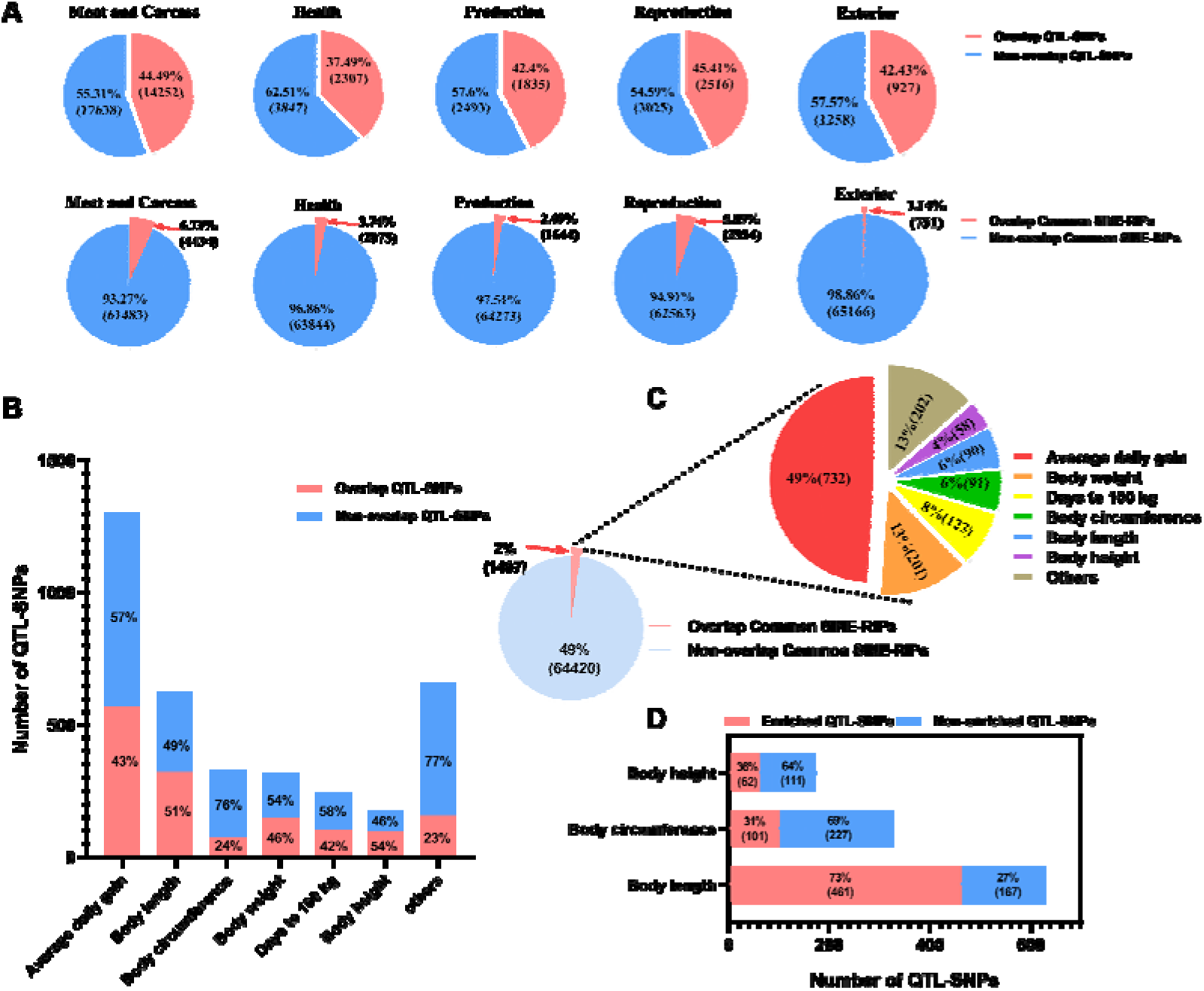
Intersection analysis of Common SINE RIP genomic regions with quantitative trait loci associated SNPs (QTL-SNPs). **(A)** Pie chart showing the percentage of overlap between QTL-SNPs (up) and Common SINE-RIPs (down) to the total numbers for each type of traits. **(B)** Histogram of the proportions of QTL-SNPs of growth traits overlapping with Common SINE-RIPs. **(C)** Pie chart of SINE-RIPs overlapping with QTL-SNPs of growth traits and their proportions for each type trait. **(D)** Histogram of the proportion of the proportions of QTL-SNPs of three body size traits (body length, body height, and body circumference) enriched for 34 SINE-RIPs.

To further explore the potential association between SINE-RIPs and body size traits, we examined the enrichment of QTL-SNPs related to body size (including width, depth, weight, circumference, length, and height) within 100 kb upstream and downstream genomic regions of common SINE-RIPs. We identified 34 SINE-RIPs that each enriched more than 50 QTL-SNPs associated with body size. Specifically, 73% (461) of QTL-SNPs for body length, 31% (101) of those for body circumference, and 36% (62) of those for body height were enriched within these regions (Fig. 6D and data S6). However, no significant enrichment was observed for QTL-SNPs linked to body width, depth, and weight. These 34 SINE-RIP form five clusters, located on chromosomes 3, 4, 5, 11, and 13. Each cluster contains several SINE-RIPs (4-9), closely distributed in specific genomic regions ranging from 296.94 kb to 554.79 kb (Fig. S7 and fig. S8). More specifically, the cluster on chromosome 3 comprises seven SINE-RIPs, collectively enriching 281 QTL-SNP for body length, spanning 554.79 kb. The cluster on chromosome 4 spans about 333.87 kb and contains four SINE-RIP, with each region enriching 82 or 83 QTL-SNP for body length. The chromosome 5 cluster, covering approximately 296.94 kb, includes six SINE-RIP, each enriching 61 or 54 QTL-SNP for body height. The chromosome 11 cluster contains nine SINE-RIP, spanning about 343.59 kb, with each region enriching 85, 96, or 101 QTL-SNPs for body circumference. Finally, the cluster on chromosome 13 consists of eight SINE-RIP and spans around 388.09 kb, with each region enhancing 58, 75, 96, or 97 QTL-SNP for body length (Fig. S7 and fig. S8).

Five genomic regions were further annotated for functional genes. In the SINE-RIP cluster regions on chromosome 4, five known protein-coding genes (*PLAG1, PENK, SDR16C5, MOS, RPS20*) were identified. On chromosome 5, four known protein-coding genes (*LALBA, KANLS2, WASHC4, APPL2*) and several unknown protein-coding genes were identified. On chromosome 11, four lncRNA genes were identified. A SINE-RIP (SRIP-17676) was located in intron 1 of *PLAG1*, and three SINE-RIPs (SRIP-19409, SRIP-19410 and SRIP-55826) were all found in introns 1, of *KANLS2*, respectively (Fig. S7 and Fig. S8). The genotype frequencies of these 34 SINE-RIPs varied significantly among different pig groups based on body size, including body length, height, and circumference. Specifically, the genotype frequency of SINE+/+ at the SRIP-17676 locus, located in intron 1 of *PLAG1*, was rare in pigs with short and medium body lengths but prevalent in those with long body lengths (Fig. S3). This locus was also significantly associated with body weight and length according to GWAS analysis (Fig. S3 and S4).

### Pig RIP database

After processing and redundancy removal, a total of 82,227 non-redundant SINE-RIPs were identified and subsequently made available through the pig RIP database (http://te.yzu.edu.cn/). These curated polymorphic loci represent a comprehensive catalog of SINE insertional polymorphisms in the pig genome, providing a valuable resource for genetic diversity studies and breed identification applications.

A comprehensive and user-friendly Pig RIPs database website was successfully developed through frontend optimization and structural refinement. The database provides free public access at http://te.yzu.edu.cn/, with RIP data available for download via a dedicated web interface at http://te.yzu.edu.cn/download/download_page/download.php. The integrated functional modules facilitate efficient access to and visualization of pig SINE insertion polymorphism data, supporting various research applications in pig genomics and breeding.

We integrated the main functions of pigRIPdb database (Home, RIPBrowser, RIPSeach, Blast, Download, Tools and Links, Help) into the browser such that it is conveniently accessible within most of the browsing windows via a standard button in the browser, as shown in Fig. 7A. The Jbrowse, which has gained wide used was deployed for the pig RIP database. We first installed the Jbrowse and the latest version of the pig genome database (Sscrofa11.1), and then added new codes and modified some existing codes for the browser to accommodate pig RIP database data as its tracks. The pig RIP database data is displayed tracks in the browser with each for polymorphic SINE elements, respectively. Since pig RIP database is integrated as a part of the browser, the RIP data can be queried using the utilities available from the browser. Being able to browse RIPs alongside genome annotation information offers a significant advantage by providing the users a graphic visualization of the genomic context of each RIP (Fig. 7B and C).

**Figure 7.**
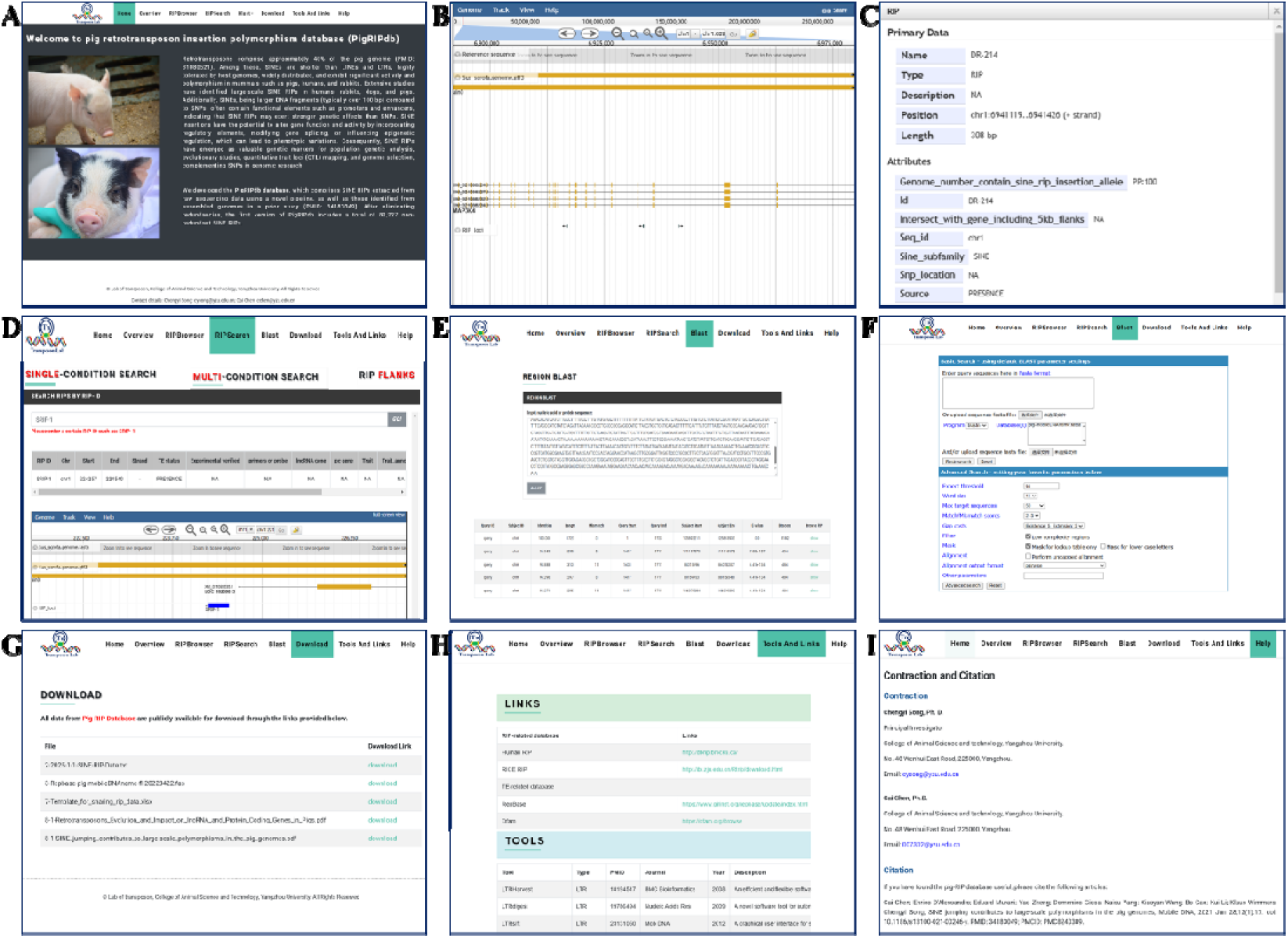
Screenshots of key features in pigRIPdb. **(A)** The “ Home” page interface of pigRIPdb. **(B)** and **(C)** The “RIPBrowser” module, which enables integration of SINE-RIPs with genome annotation data, and detailed information becomes accessible upon clicking the SINE RIP label. **(D)** The “RIPSearch” module, designed for users to retrieve specific SINE-RIPs identified in this study. **(E)** and **(F)** The “BLAST” module, allowing users to identify potential TEs or SINE-RIPs within query sequences. **(G)** The “Download” module, p(*3, 5*)roviding access to the pig RIPs dataset. **(H)** The “Tools and Links” page, offering some links of relevant database resources and analytical tools for pig and TE research. **(I)** The “Help” page, containing user guidance and other basic information.

To accommodate queries the pigRIPdb database, we implemented a special query interface called RIPSeach. RIPSeach is powered by the MySQL database engine. The query utilities of search are divided into the “SINGLE-CONDITION SEARCH”, “MULTI-CONDITION SEARCH”, “RIP FLANKS” three sections. There are three search methods (“Search RIPs by RIP-ID”, “Search RIPs by region”, “Search RIPs by gene”) in “SINGLE-CONDITION SEARCH” section and two search methods (“Search flanking RIPs of RIP-ID”, “Search flanking RIPs of Gene”) in “RIP FLANKS” section, respectively. “Search RIPs by RIP-ID” and “Search RIPs by gene” allows users to search by RIP IDs and gene name respectively, while “Search RIPs by region” allows users to search RIPs by chromosome region. “Search flanking RIPs of RIP-ID”, and “Search flanking RIPs of Gene” allows users to search the RIPs in the flank regions of a specific RIP and a specific gene respectively. In “MULTI-CONDITION SEARCH” section, we allow users search RIPs combine six filter conditions (“region”, “strand”, “RIP type”, “TE status”, “overlap with protein coding gene” and “overlap with lncRNA gene”) (Fig. 7D).

The BLAST search tool which contains two parts RIP Blast and basic Blast, has been deployed to determine whether the query sequences submitted by users encompass identified SINE-RIP loci or transposable elements, respectively (Fig. 7E and F). On the results page of BLAST search, each hit will be linked to the RIP interface. In this way, the user can know which RIP located to the matched segment of query sequences.

In addition to the main utilities described above, we also provide several other related resources on the pig RIP database. These include a complete list of downloadable data in flat files, all of which are accessed via the download menu bars at the top of the main page, which makes the entire pig RIPs data available that allows advanced users to perform systematic analysis of the large-scale RIP data (Fig. 7G and H). On the “Help” page, we provide some special instructions for using pig RIP database, which may not be so obvious to users (Fig. 7I).

## Discussion

### Evaluation and Application of SINE-RIPs

Significant differential mobilome profiles in abundance and composition were observed across livestock genomes, with genome coverages ranging from 26.1% to 42.9%, predominantly composed of LINEs, SINEs, and LTRs (*35*). LINEs constitute the largest components of these genomes, encompassing approximately 19% of the pig (*5*), 20% of the rabbit, 23% of the cattle (*36*), 27% of the goat (*37*), and 20% of the horse (*38*) genomes. LTRs account for 8% of pig, 5% of goat, 3% of cattle, and 4% of rabbit genomic sequences, which is substantially lower than the contribution of LINEs. The abundance of SINEs varies significantly among livestock, occupying about 11% of the pig, 15% of the rabbit, 12% of the goat, 7% of the horse, and 18% of the cattle genomes. Compared to LINEs and LTRs, SINEs are shorter, more tolerated by host genomes, widely distributed, and exhibit high activity and polymorphism in mammals such as pigs (*5*), humans (*24, 39, 40*) and rabbits (*6*). Analysis of data from 5,675 human genomes identified a total of 36,699 retrotransposon insertion polymorphisms (RIPs), primarily generated by Alu elements, a subtype of SINEs (26,553 instances). In contrast, RIPs mediated by LINE, SVA, and HERV-K elements numbered 7,353; 2,667; and 126, respectively. Another study identified Alu elements as major contributors to RIPs, finding 104,350 nonredundant RIPs across 57,919 human samples. Of these, 80,562 RIPs were mediated by Alu elements, while 16,525 were attributed to L1, 6,956 to SVA, and 307 to HERV-K (*40*). In pigs, including our own research and that of others, numerous polymorphic SINE loci have been identified. For example, Zhao et al. identified 211,067 SINE-RIPs (*41*), and Cai et al. reported over 3,600 SINE-RIPs (*20*). Large-scale polymorphic SINE loci have also been discovered in dogs (*42*). Furthermore, SINEs, being larger DNA fragments (generally over 100 bp) compared to SNPs, typically contain functional elements like promoters and enhancers (*8, 43*), suggesting that SINE-RIPs exert stronger genetic effects than SNPs. SINE insertions can alter gene function and activity by incorporating regulatory elements (*44, 45*), modifying gene splicing modes (*46*), or altering epigenetic regulation (*47*), leading to phenotypic variations (*46, 48, 49*), as extensively reported (*4*). Therefore, SINE-RIPs may serve as valuable genetic markers for population genetic analysis, evolution, QTL mapping, and even genome selection, complementing SNPs in genomic studies.

The developed TypeSINE pipeline enables the identification of a total of 749,835 SINE-RIPs from 297 domestic pigs and 48 wild pigs. Most of these are rare polymorphic SINE loci, with SINE insertion allele (SINE^+^) frequencies of less than 5% or greater than 95%. In contrast, the common SINE-RIPs (SINE^+^ frequencies between 5% and 95%) account for only 9% (65,917) of the total detected RIPs. SINE-RIPs are evenly distributed across the genome, with each genome containing approximately 22 common and 260 rare SINE-RIPs per megabase. Approximately 40% of SINE-RIPs are located within introns of protein-coding genes, indicating that they may have impact on these genes. Their effects on gene function and phenotypes merit further investigation, particularly for candidate genes associated with economically important traits.

Population genetic analyses using SINE-RIPs distinctly categorized domestic pigs into European and Asian groups, indicating independent domestication events in Europe and Asia. This finding is consistent with previous studies based on SNPs (*33, 34*), mtDNA (*50-53*) and ancient genomic DNA analysis (*54, 55*). However, ongoing admixture has been detected in both European and Asian domestic pigs, with minor ancestral components from European domestic pigs and wild boars present in Asian domestic pigs and vice versa. Overall, our RIP-based findings, in conjunction with SNP-based (*33, 34*), mtDNA-based (*50-52*), and ancient genomic DNA studies (*54, 55*), provide clear evidence of independent domestication of wild boar subspecies in Europe and Asia, accompanied by limited continual admixture and human-mediated introgression.

With the obtained SINE-RIPs, we also tried to evaluate their application potential in mapping the functional genomic regions and candidate genes of body size by using GWAS and by the intersection analysis between SINE-RIPs and QTL-SNPs of body size. The intersection analysis revealed that five genomic regions (located on chromosomes 3, 4, 5, 11, and 13) contain SINE-RIP clusters enriched QTL-SNPs of body size (> 50 QTL-SNPs for each RIP), suggesting these may be the functional genomic regions of body size. Furthermore, nine known protein-coding genes (*PLAG1*, *PENK*, *SDR16C5*, *MOS*, *RPS20*, *LALBA*, *KANLS2*, *WASHC4*, *APPL2*) in the SINE cluster regions of chromosome 4 and 5 were identified. While the GWAS analysis identified 42 SINE-RIPs significantly associated with body size, and 19 of them located in the introns of the protein-coding genes (*ADAM23*, *ANK1*, *ANO4*, *APPL2*, *DNAH6*, *DNER*, *DNPEP*, *FRMD3*, *KIAA2012*, *LYN*, *MFSD6*, *NRP1*, *NUP214*, *PLAG1*, *PPARG*, *SLC12A2*, *SLC38A1*, *VWF*, and *XPNPEP3*), and one RIP overlaps with the 3’UTR of protein-coding gene (*CASP10*). Our data demonstrated that *PLAG1* and *APPL2* were identified as candidate genes by both of GWAS and intersection analysis. and *PLAG1*, a zinc finger transcription factor gene, has been suggested as a candidate key gene regulating body size in previous studies (*56-58*), Rubin et al. reported that genotype combinations at two loci, *LCORL* and *PLAG1*, together explained a difference of 5.3 cm in body length in domestic pigs (*59*), and *PLAG1* is also found to be associated with bovine stature (*60*). These data strongly suggest that *PLAG1* may be involved in regulating body size, but the mechanism remains unknown. Future research is needed to validating how these SINE-RIPs, particularly SINE-RIP in intron1 of *PLAG1*, regulate target genes in specific tissues and affect phenotypes.

### SINE-RIP program

Numerous pipelines have been developed to mine retrotransposon insertion polymorphisms (structural variations) in genomes, including tools developed since 2020, such as TypeTE (*29*), HiTea (*61*), TEfinder (*62*), RetroSnake (*63*), MEHunter (*64*), T-lex3 (*65*), and TEMP2 (*66*). Despite the variety of available methods, MELT (*67*) remains the most popular due to its superior performance in identifying variants across all types of retrotransposons, such as LINE, SINE, and ERV, and because it currently is the only tool with continual support and documentation allowing for direct genotyping of both reference and non-reference RIPs. While most tools are designed to detect RIPs generated by all retrotransposon types, only a few, like TypeTE (*29*), Alu-detect (*30*), AluMine (*31*), and polyDetect (*32*), focus exclusively on SINE-RIP detection in humans. Notably, their performance in SINE-RIP genotyping has not been thoroughly evaluated. In this study, we integrated multiple bioinformatics tools and employed a modular design for high-throughput analysis, developing a novel methodology for the rapid discovery of polymorphic SINE insertions from raw sequencing data. Our method enables the direct genotyping of known and novel SINE insertions from raw reads, though it is unsuitable for use with assembled genomes or long-read sequencing data. Utilizing a robust PCR-based dataset spanning 200+ loci, we show TypeSINE achieves >80% genotyping accuracy, matching TypeTE’s performance (*29*). Although TypeSINE and MELT demonstrate comparable processing speeds for small datasets (n=50), MELT detects only 38% of SINE insertion polymorphic sites identified by TypeSINE. Since TypeTE relies on MELT-derived breakpoints (*29*), it similarly underestimates SINE polymorphism rates. The conserved evolutionary patterns of SINEs across mammals suggest TypeSINE’s broad applicability for polymorphism discovery of SINEs in diverse species, including livestock and humans. These findings establish TypeSINE as an efficient, scalable solution for genome-wide SINE-RIP analysis, advancing population genomics research.

### SINE-RIP Database

Besides the NCBI database, which houses both assembled pig genomes and unassembled raw sequencing data under ongoing updates, several other pig genome-related databases are available. The Pig Genome (https://www.sanger.ac.uk/data/pig-genome/), initiated by the Sanger Institute in the UK, provides access to maps, clones, and resources from the Porcine Genome Sequencing Project. Additionally, the pig pan-genome (http://animal.omics.pro/code/index.php/panPig) includes a highly assembled reference genome (2.4 Gb, Duroc genome, Sscrofa11.1) and 72.5 Mb of pan-sequences from five European commercial breeds and six Chinese local pig breeds. Furthermore, several integrated databases or portals, such as PigBiobank (http://pigbiobank.farmgtex.org), ISwine (http://iswine.iomics.pro/pig-iqgs/iqgs/index), and PigGTEx (http://piggtex.farmgtex.org), offer comprehensive knowledge bases by combining genome, transcriptome, genetic regulatory effects, and phenotypes. To date, two pig genetic variation databases have been released: PigVar (*68*) and 1KCIGP (*69*). The 1KCIGP database contains 50,590,445 SNPs and 12,870,753 INDELs, while the PigVar database is currently inactive. The 1KCIGP was released recently in 2023. These databases provide a rich dataset for the scientific community and are invaluable for understanding pig genomics, the regulation of complex traits, and phenotypes.

In the current study, we developed the pigRIPdb database based on SINE-RIPs mined from raw sequencing data using a newly developed pipeline, along with SINE-RIPs mined from assembled genomes in a previous study (*20*). The first version of the pigRIPdb curates 82,227 non-redundant SINE-RIPs from over 79 pig breeds (including five commercial breeds, 26 European breeds, and 40 Asian native breeds), Eurasian wild boars (13 individuals), and 18 assembled genome datasets (5 commercial breeds and 13 native breeds). pigRIPdb offers diverse primary functions: RIP data browsing, visualization, searching, annotation, and sharing. By integrating GWAS findings and gene and RIP annotations in the genome, pigRIPdb could aid in identifying candidate causal genes and variants underlying economically important traits in pigs. Additionally, the large-scale SINE-RIPs data could assist in the fine mapping of QTLs and their evaluation.

To the best of our knowledge, the pigRIPdb database is the first to archive large-scale retrotransposon insertion polymorphism data in the pig genome. While SINEs constitute only about 11% of genomic sequences—substantially lower than LINEs, which make up approximately 19%—they are more widely distributed in the pig genome compared to LINEs and ERVs (*5*). This broader distribution is attributed to their shorter length, which allows them to be more tolerated in the genome, making them promising genetic markers. Our lab continues to mine new SINE-RIPs using additional genomic sequencing data from public databases, as these resources expand and are regularly updated. Furthermore, we are also investigating RIPs generated by LINEs and ERVs, which may occur at lower frequencies than those generated by SINEs. We will consistently update the pigRIPdb with newly obtained RIPs. In future versions, pigRIPdb will encompass more SINE-RIPs from various populations and breeds, as well as LINE and ERV RIPs data. The platform will also feature user-friendly tools and a robust backend framework to enable interactive and real-time analysis. We anticipate that it will serve as a state-of-the-art, easy-to-use, and open-access resource for genomic variations mediated by retrotransposons in pigs.

## Conclusion

In this study, we developed TypeSINE, a tool designed to detect SINE retrotransposon insertion polymorphisms (SINE-RIPs) directly from raw next-generation sequencing (NGS) data. We identified 749,835 SINE-RIPs, with the majority (683,918) being rare RIPs, while common RIPs constitute only 9% of the total. More than 40% of SINE-RIPs are located in the introns of protein-coding genes. We also established pigRIPdb, a database containing over 82,227 SINE-RIPs, offering functionalities such as browsing, visualization, searching, annotation, and sharing to facilitate functional and evolutionary analyses of SINE-RIPs. Our SINE-RIP-based findings suggest that Asian and European domestic pigs originated from local wild boars and were domesticated independently, with limited admixture. Additionally, GWAS analyses identified genomic regions potentially associated with body size and indicated that *PLAG1* might be a key gene involved in regulating body size. Our findings underscore the significant roles of SINE-RIPs in the evolution of pig genomes and provide valuable insights for genetic and evolutionary studies in pigs.

## Materials and Methods

### Data and samples

We collected *Sus* whole genome next-generation (WGS) sequencing data with a sequencing depth > 10, from NCBI Sequence Read Archive (SRA, http://www.ncbi.nlm.nih.gov/sra/). A total of 362 *Sus scrofa* WGS-seq datasets and of 7 outgroup individuals (1 *Babyrousa babyrussa,* 1 *Sus cebifrons,* 1 *Sus verrucosus and* 4 *Porcula salvania*) were downloaded from European Nucleotide Archive (ENA, https://www.ebi.ac.uk/ena). Detailed samples information is provided in data S7.

### Data preprocessing and SINE-RIPs calling and genotyping

We developed TypeSINE, a computational tool for detecting and genotyping SINE retrotransposon insertion polymorphisms (SINE-RIPs) from next-generation sequencing data. The program is implemented in Python and relies on bioinformatics tools including BWA-MEM2(*70*), SAMTOOLS(*71*), and BEDTOOLS(*72*), employing a modular design for high-throughput analysis. Pipeline A performs SINE-RIP detection based on reference genome SINE insertion markers, utilizing a split-read strategy with stringent parameter settings. For sequence alignment using BWA-MEM2, we set the minimum seed length to k=15 to ensure specificity, minimum alignment score to T=50 to control false positives, and employed high mismatch penalty (B=4) and gap open penalty (O=4) to improve alignment accuracy. The program requires reads to completely span junction sites and determines genotypes based on upstream and downstream anchor sequence alignment results: homozygous insertion (1/1) when complete insertion sequence is detected, heterozygous (1/0) when both insertion and deletion signals are present, and homozygous deletion (0/0) when only deletion signal is detected. Pipeline B performs de novo detection of non-reference SINE insertions, we use the stringent parameters (k=15, T=30, B=1, O=1) to ensure only a single base mismatch or gap is allowed for seed sequence alignment. Reads matching seed sequences undergo special masking treatment: retaining at least 30 bp sequences at the upstream of matching seed sequences while replacing the matching region with N bases. These masked reads are then realigned to the reference genome using the same parameters. Candidate sites must meet the following criteria: the sequence length of reads aligned to the reference genome exceeds 20 bp, no hard clipping flags (indicating a unique alignment), the direction and coordinates of the sequences aligned to the reference genome and the seed sequences in the reads must be consistent and contiguous. (avoiding false positives), and support from at least 2 independent reads. The exact details of all mining process including pipeline A and B are described in the corresponding scripts available from GitHub (https://github.com/NaisuYang/TypeSINE).

In this study, we used the *Sus scrofa* 11.1 reference genome obtained from https://www.ncbi.nlm.nih.gov/datasets/genome/GCF_000003025.6/. The SINE insertion markers based on the reference genome were annotated using RepeatMasker (v 4.1.5) (*73*) with the following parameters: -e rmblast -pa 200 -s -cutoff 250 -no_is -nolow. FASTA files of all SINE consensus sequences used for annotation are deposited as table S1.

### Small-scale mining tests of TypeSINE and MELT programs

A small-scale SINE-RIP mining test was conducted using both TypeTE and MELT programs on sequencing data from 50 individual samples with an average sequencing depth of 37X (Data S8). Under identical computational resource conditions, we documented the processing time required for each single task during data processing for both TypeTE and MELT. Based on the final output from both programs, we extracted the SINE-RIPs sites and performed statistical counting analyses.

### PCR detection for common SINE-RIPs

We stratified 65,917 common SINE-RIPs into five frequency groups based on their insertion allele frequencies (IAF): 5% ≤ IAF < 10%, 10% ≤ IAF < 20%, 20% ≤ IAF < 80%, 80% ≤ IAF < 90%, and 90% ≤ IAF ≤ 95%. For experimental validation, we selected 262 SINE-RIPs with flanking sequences of 500 bp upstream and downstream. PCR primers were designed using NCBI Primer-BLAST (Primer-BLAST, https://www.ncbi.nlm.nih.gov/tools/primer-blast/index.cgi?LINK_LOC=BlastHome) with default parameters, optimizing for amplicon sizes between 250-1000 bp. The validation panel comprised 12 diverse pig breeds: four commercial breeds (Duroc, Large White, Landrace, and Pietrain), seven indigenous Chinese breeds (Meishan, Erhualian, Bama, Qingping, Min, Wuzhishan, and Sichuan Tibetan), and wild boar. For each breed, equimolar DNA from three unrelated individuals was pooled to create representative breed-specific templates. PCR amplification was performed, followed by agarose gel electrophoresis to evaluate SINE-RIP polymorphisms across the breed panel.

### Annotation of SINE-RIPs to genomic features

We retrieved the comprehensive annotation files for the *Sus scrofa* genome assembly 11.1 (Sscrofa11.1) from the Ensembl database (release 113; https://ftp.ensembl.org/pub/release-113/gff3/sus_scrofa/Sus_scrofa.Sscrofa11.1.113.gff3.gz). To characterize the genomic distribution of SINE-RIPs, we performed systematic intersection analysis using BEDTOOLS (v2.30.0) with the -intersect command on a UNIX platform. Both common (MAF ≥ 0.05) and rare (MAF < 0.05) SINE-RIPs were overlapped with various annotated genomic features. We quantified the distribution of SINE-RIPs across different genomic regions, including intergenic regions (defined as sequences between genes), intragenic regions (encompassing all gene-associated sequences), protein-coding genes (pcgenes), exons, introns, 5’ untranslated regions (5’ UTR), 3’ untranslated regions (3’ UTR), coding sequences (CDS), long non-coding RNAs (lncRNA), and pseudogene regions. For overlapping features, we implemented a hierarchical classification system where each SINE-RIP was assigned to its most specific applicable category.

### SINE-RIPs overlapping chromosome and insertion characteristics

We utilized the R package CMplot (v4.2.0) to generate high-resolution visualization of SINE-RIP distribution density across chromosomes. Our analysis focused on the genomic positions of both common (n = 63,191, IAF ≥ 0.05) and rare (n = 635,967, IAF < 0.05) SINE-RIPs located on autosomes and the X chromosome. To investigate the distribution patterns around protein-coding genes, we extracted genomic coordinates extending 5,000 bp upstream and downstream of annotated gene boundaries from the *Sus scrofa* reference genome (Sscrofa11.1). Using BEDTOOLS (v2.30.0) with the -intersect command, we calculated the coverage density of common and rare SINE-RIPs in consecutive 500 bp intervals across these regions.

We quantified common SINE-RIPs within protein-coding genes and recorded the number of SINEs in forward and reverse insertion relative to the gene’s transcriptional direction. Furthermore, we calculated the distance (L) between the SINE insertion start site (SS) and the gene transcription start site (GS) using the formula L = |SS – GS|. The relative position of each SINE insertion within the gene was then determined by computing the ratio of this distance (L) to the total gene transcript length (GL), expressed as a percentage: Relative Position (%) = (L/GL) × 100%.

To characterize the sequence context of SINE insertion sites, we employed BEDTOOLS -flank and -getfasta commands to extract the 30 bp flanking sequences on both sides of common SINE-RIPs. These sequences were then analyzed for nucleotide composition features using the WEBLOGO platform (v3.7.4; https://weblogo.berkeley.edu/logo.cgi) (*74*) with default parameters. To characterize the broader sequence context, we extracted 200 bp upstream and downstream sequences surrounding common SINE-RIPs and developed a custom perl script to compute the base-by-base GC content across these regions. The GC content was calculated using a sliding window approach with 1 bp steps.

### Map annotation and common SINE-RIP cross-statistics between different groups

We classified all pig breeds into five major population groups based on their ancestral geographical origins: Asian domestic pigs, Asian wild boars, European domestic pigs, European wild boars, and outgroups, and Angora native pigs, Duroc, Yucatan miniature pigs, and Creole pigs, were classified as European domestic pigs. To visualize the geographical distribution of these breeds, we utilized the R packages ggmap (v3.0.0) and maptools (v1.1.4) with breeds geographical coordinates (latitude and longitude) obtained from historical records and literature (Table S10). For population-specific SINE-RIP analysis, we examined 65,917 common SINE-RIPs across all five populations. Custom Python scripts were developed to identify SINE-RIPs with non-zero insertion allele frequencies in each population. The intersection patterns of these SINE-RIPs among the five populations were visualized using the R package UpSetR (v1.4.0) (*75*), which generates combination matrix layouts to display complex intersection patterns more effectively than traditional Venn diagrams.

### Analysis of genetic admixture

We performed population structure analysis using ADMIXTURE software (v1.3.0) (*76*) based on the genotype matrix of common SINE-RIPs. To determine the optimal number of ancestral populations, we conducted multiple analyses with varying numbers of genetic clusters (K) ranging from 1 to 10 across 369 individual pigs. Cross-validation error was calculated for each K value to assess model fit (Table S11). The analysis was performed using default parameters with a convergence threshold of 0.0001 and 1,000 bootstrap replicates. Population structure results were visualized using the R packages factoextra.

### Principal component analysis

Principal component analysis (PCA) was performed to investigate the population structure based on 65,917 common SINE-RIPs genotypes. Initially, we used VCFTOOLS (v0.1.16) to convert the variant call format (VCF) files into PLINK-compatible binary format. The data conversion was performed using default parameters while maintaining genotype quality information. Subsequently, we conducted PCA using PLINK (v1.9) (*77*) with the --pca command, calculating the first 5 principal components. The resulting principal components were visualized using the R package ggplot2 (v3.4.0), with the first two principal components plotted to reveal major population structure patterns.

### Fst analysis

To identify Common SINE-RIPs differentiation among populations, we calculated fixation indices (Fst) using VCFTOOLS (v0.1.16). Pairwise Fst analyses were conducted between Asian domestic pigs and four comparative groups (Asian wild boars, European domestic pigs, European wild boars, and outgroups). Similarly, we calculated pairwise Fst values between European domestic pigs and the same four comparative groups (Asian domestic pigs, Asian wild boars, European wild boars, and outgroups). For each pairwise comparison, the mean Fst value across all SINE-RIP was calculated to quantify the overall genetic differentiation between populations.

### Phylogenetic tree

We performed phylogenetic analysis using a dataset of 65,917 common SINE-RIPs. The genotype data was first converted to .VCF files, which were then used to calculate pairwise genetic distances between individuals using VCF2Dis v1.50 (https://github.com/hewm2008/VCF2Dis) with default parameters. The resulting distance matrix was converted to Newick (.NWK) format using the Neighbor-Joining (NJ) tree algorithm implemented in FastME 2.0 (*78*) through the FastMe bioinformatics platform (www.atgc-montpellier.fr/fastme/). The phylogenetic analysis included 362 *Sus scrofa* individuals and 7 outgroup specimens. The final phylogenetic tree was visualized and annotated using the Interactive Tree of Life (iTOL v6.4, https://itol.embl.de/) platform, with bootstrap support values displayed for major clades.

### Phenotype definition

We characterized four key morphological phenotypes based on breed-specific features: body weight, body length, body height, and body circumference. Phenotypic data were systematically collected from multiple authoritative sources, including ‘Animal Genetic Resources in China (Pigs)’, Wikipedia, peer-reviewed publications on breed genetic resources, and official breed registry databases (Data S9). And breeds lacking records for specific characteristic features were marked as missing. The four phenotypes are defined as follows:

Body weight is divided into five categories: small (<70 kg), small-medium (>70 kg and < 100 kg), medium (>100 kg and < 150 kg), medium-large (>150 kg and < 200 kg), and large (>200 kg), which are coded as 1, 2, 3, 4, and 5 in the phenotypic variable file.

Body length is divided into three categories: short (<100 cm), medium (>100 cm and < 140 cm) and long (>140 cm), which are coded as 1, 2, and 3 in the phenotypic variable file.

Body height is divided into three categories: short (< 50 cm), medium (> 50 cm and < 80 cm) and tall (> 80 cm), which are coded as 1, 2, and 3 in the phenotypic variable file.

Body circumference is divided into three categories: short (<100 cm), medium (>100 cm and < 120 cm) and long (>120 cm), which are coded as 1, 2, and 3 in the phenotypic variable file.

It is important to note that phenotypic data were not directly measured from the 362 individual pigs. Instead, phenotype classifications were assigned based on breed-level trait descriptions obtained from authoritative sources. Each individual was labeled according to the typical morphological characteristics of its breed, making this a breed-informed rather than individual-measured GWAS

### GWAS

In this study, we conducted a GWAS to examine the association between common SINE-RIPs genotypes and phenotypic traits. Association tests were performed using a MLM in the EMMAX-intel64 (*79*) program (https://genome.sph.umich.edu/wiki/EMMAX#Citing_EMMAX) on UNIX. Prior to analysis, a total of 65,917 common SINE-RIP loci were pre-processed by retaining only those located on chromosomes 1–18 and the X chromosome. Loci with a heterozygous SINE insertion frequency greater than 80% among 362 *Sus scrofa* individuals were filtered out, resulting in the retention of 598,142 common SINE-RIPs. Genotype data for these individuals were converted into VCF format, where homozygous insertions were denoted as 1/1, homozygous deletions as 0/0, heterozygous insertions as 1/0, and undetected genotypes as ./.. Subsequently, VCFTOOLS and PLINK were used to convert the genotype data into .tmap and .ped formats for further analysis. After completing the file format conversion in the above step, use the EMMAX-kin-intel64 program to calculate and generate the kinship matrix between individuals. Use the add covariate option -c and phenotypic data option -p to accompany the EMMAX-intel64 program to generate a .ps file and obtain the *P* value of each SINE-RIP site, and define the significance threshold as *P* < 1 × 10^-5. After we obtained the significant SINE-RIPs for the four phenotypes analyzed in GWAS, we used the compiled Python script to extract the genotype information of SINE-RIP in different populations, calculated the percentage of the two alleles and the percentage of the three genotypes of SINE-RIP in 362 individuals, and used the Python scipy.stats algorithm to perform a Chi-square test on the percentage of the alleles of the significant SINE-RIPs.

### Data collection for QTL

We retrieved pig QTL marker data from the Animal QTL Database (animalQTLdb, release 54; https://www.animalgenome.org/cgi-bin/QTLdb/SS/index) (*80*). The dataset comprised 50,098 QTL-SNPs (4-base span markers) associated with five major trait categories: Meat and Carcass (31,890 QTL-SNPs), Health (6,154), Exterior (2,185), Production (4,328), and Reproduction (5,541). Among the Production traits, body size-related QTL-SNPs were specifically distributed across body length (n=628), body weight (n=319), body height (n=173), and body circumference (n=328). To identify potential associations between common SINE-RIPs and known QTLs, we performed intersection analysis using BEDTOOLS (v2.30.0) with the -intersect option. For SINE-RIPs coinciding with body size-related QTL regions, we conducted frequency analysis across 362 *Sus scrofa* individuals, calculating both allele frequencies and genotype distributions. Statistical significance was assessed using Chi-square tests (implemented in scipy.stats, v1.7.0) to evaluate the distribution patterns of these SINE-RIPs genotype, following the same analytical approach used for GWAS-significant variants.

### Construction of SINE-RIPs online database

#### SINE-RIPs collection

Data for polymorphic SINE loci were collected through two approaches: (I.) mining by TypeSINE from short reads generated by next-generation sequencing technologies, and (II.) extraction of 36,284 SINE-RIPs using a genome-wide SINE-RIP mining protocol applied to 18 assembled genomes (*20*). The datasets were merged and processed to remove redundancy. Precise locations of each RIP were mapped according to the current pig genome reference sequences (UCSC Genome Browser assembly ID: susScr11; Sequencing/Assembly provider ID: The Swine Genome Sequencing Consortium (SGSC) Sscrofa11.1).

#### Database Design

The pigRIPdb Database was constructed using a combination of HTML, JavaScript, and PHP for the interactive interface development, while MySQL (version 5.7.44) was implemented for data management and sharing. The database architecture was designed with multiple functional modules including RIP overview, browser, search, BLAST, download, tools, links, and help sections. Logical relationships were optimized and appropriate API mappings were created to ensure seamless integration between all modules. The database was deployed on a UIS-B390-G2 server machine (H3C, China) running the Anolis 8.8 operating system.

## Acknowledgments

We thank Viktor Stéger (Institute of Genetics and Biotechnology, Hungarian University of Agriculture and Life Sciences), Li Zhu (Sichuan Agricultural University), TianFang Xiao (Fujian Agriculture and Forestry University), YongGang Liu (Yunnan Agricultural University) for providing valuable DNA samples.

## Author contributions

Conceptualization: C.S, N.Y and Y.Z. Methodology: Y.Z., N.Y and C.C. Resources: C.S and X.W. Investigation: C.S., Y.Z. and N.Y. Experimental verification: Y.Z., H.C. and S.S. Data analysis: Y.Z. and N.Y. Software Development: N.Y. and Y.Z. Visualization: Y.Z. Technical Consulting: K.W., E.M., H.R., W.Z. L.Z. and B.G. Funding acquisition: C.S. Supervision: C.S., X.W. and C.C. Writing—original draft: C.S., Y.Z., N.Y. and C.C. Writing—review & editing:

## Data and materials availability

All data are available in the main text and/or the Supplementary Materials.

## Supplementary Materials

Figs. S1 to S7

Tables S1 to S10

Data S1

## Other Supplementary Materials for this manuscript include the following

Data S2 to S9

