## Supplementary figure and table for "TypeSINE: Genome-Wide Detection of SINE Retrotransposon Polymorphisms Reveals Functional Variants Linked to Body Size Variation in Pigs"

Science Advances
AAAS

### Supplementary Materials for

#### Large-scale mining of SINE retrotransposon insertion polymorphisms in pig genomes identified potentially functional genomic regions linked to body size.

Yao Zheng *et al.*

**This PDF file includes:**

Figs. S1 to S8
Tables S1 to S11
Data S1

**Other Supplementary Materials for this manuscript include the following:**

Data S2 to S9

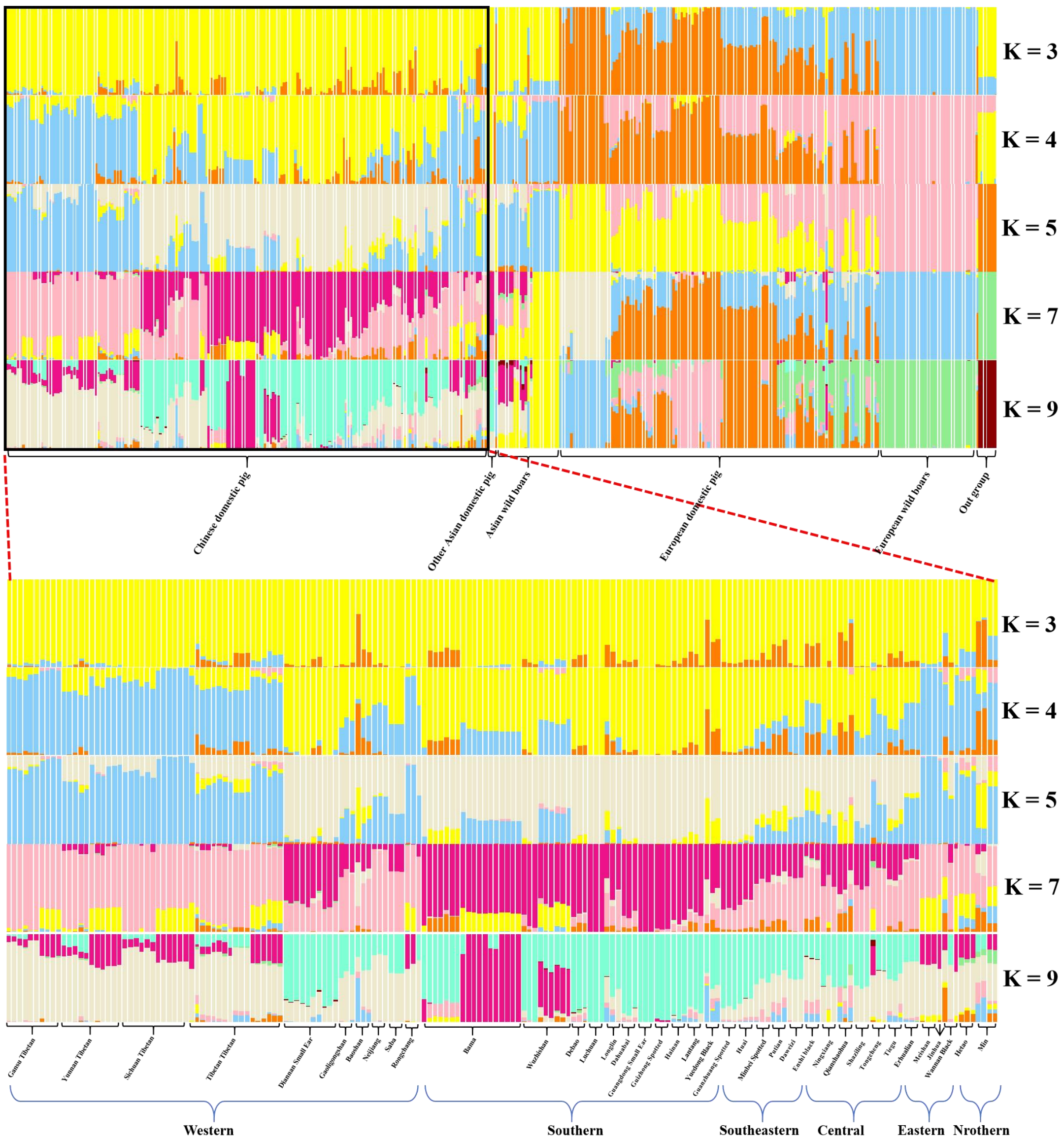


**Fig. S1.**

The genetic clustering analysis of Chinese domestic pigs

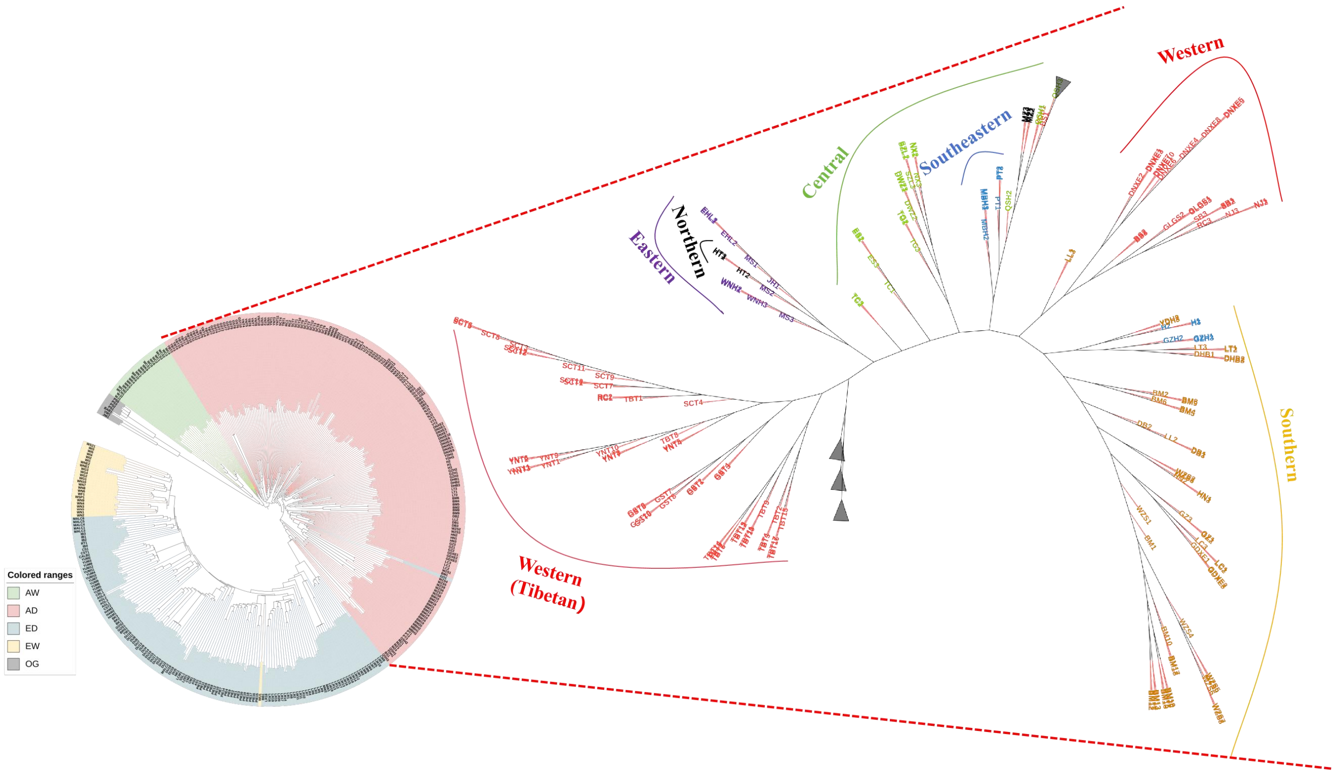


**Fig. S2.**

The phylogenetic tree of Chinese domestic pigs. In the right half of the phylogenetic tree, individuals in red font represent domestic pig populations from Western China, individuals in purple font represent domestic pig populations from Eastern China, individuals in black font represent domestic pig populations from Northern China, individuals in green font represent domestic pig populations from Central China, individuals in blue font represent domestic pig populations from Southern China, and individuals in yellow font represent domestic pig populations from Southeastern China.

**A** SINE-RIPs significantly associated with body weight by GWAS analysis

| SINE-RIP | Location | Insertion type/Orientation | Anotation | $-\log_{10}(P)$ | Chi-square test *P* value [a] | Small inviduals:80 breeds:9 | Small-Medium inviduals:59 breeds:13 | Medium inviduals:62 breeds:23 | Large-Medium inviduals:11 breeds:5 | Large inviduals:94 breeds:13 |
|---|---|---|---|---|---|---|---|---|---|---|
| SRIP-1088 | chr1:114659197-114659493 | deletion/+ | / | 5.42 | 5.31E-40*** | | | | | |
| SRIP-29571 | chr1:271114588-271114589 | insertion/+ | *NUP214* [b] | 5.63 | 1.81E-22*** | | | | | |
| SRIP-17676 | chr4:75648999-75649292 | deletion/- | *PLAG1* [b] | 9.51 | 9.30E-63*** | | | | | |
| SRIP-17680 | chr4:75874803-75875073 | deletion/+ | *LYN* [b] | 9.82 | 1.92E-60*** | | | | | |
| SRIP-55579 | chr5:66073491-66073492 | insertion/+ | / | 6.19 | 0.003152* | | | | | |
| SRIP-55780 | chr5:77025651-77025652 | insertion/- | *SLC38A1* [b] | 5.53 | 1.54E-28*** | | | | | |
| SRIP-58040 | chr6:102395985-102395986 | insertion/+ | / | 5.38 | 0.000575** | | | | | |
| SRIP-25527 | chr9:107520300-107520585 | deletion/- | / | 5.49 | 5.55E-13*** | | | | | |
| SRIP-30333 | chr10:31404571-31404572 | insertion/+ | *FRMD3* [b] | 5.35 | 1.06E-26*** | | | | | |
| SRIP-30900 | chr10:56399164-56399165 | insertion/- | *NRP1* [b] | 6.39 | 1.97E-08*** | | | | | |
| SRIP-9378 | chr15:104549657-104549951 | deletion/- | / | 5.68 | 9.48E-31*** | | | | | |
| SRIP-9383 | chr15:104894664-104894956 | deletion/+ | *CASP10* [b] | 6.23 | 1.55E-31*** | | | | | |
| SRIP-40531 | chr15:105764199-105764200 | insertion/- | *KIAA2012* [b] | 5.51 | 1.07E-13*** | | | | | |
| SRIP-42782 | chr17:10795636-10795637 | insertion/+ | *ANK1* [b] | 6.08 | 0.000034*** | | | | | |
| SRIP-42799 | chr17:11271260-11271261 | insertion/- | / | 5.41 | 0.001265* | | | | | |

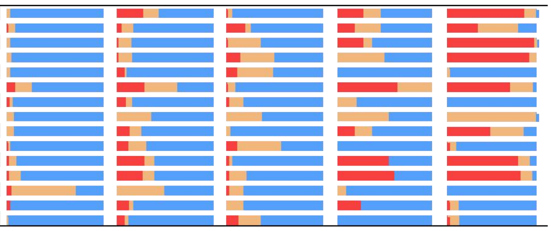

**B** Body Weight of breeds information

| Categories | Range/kg | Total number of breeds | Total number of individuals | Breeds (individual number) |
|---|---|---|---|---|
| Small | < 70 | 9 | 80 | creole (1) Göttingen minipig (1)Bama (18) Diannan Small Ear (10) Gansu Tibetan (10) Gaoligongshan (3) Tibetan Tibetan (17) Wuzhishan (9) Yunnan Tibetan (11) |
| Small-Medium | 70 - 100 | 13 | 59 | Mangalica (6) Yucatan miniature pig (12) Debao (3) Guanzhuang Spotted (3) Guizhong Spotted (3) Hainan (3) Huai (3) Jinhua (1) Lantang (3) Luchuan (3) Min (4) Minbei Spotted (3) Sichuan Tibetan (12) |
| Medium | 100 - 150 | 23 | 62 | Calabrese (1) iberian (4) Leicoma (1) Nera_Siciliana (1) Retinto (1) Baoshan (3) Dahuabai (3) Daweizi (3) Enshi black (3) Erhualian (3) Guangdong Small Ear (3) Hetao (3) Longlin (3) Meishan (3) Ningxiang (3) Putian (3) Qianshaohua (3) Saba (3) Shaziling (3) Tiegu (3) Tongcheng (3) Wannan Black (3) Yuedong Black (3) |
| Large-Medium | 150 - 200 | 5 | 11 | Casertana (2) Chato_Murciano (2) Cinta_Senese (1) Neijiang (3) Rongchang (3) |
| Large | > 200 | 13 | 94 | Angler_Sattleschwein (2) British_Saddleback (1) Bunte_Bentheimer (1) Gloucester_Old_Spot (2) Linderodsvin (1) Middle_White (2) Tamworth (1) Berkshire (3) Duroc (19) Hampshire (3) Landrace (20) Pietrain (20) Yorkshire (19) |

#### Fig. S3.

Genomic annotation for SINE-RIPs significantly associated with body weight by GWAS analysis (A) and breed information for different mature body weight categories (B). The bars represent the percentage of genotypes, where red represents "+/+", yellow represents "+/-", and blue represents "-/-". [a] Significant differences of genotype frequencies between different groups were tested by Chi-square test, and labeled as ** P< 0.05, *** P< 0.001, **** P< 0.0001. [b] represents SINE-RIP inserted in protein coding genes.

#### A SINE-RIPs significantly associated with body length by GWAS analysis

| SINE-RIP | Location | Insertion type/Orientation | Anotation | $-\log_{10}(P)$ | Chi-square test $P$ value [a] | Short invividuals:102 breeds:11 | Medium invividuals:81 breeds:26 | Long invividuals:107 breeds:15 |
|---|---|---|---|---|---|---|---|---|
| SRIP-27823 | chr1:139293952-139293953 | insertion/- | / | 5.41 | 0.006534* | | | |
| SRIP-17676 | chr4:75648999-75649292 | insertion/- | *PLAG1* [b] | 7.97 | 5.16E-35*** | | | |
| SRIP-17680 | chr4:75874803-75875073 | deletion/+ | *LYN* [b] | 5.41 | 1.47E-32*** | | | |
| SRIP-55534 | chr5:64580231-64580232 | insertion/- | *VWF* [b] | 5.04 | 5.54E-13*** | | | |
| SRIP-62486 | chr8:79927507-79927508 | insertion/- | / | 5.44 | 0.168155 | | | |
| SRIP-30333 | chr10:31404571-31404572 | deletion/+ | *FRMD3* [b] | 6.71 | 4.03E-16*** | | | |
| SRIP-34990 | chr13:68339577-68339580 | deletion/+ | *PPARG* [b] | 5.85 | 3.80E-10*** | | | |

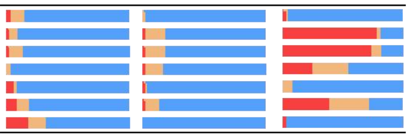

#### B Body Length of breeds information

| Categories | Range/cm | Total number of breeds | Total number of individuals | Breeds (individual number) |
|---|---|---|---|---|
| Short | < 100 | 11 | 102 | Angler_Sattleschwein (2) Bama (18) Debao (3) Diannan Small Ear (10) Gansu Tibetan (10) iberian (4) Mangalica (6) Tibetan Tibetan (17) Wuzhishan (9) Yunnan Tibetan (11) Yucatan miniature pig (12) |
| Medium | 100 - 140 | 26 | 81 | Baoshan (3) Cinta_Senese (1) Casertana (2) Dahuabai (3) Daweizi (3) Erhualian (3) Enshi black (3) Guangdong Small Ear (3) Gaoligongshan (3) Guizhong Spotted (3) Guanzhuang Spotted (3) Huai (3) Hainan (3) Jinhua (1) Luchuan (3) Longlin (3) Lantang (3) Minbei Spotted (3) Meishan (3) Min (4) Putian (3) Qianshaohua (3) Sichuan Tibetan (12) Nera_Siciliana (1) Shaziling (3) Yuedong Black (3) |
| Long | > 140 | 15 | 107 | Berkshire (3) Calabrese (1) Duroc (19) Hampshire (3) Hetao (3) Landrace (20) Yorkshire (19) Neijiang (3) Pietrain (20) Rongchang (3) Saba (3) Tongcheng (3) Tiegu (3) Tamworth (1) Wannan Black (3) |

**Fig. S4.**

Genomic annotation for SINE-RIPs significantly associated with body length by GWAS analysis (A) and breed information for different mature body length categories (B). The bars represent the percentage of genotypes, where red represents "+/+", yellow represents "+/-", and blue represents "-/-". a Significant differences of genotype frequencies between different groups were tested by Chi-square test, and labeled as ** P< 0.05, *** P< 0.001, **** P< 0.0001. b represents SINE-RIP inserted in protein coding genes.

**A** **SINE-RIPs significantly associated with body height by GWAS analysis**

| SINE-RIP | Location | Insertion type/Orientation | Anotation | -log10(*P*) | Chi-square test *P* value | Short invidual:29 breeds:3 | Medium invidual:163 breeds:39 | Tall invidual:97 breeds:12 |
|---|---|---|---|---|---|---|---|---|
| SRIP-994 | chr1:104712337-104712627 | deletion/+ | / | 6.65 | 5.78E-34*** | | | |
| SRIP-1603 | chr1:178975470-178975757 | deletion/+ | / | 7.36 | 1.32E-41*** | | | |
| SRIP-2141 | chr1:246455668-246455961 | insertion/- | / | 5.15 | 1.49E-54*** | | | |
| SRIP-16033 | chr3:59877939-59878206 | deletion/+ | *DNAH6* | 5.19 | 1.33E-25*** | | | |
| SRIP-52512 | chr4:21568066-21568067 | insertion/- | / | 5.05 | 1.09E-46*** | | | |
| SRIP-54596 | chr5:7568092-7568093 | deletion/+ | *XPNPEP3* | 6.27 | 2.03E-17*** | | | |
| SRIP-55904 | chr5:83115089-83115090 | deletion/+ | *ANO4* | 5.12 | 7.56E-33*** | | | |
| SRIP-60073 | chr7:43588729-43588730 | insertion/- | / | 5.61 | 0.038438* | | | |
| SRIP-60264 | chr7:55546210-55546211 | deletion/+ | / | 5.51 | 2.82E-11*** | | | |
| SRIP-23621 | chr8:46573773-46574066 | deletion/+ | / | 5.18 | 3.56E-37*** | | | |
| SRIP-36689 | chr13:204479511-204479512 | insertion/- | / | 5.39 | 5.17E-31*** | | | |
| SRIP-40382 | chr15:95048079-95048080 | insertion/- | *MFSD6* | 5.33 | 4.22E-35*** | | | |
| SRIP-40597 | chr15:109811627-109811628 | deletion/+ | *ADAM23* | 5.92 | 0.048431* | | | |
| SRIP-9659 | chr15:130612553-130612813 | deletion/+ | *DNER* | 5.21 | 1.20E-34*** | | | |
| SRIP-42799 | chr17:11271260-11271261 | insertion/- | / | 5.74 | 0.000021*** | | | |
| SRIP-43803 | chr17:60055863-60055864 | deletion/+ | / | 5.52 | 5.00E-07*** | | | |
| SRIP-43878 | chr18:2229234-2229235 | deletion/+ | / | 5.47 | 5.57E-35*** | | | |

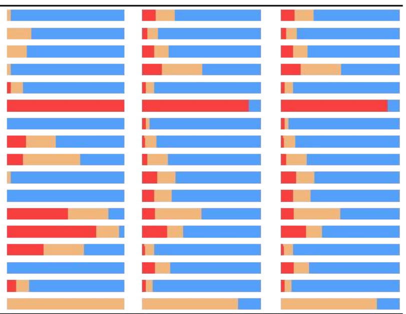

**B** **Body Height of breeds information**

| Categories | Range/cm | Total number of breeds | Total number of individuals | Breeds (individual number) |
|---|---|---|---|---|
| Short | < 50 | 3 | 29 | Bama (18) Göttingen minipig (1) Gansu Tibetan (10) |
| Medium | 50 - 80 | 39 | 163 | Bunte_Bentheimer (1) Baoshan (3) Debao (3) Dahuabai (3) Diannan Small Ear (10) Daweizi (3) Erhualian (3) Enshi black (3) Guangdong Small Ear (3) Gaoligongshan (3) Guizhong Spotted (3) Guanzhuang Spotted (3) Huai (3) Hainan (3) Jinhua (1) Luchuan (3) Linderodsvin (1) Longlin (3) Lantang (3) Mangalica (6) Minbei Spotted (3) Meishan (3) Neijiang (3) Putian (3) Qianshaohua (3) Rongchang (3) Saba (3) Sichuan Tibetan (12) Nera_Siciliana (1) Shaziling (3) Tibetan Tibetan (17) Tongcheng (3) Tiegu (3) Tamworth (1) Wannan Black (3) Wuzhishan (9) Yuedong Black (3) Yunnan Tibetan (11) Yucatan miniature pig (12) |
| Tall | > 80 | 12 | 97 | Angler_Sattleschwein (2) Berkshire (3) Calabrese (1) Chato_Murciano (2) Cinta_Senese (1) Duroc (19) Hampshire (3) Hetao (3) Landrace (20) Yorkshire (19) Min (4) Pietrain (20) |

**Fig. S5.**

Genomic annotation for SINE-RIPs significantly associated with body height by GWAS analysis (A) and breed information for different mature body height categories (B). The bars represent the percentage of genotypes, where red represents "+/+", yellow represents "+/-", and blue represents "-/-". a Significant differences of genotype frequencies between different groups were tested by Chi-square test, and labeled as ** P< 0.05, *** P< 0.001, **** P< 0.0001. b represents SINE-RIP inserted in protein coding genes.

#### A SINE-RIPs significantly associated with body circumference by GWAS analysis

| SINE-RIP | Location | Insertion type/Orientation | Anotation | -log10(*P*) | Chi-square test *P* value [a] | Short invidusals:78 breeds:7 | Medium invidusals:72 breeds:21 | Long invidusals:119 breeds:19 |
|---|---|---|---|---|---|---|---|---|
| SRIP-48777 | chr2:131051894-131051895 | insertion/- | *SLC12A2* [b] | 5.21 | 2.00E-12*** | | | |
| SRIP-53735 | chr4:110993604-110993605 | insertion/- | / | 5.60 | 0.004447* | | | |
| SRIP-55828 | chr5:79361134-79361135 | insertion/- | *APPL2* [b] | 6.44 | 7.04E-07*** | | | |
| SRIP-58209 | chr6:115923318-115923319 | insertion/- | / | 5.07 | 2.75E-09*** | | | |
| SRIP-40807 | chr15:121407843-121407844 | deletion/+ | *DNPEP* [b] | 5.36 | 0.000008*** | | | |
| SRIP-11543 | chr18:14759288-14759575 | deletion/+ | / | 5.69 | 0.000007*** | | | |

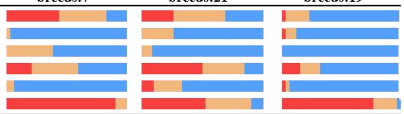

#### B Body Circumference of breeds information

| Categories | Range/cm | Total number of breeds | Total number of individuals | Breeds (individual number) |
|---|---|---|---|---|
| Short | < 100 | 7 | 78 | Bama (18) Diannan Small Ear (10) Gaoligongshan (3) Gansu Tibetan (10) Tibetan Tibetan (17) Wuzhishan (9) Yunnan Tibetan (11) |
| Medium | 100 - 120 | 21 | 72 | Baoshan (3) Debao (3) Dahuabai (3) Daweizi (3) Enshi black (3) Guangdong Small Ear (3) Guizhong Spotted (3) Guanzhuang Spotted (3) Huai (3) Hainan (3) Luchuan (3) Longlin (3) Lantang (3) Minbei Spotted (3) Meishan (3) Putian (3) Qianshaohua (3) Rongchang (3) Saba (3) Sichuan Tibetan (12) Shaziling (3) |
| Long | >120 | 19 | 119 | Berkshire (3) Calabrese (1) Cinta_Senese (1) Duroc (19) Erhualian (3) Hampshire (3) Hetao (3) Jinhua (1) Landrace (20) Yorkshire (19) Mangalica (6) Min (4) Neijiang (3) Pietrain (20) Nera_Siciliana (1) Tongcheng (3) Tiegu (3) Wannan Black (3) Yuedong Black (3) |

**Fig. S6.**

Genomic annotation for SINE-RIPs significantly associated with body circumference by GWAS analysis (A) and breed information for different mature body circumference categories (B). The bars represent the percentage of genotypes, where red represents "+/+", yellow represents "+/-", and blue represents "-/-". a Significant differences of genotype frequencies between different groups were tested by Chi-square test, and labeled as ** P< 0.05, *** P< 0.001, **** P< 0.0001. b represents SINE-RIP inserted in protein coding genes.

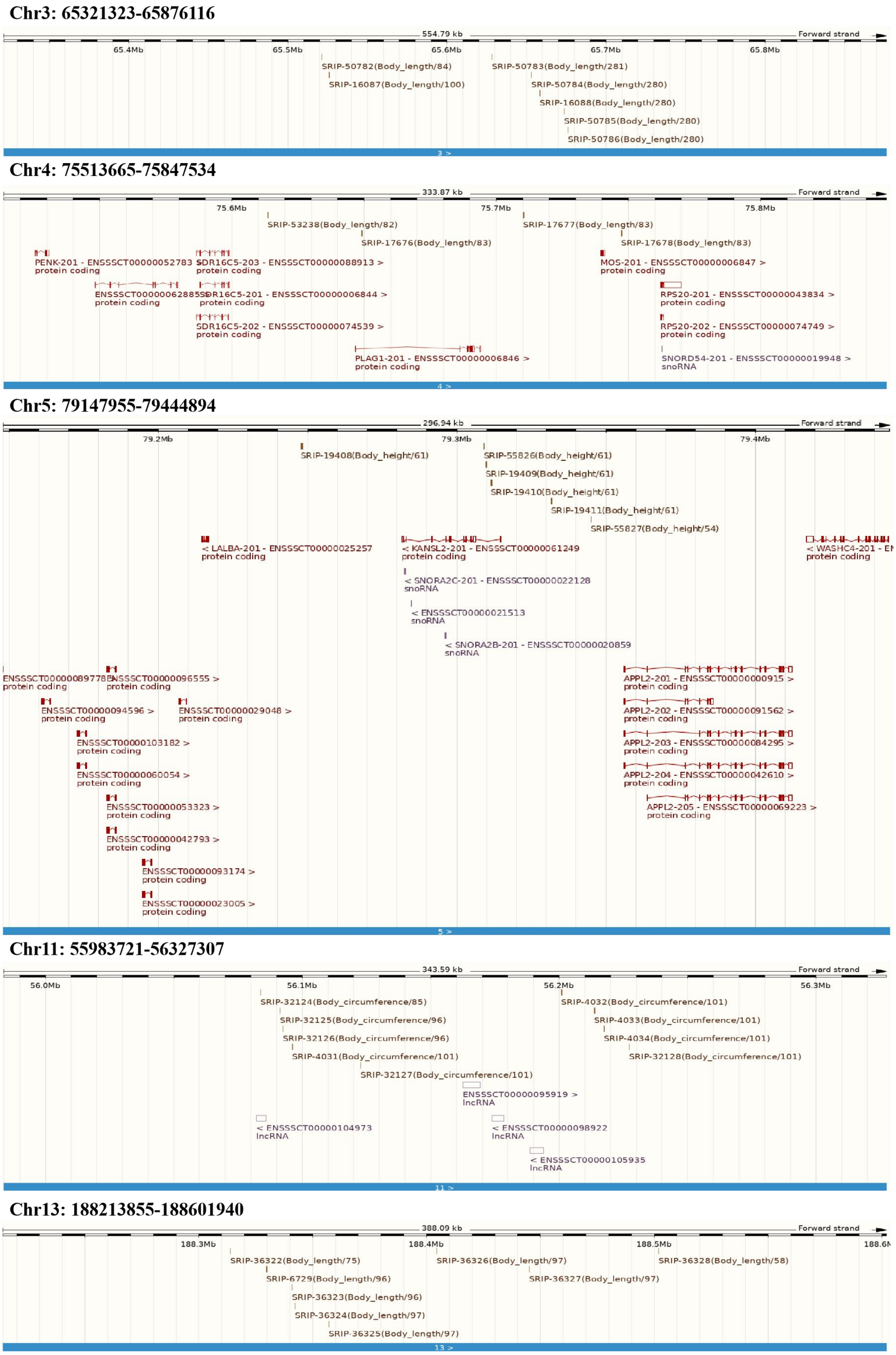


**Fig. S7. Annotation for five genomic regions, containing 34 SINE-RIPs enriched QTL-SNPs of body size (body length, body height, and body circumference).**

#### Annotation of 34 SINE-RIPs enriching QTL-SNPs of body size

| Genomic region of Growth traits association | SINE-RIP | Location | Insertion type /Orientation | Number of overlap QTLs | Gene anotation | Chi-square test *P* value [a] |
|---|---|---|---|---|---|---|
| Chromosome 3<br>Genomic region: 65321323-65876116<br>Spanning size: 554.79 kb<br>281 length QTL-SNPs | SRIP-50782 | 65521323-65521324 | ins / - | 84 | / | 0.03018* |
| | SRIP-16087 | 65525944-65526238 | del / - | 100 | / | 0.0002** |
| | SRIP-50783 | 65628283-65628284 | ins / + | 281 | / | 1.15E-09*** |
| | SRIP-50784 | 65652980-65652981 | ins / - | 280 | / | 0.105986 |
| | SRIP-16088 | 65658298-65658596 | del / + | 280 | / | 0.060804 |
| | SRIP-50785 | 65673667-65673668 | ins / + | 280 | / | 0.004759* |
| | SRIP-50786 | 65676115-65676116 | ins / + | 280 | / | 0.005608* |
| Chromosome 4<br>Genomic region: 75513665-75847534<br>Spanning size:333.87 kb<br>83 Body length QTL-SNPs) | SRIP-53238 | 75613665-75613666 | ins / + | 82 | / | 0.005132* |
| | SRIP-17676 | 75648999-75649292 | del / - | 83 | *PLAG1* [b] | 5.16E-35*** |
| | SRIP-17677 | 75710213-75710512 | del / + | 83 | / | 1 |
| | SRIP-17678 | 75747356-75747534 | del / - | 83 | / | 0.143955 |
| Chromosome13<br>Gebomic region: 188213855-188601940<br>Spanning size: 388.09 kb<br>97 Body length QTLs-SNP | SRIP-36322 | 188313855-188313856 | ins / + | 75 | / | 0.000295** |
| | SRIP-6729 | 188329605-188329912 | del / - | 96 | / | 0.000105** |
| | SRIP-36323 | 188340883-188340884 | ins / + | 96 | / | 0.050224 |
| | SRIP-36324 | 188342324-188342325 | ins / - | 97 | / | 0.000671** |
| | SRIP-36325 | 188357041-188357042 | ins / - | 97 | / | 3.91E-11*** |
| | SRIP-36326 | 188404445-188404446 | ins / + | 97 | / | 0.032532* |
| | SRIP-36327 | 188445159-188445160 | ins / + | 97 | / | 0.000027*** |
| | SRIP-36328 | 188501939-188501940 | ins / + | 58 | / | 2.47E-09*** |
| Chromosome 5<br>Genomic region: 79147955-79444894<br>Spanning size: 296.94 kb<br>61 Body height QTL-SNPs | SRIP-19408 | 79247955-79248261 | del / - | 61 | / | 7.61E-24*** |
| | SRIP-55826 | 79309284-79309285 | ins / + | 61 | *KANSL2* [b] | 0.004482* |
| | SRIP-19409 | 79309789-79310083 | del / - | 61 | *KANSL2* [b] | 0.007458* |
| | SRIP-19410 | 79311620-79311894 | del / - | 61 | *KANSL2* [b] | 0.96821 |
| | SRIP-19411 | 79331721-79331985 | del / + | 61 | / | 0.000188** |
| | SRIP-55827 | 79344893-79344894 | ins / + | 54 | / | 1.47E-09*** |
| Chromosome 11<br>Genomic region: 55983721-56327307<br>Spanning size: 343.59 kb<br>101 Body circumference QTL-SNPs | SRIP-32124 | 56083721-56083722 | ins / + | 85 | *ENSSSCG00000061191* [c] | 0.479065 |
| | SRIP-32125 | 56091281-56091282 | ins / + | 96 | / | 0.056519 |
| | SRIP-32126 | 56092468-56092469 | ins / + | 96 | / | 0.61376 |
| | SRIP-4031 | 56096010-56096313 | del / + | 101 | / | 3.07E-09*** |
| | SRIP-32127 | 56122737-56122738 | ins / + | 101 | / | 0.003089* |
| | SRIP-4032 | 56200825-56201124 | del / + | 101 | / | 0.000098*** |
| | SRIP-4033 | 56213693-56213987 | del / - | 101 | / | 0.000001*** |
| | SRIP-4034 | 56217358-56217622 | del / + | 101 | / | 3.36E-19*** |
| | SRIP-32128 | 56227306-56227307 | ins / + | 101 | / | 3.71E-12*** |

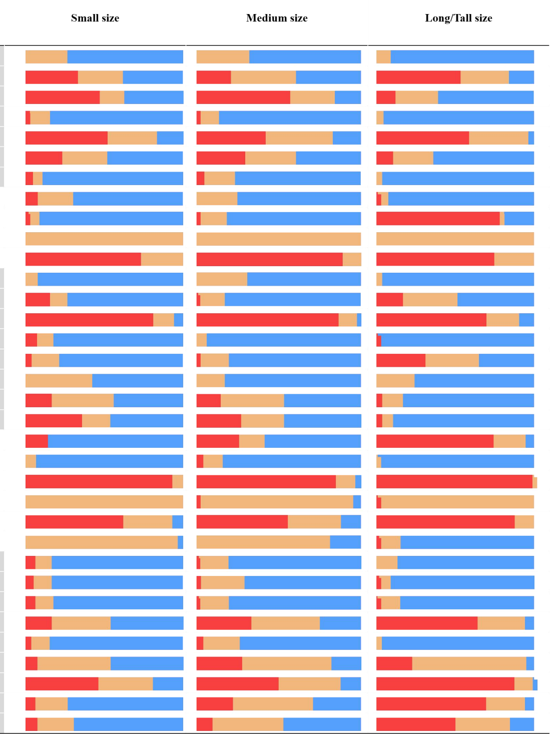


**Fig. S8.**

Genomic annotation for 34 SINE-RIPs enriched QTL-SNPs of body size (body weight, body length, body height and body circumference). The bars represent the percentage of genotypes, where red represents "+/+", yellow represents "+/-", and blue represents "-/-". a Significant differences of genotype frequencies between different groups were detected by Chi-square test, and labeled as ** P< 0.05, *** P< 0.001, **** P< 0.0001. b represents SINE-RIP inserted in protein coding genes. c. represents SINE-RIP inserted in lncRNA gene.

**Table S1.**

The SINEA1-3 consensus sequences

| SINE subfamilies | consensus sequences |
|---|---|
| SINEA1 | GGAGTTCCCGTCGTGGCGCAGTGGTTAACGAATCCGACTAGGAACCATGAGGTTGCGGGTTC<br>GGTCCCTGCCCTTGCTCAGTGGGTTAACGATCCGGCGTTGCCGTGAGCTGTGGTGTAGGTTGC<br>AGACGCGGCTCGGATCCCGCGTTGCTGTGGCTCTGGCGTAGGCCGGTGGCTACAGCTCCGATT<br>CGACCCCTAGCCTGGGAACCTCCATATGCCGCGGGAGCGGCCCAAGAAATAGCAACAACAAC<br>AACAACAAAAAAGACAAAAAGACCAAAAAAAAAAAAAAAAAAAA |
| SINEA2 | GGAGTTCCCGTCGTGGCGCAGTGGTTAACGAATCCGACTAGGAACCATGAGGTTGCGGGTTC<br>GGTCCCTGCCCTTGCTCAGTGGGTTAACGATCCGGCGTTGCCGTGAGCTGTGGTGTAGGTTGC<br>AGACGCGGCTCGGATCCCGCGTTGCTGTGGCTCTGGCGTAGGCCGGCGGCTACAGCTCCGATT<br>CGACCCCTAGCCTGGGAACCTCCATATGCCGCGGGAGCGGCCCAAGAAATAGCAAAAAGACA<br>AAAAAAAAAAAAA |
| SINEA3 | GGAGTTCCCGTCGTGGCGCAGTGGTTAACGAATCCGACTAGGAACCATGAGGTTGCGGGTTC<br>GGTCCCTGCCCTTGCTCAGTGGGTTAACGATCCGGCGTTGCCGTGAGCTGTGGTGTAGGTTGC<br>AGACGCGGCTCGGATCCCGCGTTGCTGTGGCTCTGGCGTAGGCCGGTGGCTACAGCTCCGATT<br>CGACCCCTAGCCTGGGAACCTCCATATGCCGCGGGAGCGGCCCAAGAAATAGCAAAAAGACA<br>AAAAAAAAAA |

**Table S2.**

Test results of TypeSINE and MELT

| | Number of test samples | Average depth | Total time (minutes) | Number of SINE-RIP | MAF≥0.05 | MAF<0.05 |
|---|---|---|---|---|---|---|
| TypeSINE | 50 | 37X | 99597 | 191218 | 53932 | 137286 |
| MELT | | | 93106 | 73412 | 41930 | 31482 |

**Table S3.**

SINE insertion (SINE+ ) allele frequencies detected based on the mined RIPs by using the sequenced genomes of domesticated pigs (314 biosamples) and wild pigs (48 biosamples)

| RIP_ND* (%) | Number of samples | SINE insertion (SINE+ ) allele frequencies (IAF) | | | | | | | total |
|---|---|---|---|---|---|---|---|---|---|
| | | 0-0.05 | 0.05-0.1 | 0.1-0.2 | 0.2-0.8 | 0.8-0.9 | 0.9-0.95 | 0.95-1 | |
| 25 | 91 | 520769 | 12472 | 10367 | 29763 | 3892 | 2683 | 75909 | 655855 |
| 50 | 182 | 567985 | 13628 | 11167 | 31135 | 4079 | 2829 | 78148 | 708971 |
| 75 | 272 | 604931 | 14897 | 11913 | 32048 | 4169 | 2890 | 78987 | 749835 |
| 100 | 362 | 647381 | 25223 | 19767 | 38990 | 4281 | 2973 | 79143 | 817758 |

* RIP_ND(%): The maximum ratio of samples with undetected genotypes.

**Table S4.**

Summary of samples information

| | abbreviation | Breed name | Number of biosamples | Number of SRAs |
|---|---|---|---|---|
| Domestic pig | BM | Bama | 18 | 21 |
| | BS | Baoshan pig | 3 | 3 |
| | DHB | Dahuabai pig | 3 | 3 |
| | DWZ | Daweizi | 3 | 3 |
| | DB | Debao pig | 3 | 3 |
| | DNXE | Diannan Small Ear pig | 10 | 10 |
| | ES | Enshi black | 3 | 3 |
| | EHL | Erhualian | 3 | 3 |
| | GST | Gansu Tibetan | 10 | 14 |
| | GLGS | Gaoligongshan pig | 3 | 3 |
| | GDXE | Guangdong Small Ear pig | 3 | 3 |
| | GZH | Guanzhuang Spotted pig | 3 | 3 |
| | GZ | Guizhong Spotted pig | 3 | 3 |
| | HN | Hainan pig | 3 | 3 |
| | HT | Hetao | 3 | 6 |
| | H | Huai pig | 3 | 3 |
| | JH | Jinhua | 1 | 5 |
| | LT | Lantang | 3 | 3 |
| | LL | Longlin pig | 3 | 3 |
| | LC | Luchuan | 3 | 3 |
| | MS | Meishan | 3 | 6 |
| | MZ | Min | 4 | 4 |
| | MBH | Minbei Spotted pig | 3 | 3 |
| | NJ | Neijiang pig | 3 | 3 |
| | NX | Ningxiang | 3 | 3 |
| | PT | Putian pig | 3 | 3 |
| | QSH | Qinshaohua | 3 | 3 |
| | RC | Rongchang | 3 | 6 |
| | SB | Saba | 3 | 3 |
| | SZL | shaziling | 3 | 3 |
| | SCT | Sichuan Tibetan | 12 | 18 |
| | TBT | Tibetan Tibetan | 17 | 29 |
| | TG | Tiegu | 3 | 3 |
| | TC | Tongcheng | 3 | 4 |
| | WNH | Wannan Black pig | 3 | 3 |
| | WZS | Wuzhishan | 9 | 15 |

| | | | | |
|---|---|---|---|---|
| | YDH | Yuedong Black pig | 3 | 3 |
| | YNT | Yunnan Tibetan | 11 | 17 |
| | AS | Angler_Sattleschwein | 2 | 6 |
| | AGL | Angola | 4 | 4 |
| | BRS | British_Saddleback | 1 | 2 |
| | BB | Bunte_Bentheimer | 1 | 1 |
| | CALB | Calabrese | 1 | 4 |
| | CST | Casertana | 2 | 6 |
| | CM | Chato_Murciano | 2 | 5 |
| | CS | Cinta_Senese | 1 | 3 |
| | CRL | creole | 1 | 1 |
| | GOS | Gloucester_Old_Spot | 2 | 2 |
| | IB | iberian | 4 | 6 |
| | LCM | Leicoma | 1 | 2 |
| | LDD | Linderodsvin | 1 | 2 |
| | MALC | Mangalica | 6 | 10 |
| | MW | Middle_White | 2 | 4 |
| | NK | Nanchukmacdon | 1 | 1 |
| | SSL | Nera_Siciliana | 1 | 4 |
| | RT | Retinto | 1 | 4 |
| | TW | Tamworth | 1 | 3 |
| | VT | Vietnamese native pig | 3 | 3 |
| | GO | Göttingen minipig | 1 | 4 |
| | YUM | Yucatan miniature pig | 12 | 12 |
| | BK | Berkshire | 3 | 8 |
| | DU | Duroc | 19 | 24 |
| | HAM | Hampshire | 3 | 9 |
| | LR | Landrace | 20 | 33 |
| | PIE | Pietrain | 20 | 23 |
| | LW | Yorkshire | 19 | 20 |
| Wild boar | WC | Wild_China | 6 | 16 |
| | WCN | Wild_Northern China | 2 | 10 |
| | WCS | Wild_South China | 3 | 17 |
| | WF | Wild_Japan | 1 | 8 |
| | WG | Wild_Korea | 4 | 12 |
| | WI | Wild_France | 5 | 12 |
| | WJ | Wild_Germany | 2 | 9 |
| | WK | Wild_Italy | 10 | 29 |
| | WN | Wild_Netherlands | 10 | 26 |
| | WNE | Wild_Eastern Europe | 2 | 6 |
| | WSP | Wild_Spain | 1 | 1 |
| | WSZ | Wild_Switzerland | 1 | 2 |
| | WU | Wild_Ukraine | 1 | 2 |

| | | | | |
|---|---|---|---|---|
| Outgroup | BAB | Babyrousa babyrussa | 1 | 1 |
| | PS | Porcula salvania | 4 | 4 |
| | VW | Sus cebifrons | 1 | 1 |
| | JW | Sus verrucosus | 1 | 20 |

**Table S5.**

Significant SINE-RIPs for GWAS (body weight)

| SINE-RIP | chromosome | ID | p_score | -log10(P) |
|---|---|---|---|---|
| SRIP-1088 | 1 | pig_ctg1:114659197-114659493(+) | 3.82E-06 | 5.417501801 |
| SRIP-29571 | 1 | pig_ctg1:271114588-271114589(+) | 2.32E-06 | 5.634330952 |
| SRIP-17676 | 4 | pig_ctg4:75648999-75649292(-) | 3.08E-10 | 9.511281475 |
| SRIP-17680 | 4 | pig_ctg4:75874803-75875073(+) | 1.53E-10 | 9.81632315 |
| SRIP-55579 | 5 | pig_ctg5:66073491-66073492(+) | 6.45E-07 | 6.19019324 |
| SRIP-55780 | 5 | pig_ctg5:77025651-77025652(-) | 2.95E-06 | 5.529680672 |
| SRIP-58040 | 6 | pig_ctg6:102395985-102395986(+) | 4.13E-06 | 5.38450779 |
| SRIP-25527 | 9 | pig_ctg9:107520300-107520585(-) | 3.27E-06 | 5.485739507 |
| SRIP-30333 | 10 | pig_ctg10:31404571-31404572(+) | 4.46E-06 | 5.350564651 |
| SRIP-30900 | 10 | pig_ctg10:56399164-56399165(-) | 4.10E-07 | 6.387671554 |
| SRIP-9378 | 15 | pig_ctg15:104549657-104549951(-) | 2.08E-06 | 5.681444809 |
| SRIP-9383 | 15 | pig_ctg15:104894664-104894956(+) | 5.83E-07 | 6.234183987 |
| SRIP-40531 | 15 | pig_ctg15:105764199-105764200(-) | 3.10E-06 | 5.508630933 |
| SRIP-42782 | 17 | pig_ctg17:10795636-10795637(+) | 8.36E-07 | 6.077623093 |
| SRIP-42799 | 17 | pig_ctg17:11271260-11271261(-) | 3.87E-06 | 5.412708469 |

**Table S6.**

Significant SINE-RIPs for GWAS (body length)

| SINE-RIP | chromosome | ID | p_score | -log10(P) |
|---|---|---|---|---|
| SRIP-27823 | 1 | pig_ctg1:139293952-139293953(-) | 3.89E-06 | 5.410421804 |
| SRIP-17676 | 4 | pig_ctg4:75648999-75649292(-) | 1.07E-08 | 7.971827774 |
| SRIP-17680 | 4 | pig_ctg4:75874803-75875073(+) | 3.89E-06 | 5.410235616 |
| SRIP-55534 | 5 | pig_ctg5:64580231-64580232(-) | 9.15E-06 | 5.038499493 |
| SRIP-62486 | 8 | pig_ctg8:79927507-79927508(-) | 3.60E-06 | 5.444100434 |
| SRIP-30333 | 10 | pig_ctg10:31404571-31404572(+) | 1.97E-07 | 6.706421571 |
| SRIP-34990 | 13 | pig_ctg13:68339577-68339580(+) | 1.41E-06 | 5.850496313 |

**Table S7.**

Significant SINE-RIPs for GWAS (body height)

| SINE-RIP | chromosome | ID | p_score | -log10(P) |
|---|---|---|---|---|
| SRIP-994 | 1 | pig_ctg1:104712337-104712627(+) | 2.24E-07 | 6.65049364 |
| SRIP-1603 | 1 | pig_ctg1:178975470-178975757(+) | 4.41E-08 | 7.355835903 |
| SRIP-2141 | 1 | pig_ctg1:246455668-246455961(+) | 7.08E-06 | 5.150089961 |
| SRIP-16033 | 3 | pig_ctg3:59877939-59878206(-) | 6.52E-06 | 5.185988994 |
| SRIP-52512 | 4 | pig_ctg4:21568066-21568067(+) | 8.99E-06 | 5.046179967 |
| SRIP-54596 | 5 | pig_ctg5:7568092-7568093(+) | 5.32E-07 | 6.27404556 |
| SRIP-55904 | 5 | pig_ctg5:83115089-83115090(-) | 7.64E-06 | 5.11665658 |
| SRIP-60073 | 7 | pig_ctg7:43588729-43588730(+) | 2.46E-06 | 5.609486558 |
| SRIP-60264 | 7 | pig_ctg7:55546210-55546211(+) | 3.08E-06 | 5.511283892 |
| SRIP-23621 | 8 | pig_ctg8:46573773-46574066(+) | 6.65E-06 | 5.177101125 |
| SRIP-36689 | 13 | pig_ctg13:204479511-204479512(-) | 4.03E-06 | 5.394497976 |
| SRIP-40382 | 15 | pig_ctg15:95048079-95048080(-) | 4.64E-06 | 5.333074153 |
| SRIP-40597 | 15 | pig_ctg15:109811627-109811628(+) | 1.21E-06 | 5.917086615 |
| SRIP-9659 | 15 | pig_ctg15:130612553-130612813(+) | 6.22E-06 | 5.206508546 |
| SRIP-42799 | 17 | pig_ctg17:11271260-11271261(-) | 1.82E-06 | 5.739111719 |
| SRIP-43803 | 17 | pig_ctg17:60055863-60055864(-) | 3.05E-06 | 5.515799462 |
| SRIP-43878 | 18 | pig_ctg18:2229234-2229235(+) | 3.41E-06 | 5.466893155 |

**Table S8.**

Significant SINE-RIPs for GWAS (body circumference)

| SINE-RIP | chromosome | ID | p_score | -log10(P) |
|---|---|---|---|---|
| SRIP-48777 | 2 | pig_ctg2:131051894-131051895(-) | 6.229335898e-06 | 5.20555825 |
| SRIP-53735 | 4 | pig_ctg4:110993604-110993605(-) | 2.508788654e-06 | 5.600535923 |
| SRIP-55828 | 5 | pig_ctg5:79361134-79361135(-) | 3.602633374e-07 | 6.443379932 |
| SRIP-58209 | 6 | pig_ctg6:115923318-115923319(-) | 8.534395292e-06 | 5.068827246 |
| SRIP-40807 | 15 | pig_ctg15:121407843-121407844(+) | 4.372820353e-06 | 5.359238364 |
| SRIP-11543 | 18 | pig_ctg18:14759288-14759575(+) | 2.035557231e-06 | 5.691316683 |

**Table S9. Protein-coding genes contain common SINE-RIPs associated with body size**

| Phenotype | SINE-RIP | Chromosome | Location | Insertion type | Candidate gene * | Reference transcript ID | Number of introns | Number of exons | Insertion region | Orientation relative to gene |
|---|---|---|---|---|---|---|---|---|---|---|
| Body weight | SRIP-29571 | chr1 | 271114588-271114589 | insertion | *NUP214* | ENSSSCG00000005711 | 35 | 36 | Intron 8 | Forward |
| | SRIP-17676 | chr4 | 75648999-75649292 | deletion | *PLAG1* | ENSSSCG00000006247 | 4 | 5 | Intron 1 | Reverse |
| | SRIP-17680 | chr4 | 75874803-75875073 | deletion | *LYN* | ENSSSCG00000006250 | 12 | 13 | Intron 1 | Reverse |
| | SRIP-55780 | chr5 | 77025651-77025652 | insertion | *SLC38A1* | ENSSSCG00000000807 | 18 | 19 | Intron 7 | Forward |
| | SRIP-30333 | chr10 | 31404571-31404572 | insertion | *FRMD3* | ENSSSCG00000010962 | 16 | 17 | Intron 4 | Forward |
| | SRIP-30900 | chr10 | 56399164-56399165 | insertion | *NRP1* | ENSSSCG00000011102 | 16 | 17 | Intron 3 | Forward |
| | SRIP-9383 | chr15 | 104894664-104894956 | deletion | *CASP10* | ENSSSCG00000026940 | 10 | 11 | 3'UTR | Forward |
| | SRIP-40531 | chr15 | 105764199-105764200 | insertion | *KIAA2012* | ENSSSCG00000016110 | 23 | 24 | Intron 14 | Reverse |
| | SRIP-42782 | chr17 | 10795636-10795637 | insertion | *ANK1* | ENSSSCG00000007022 | 43 | 44 | Intron 18 | Reverse |
| Body length | SRIP-17676 | chr4 | 75648999-75649292 | insertion | *PLAG1* | ENSSSCG00000006247 | 4 | 5 | Intron 1 | Reverse |
| | SRIP-17680 | chr4 | 75874803-75875073 | deletion | *LYN* | ENSSSCG00000006250 | 12 | 13 | Intron 1 | Reverse |
| | SRIP-55534 | chr5 | 64580231-64580232 | insertion | *VWF* | ENSSSCG00000063591 | 49 | 50 | Intron 15 | Reverse |
| | SRIP-30333 | chr10 | 31404571-31404572 | deletion | *FRMD3* | ENSSSCG00000010962 | 16 | 17 | Intron 4 | Forward |
| | SRIP-34990 | chr13 | 68339577-68339580 | deletion | *PPARG* | ENSSSCG00000011579 | 1 | 6 | Intron 1 | Forward |
| Body height | SRIP-16033 | chr3 | 59877939-59878206 | deletion | *DNAH6* | ENSSSCG00000045529 | 75 | 76 | Intron 70 | Reverse |
| | SRIP-54596 | chr5 | 7568092-7568093 | deletion | *XPNPEP3* | ENSSSCG00000033823 | 7 | 8 | Intron 1 | Reverse |
| | SRIP-55904 | chr5 | 83115089-83115090 | deletion | *ANO4* | ENSSSCG00000000873 | 25 | 26 | Intron 16 | Reverse |
| | SRIP-40382 | chr15 | 95048079-95048080 | insertion | *MFSD6* | ENSSSCG00000016052 | 6 | 7 | Intron 2 | Reverse |
| | SRIP-40597 | chr15 | 109811627-109811628 | deletion | *ADAM23* | ENSSSCG00000016131 | 24 | 25 | Intron 5 | Forward |
| | SRIP-9659 | chr15 | 130612553-130612813 | deletion | *DNER* | ENSSSCG00000016258 | 12 | 13 | Intron 4 | Reverse |
| Body | SRIP-48777 | chr2 | 131051894-131051895 | insertion | *SLC12A2* | ENSSSCG00000014255 | 27 | 28 | Intron 5 | Reverse |

| | | | | | | | | | | |
|---|---|---|---|---|---|---|---|---|---|---|
| circumference | SRIP-55828 | chr5 | 79361134-79361135 | insertion | *APPL2* | ENSSSCG00000000841 | 19 | 20 | Intron 1 | Reverse |
| | SRIP-40807 | chr15 | 121407843-121407844 | deletion | *DNPEP* | ENSSSCG00000016220 | 14 | 15 | Intron 1 | Reverse |

*Gene name abbreviates: Pucleoporin 214 (*NUP214*), Plyomorphic adenoma gene 1 (*PLAG1*), LYN proto-oncogene (*LYN*), Solute carrier family 38 member 1 (*SLC38A1*), FERM domain containing 3 (*FRMD3*), Neuropilin 1 (*NRP1*), Caspase 10 (*CASP10*), Ankyrin 1 (*ANK1*), Von Willebrand factor (*VWF*), Peroxisome proliferator activated receptor gamma (*PPARG*), Dynein axonemal heavy chain 6 (*DNAH6*), X-prolyl aminopeptidase 3 (*XPNPEP3*), Anoctamin 4 (*ANO4*), Major facilitator superfamily domain containing 6 (*MFSD6*), ADAM metallopeptidase domain 23 (*ADAM23*), Delta/notch like EGF repeat containing (*DNER*), Solute carrier family 12 member 2 (*SLC12A2*), Adaptor protein (*APPL2*), Aspartyl aminopeptidase (*DNPEP*).

**Table S10.**

map information for breeds

| Species | Type | Region_type | Abbreviation | Breed_name | Abbreviation | Sample size | Longitude | Latitude |
|---|---|---|---|---|---|---|---|---|
| Sus scrofa | Domestica_pig | Asia_domestica | AD | Bama | BM | 18 | 107.25589 | 24.143744 |
| | Domestica_pig | Asia_domestica | AD | Baoshan pig | BS | 3 | 99.159162 | 25.114711 |
| | Domestica_pig | Asia_domestica | AD | Dahuabai pig | DHB | 3 | 113.259294 | 23.130196 |
| | Domestica_pig | Asia_domestica | AD | Daweizi | DWZ | 3 | 113.238436 | 28.145077 |
| | Domestica_pig | Asia_domestica | AD | Debao pig | DB | 3 | 106.61192 | 23.326557 |
| | Domestica_pig | Asia_domestica | AD | Diannan Small Ear pig | DNXE | 10 | 100.796003 | 22.010755 |
| | Domestica_pig | Asia_domestica | AD | Enshi black | ES | 3 | 109.483308 | 30.274548 |
| | Domestica_pig | Asia_domestica | AD | Erhualian | EHL | 3 | 120.621288 | 31.311123 |
| | Domestica_pig | Asia_domestica | AD | Gansu Tibetan | GST | 10 | 103.733224 | 36.474436 |
| | Domestica_pig | Asia_domestica | AD | Gaoligongshan pig | GLGS | 3 | 98.85718 | 25.821118 |
| | Domestica_pig | Asia_domestica | AD | Guangdong Small Ear pig | GDXE | 3 | 110.91771 | 21.70813 |
| | Domestica_pig | Asia_domestica | AD | Guanzhuang Spotted pig | GZH | 3 | 116.589316 | 25.12112 |
| | Domestica_pig | Asia_domestica | AD | Guizhong Spotted pig | GZ | 3 | 106.614882 | 23.905361 |
| | Domestica_pig | Asia_domestica | AD | Hainan pig | HN | 3 | 109.712778 | 19.801667 |
| | Domestica_pig | Asia_domestica | AD | Hetao | HT | 3 | 108.270407 | 41.090083 |
| | Domestica_pig | Asia_domestica | AD | Huai pig | H | 3 | 118.338717 | 25.190186 |
| | Domestica_pig | Asia_domestica | AD | Jinhua | JH | 1 | 119.648649 | 29.108034 |
| | Domestica_pig | Asia_domestica | AD | Lantang | LT | 3 | 114.929044 | 23.416229 |
| | Domestica_pig | Asia_domestica | AD | Longlin pig | LL | 3 | 105.329102 | 24.70138 |
| | Domestica_pig | Asia_domestica | AD | Luchuan | LC | 3 | 110.263733 | 22.259792 |
| | Domestica_pig | Asia_domestica | AD | Meishan | MS | 3 | 121.126477 | 31.459037 |

| | | | | | | | |
|---|---|---|---|---|---|---|---|
| Domestica_pig | Asia_domestica | AD | Min | MZ | 4 | 126.627618 | 45.759363 |
| Domestica_pig | Asia_domestica | AD | Minbei Spotted pig | MBH | 3 | 117.832488 | 26.441067 |
| Domestica_pig | Asia_domestica | AD | Neijiang pig | NJ | 3 | 105.054491 | 29.582726 |
| Domestica_pig | Asia_domestica | AD | Ningxiang | NX | 3 | 112.365244 | 28.140423 |
| Domestica_pig | Asia_domestica | AD | Putian pig | PT | 3 | 118.850096 | 25.496588 |
| Domestica_pig | Asia_domestica | AD | Qinshaohua | QSH | 3 | 110.143549 | 27.659483 |
| Domestica_pig | Asia_domestica | AD | Rongchang | RC | 3 | 105.501709 | 29.464229 |
| Domestica_pig | Asia_domestica | AD | Saba | SB | 3 | 101.704785 | 25.432036 |
| Domestica_pig | Asia_domestica | AD | shaziling | SZL | 3 | 112.776832 | 27.638853 |
| Domestica_pig | Asia_domestica | AD | Sichuan Tibetan | SCT | 12 | 107.040769 | 30.460543 |
| Domestica_pig | Asia_domestica | AD | Tibetan Tibetan | TBT | 17 | 91.117301 | 29.654205 |
| Domestica_pig | Asia_domestica | AD | Tiegu | TG | 3 | 109.806804 | 28.712045 |
| Domestica_pig | Asia_domestica | AD | Tongcheng | TC | 3 | 113.811632 | 29.248167 |
| Domestica_pig | Asia_domestica | AD | Wannan Black pig | WNH | 3 | 116.630258 | 30.659234 |
| Domestica_pig | Asia_domestica | AD | Wuzhishan | WZS | 9 | 109.512805 | 18.777493 |
| Domestica_pig | Asia_domestica | AD | Yuedong Black pig | YDH | 3 | 114.460458 | 22.893788 |
| Domestica_pig | Asia_domestica | AD | Yunnan Tibetan | YNT | 11 | 99.701783 | 27.82234 |
| Domestica_pig | Asia_domestica | AD | Vietnamese native pig | VT | 3 | 106.701139 | 10.77639 |
| Domestica_pig | Europe_domestica | ED | Angler_Sattleschwein | AS | 2 | 9.822009 | 54.1854 |
| Domestica_pig | Europe_domestica | ED | British_Saddleback | BRS | 1 | 0.46467 | 51.770468 |
| Domestica_pig | Europe_domestica | ED | Bunte_Bentheimer | BB | 1 | 6.991562 | 52.454467 |
| Domestica_pig | Europe_domestica | ED | Calabrese | CALB | 1 | 16.421935 | 40.632874 |
| Domestica_pig | Europe_domestica | ED | Casertana | CST | 2 | 14.345778 | 41.073645 |
| Domestica_pig | Europe_domestica | ED | Chato_Murciano | CM | 2 | -5.385325 | 40.797611 |
| Domestica_pig | Europe_domestica | ED | Cinta_Senese | CS | 1 | 14.231308 | 40.850138 |
| Domestica_pig | Europe_domestica | ED | creole | CRL | 1 | -72.339593 | 18.547327 |

| | | | | | | | |
|---|---|---|---|---|---|---|---|
| Domestica_pig | Europe_domestica | ED | Gloucester_Old_Spot | GOS | 2 | -2.166667 | 51.833331 |
| Domestica_pig | Europe_domestica | ED | iberian | IB | 4 | -4.497437 | 39.8964 |
| Domestica_pig | Europe_domestica | ED | Leicoma | LCM | 1 | 11.279543 | 49.049026 |
| Domestica_pig | Europe_domestica | ED | Linderodsvin | LDD | 1 | 18.071093 | 59.325117 |
| Domestica_pig | Europe_domestica | ED | Mangalica | MALC | 6 | 19.146094 | 47.48139 |
| Domestica_pig | Europe_domestica | ED | Middle_White | MW | 2 | -1.38525 | 53.982527 |
| Domestica_pig | Europe_domestica | ED | Nanchukmacdon | NK | 1 | 126.978291 | 37.566679 |
| Domestica_pig | Europe_domestica | ED | Nera_Siciliana | SSL | 1 | 14.155048 | 37.587794 |
| Domestica_pig | Europe_domestica | ED | Retinto | RT | 1 | -5.204184 | 39.176465 |
| Domestica_pig | Europe_domestica | ED | Tamworth | TW | 1 | -2.007455 | 52.824694 |
| Domestica_pig | Europe_domestica | ED | Göttingen minipig | GO | 1 | 9.935181 | 51.532833 |
| Domestica_pig | America_domestica | ED | Yucatan miniature pig | YUM | 12 | -88.875567 | 20.684596 |
| Domestica_pig | Africa_domestica | ED | Angola | AGL | 4 | 13.243951 | -8.82727 |
| Domestica_pig | Europe_commercial | ED | Berkshire | BK | 3 | -1.031873 | 51.453489 |
| Domestica_pig | America_commercial | ED | Duroc | DU | 19 | -74.404162 | 40.075738 |
| Domestica_pig | Europe_commercial | ED | Hampshire | HAM | 3 | -1.243313 | 51.044981 |
| Domestica_pig | Europe_commercial | ED | Landrace | LR | 20 | 12.570072 | 55.686724 |
| Domestica_pig | Europe_commercial | ED | Pietrain | PIE | 20 | 4.917548 | 50.725484 |
| Domestica_pig | Europe_commercial | ED | Yorkshire | LW | 19 | -1.38525 | 53.982527 |
| Wild_boar | Asia_wild | AW | Wild_China | WC | 6 | 118.338717 | 25.190186 |
| Wild_boar | Asia_wild | AW | Wild_Northern China | WCN | 2 | 123.045639 | 42.049457 |
| Wild_boar | Asia_wild | AW | Wild_South China | WCS | 3 | 118.338717 | 25.190186 |
| Wild_boar | Asia_wild | AW | Wild_Japan | WJ | 2 | 139.762221 | 35.682194 |
| Wild_boar | Asia_wild | AW | Wild_Korea | WK | 10 | 129.075236 | 35.179953 |
| Wild_boar | Europe_wild | EW | Wild_France | WF | 1 | 2.348391 | 48.853495 |
| Wild_boar | Europe_wild | EW | Wild_Germany | WG | 4 | 11.575382 | 48.137108 |

| | | | | | | | | |
|---|---|---|---|---|---|---|---|---|
| | Wild_boar | Europe_wild | EW | Wild_Italy | WI | 5 | 7.682489 | 45.067755 |
| | Wild_boar | Europe_wild | EW | Wild_Netherlands | WN | 10 | 4.892453 | 52.37308 |
| | Wild_boar | Europe_wild | EW | Wild_Eastern Europe | WNE | 2 | 21.071432 | 52.233717 |
| | Wild_boar | Europe_wild | EW | Wild_Spain | WSP | 1 | 2.177432 | 41.382894 |
| | Wild_boar | Europe_wild | EW | Wild_Switzerland | WSZ | 1 | 7.452175 | 46.948474 |
| | Wild_boar | Europe_wild | EW | Wild_Ukraine | WU | 1 | 30.524136 | 50.450034 |
| Babyrousa babyrussa | - | Out_group | OG | Babyrousa babyrussa | BAB | 1 | 106.827049 | -6.175247 |
| Sus salvanius | - | Out_group | OG | Porcula salvania | PS | 4 | 85.320582 | 27.708317 |
| Sus cebifrons | - | Out_group | OG | Sus cebifrons | VW | 1 | 120.979996 | 14.590635 |
| Sus verrucosus | - | Out_group | OG | Sus verrucosus | JW | 1 | 109.613911 | -7.327969 |

**Table S11.**

Cross-validation-error-k of Common SINE-RIPs for admixture in 369 indivduals

| K | CV_error |
|---|---|
| 1 | 0.44874 |
| 2 | 0.40129 |
| 3 | 0.38618 |
| 4 | 0.37873 |
| 5 | 0.37393 |
| 6 | 0.37387 |
| 7 | 0.37048 |
| 8 | 0.37041 |
| 9 | 0.37029 |
| 10 | 0.3711 |
