## Supplementary Data1 for "TypeSINE: Genome-Wide Detection of SINE Retrotransposon Polymorphisms Reveals Functional Variants Linked to Body Size Variation in Pigs"

Fig.S1. PCR validation information

### PCR Validation Information Sheet

| NO. | SINE RIP ID | Genomic position | Primer-Forward | Primer-Reverse | MAF | Y/N | Gel electrophoresis |
|---|---|---|---|---|---|---|---|
| 1 | SRIP-11320 | pig_ctg17:60437863-60438152(+) | GCTAGCGAGGCACACTTCTT | CGTGCTGTCAGCTCATTCATTC | 72.55 | Y | 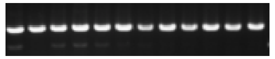 |
| 2 | SRIP-1072 | pig_ctg1:112889668-112889932(-) | AATTCTGGCTATGCTGCCTCA | CCACAGAGAGCTCGGCTATTC | 5.99 | Y | 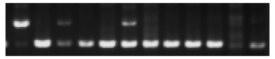 |
| 3 | SRIP-1308 | pig_ctg1:140548559-140548840(-) | CACAATGTGTCAAAGCGTTCCT | TGACCTGCACAGACTGATGTC | 6.15 | Y | 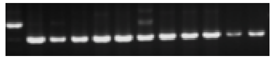 |
| 4 | SRIP-1859 | pig_ctg1:212062628-212062932(-) | GCTGCATACACATTTGCCTCT | CCTGAACAAATGAGACTGCGTT | 6.22 | Y | 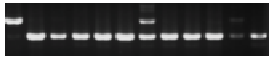 |
| 5 | SRIP-15628 | pig_ctg3:20877198-20877498(+) | CTGGTAGAAGGGCCAACAAGT | GAGCACCGTGTAGGGTTTACA | 7.58 | N | 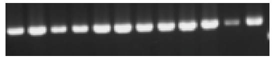 |
| 6 | SRIP-15903 | pig_ctg3:48517771-48518065(-) | GCTTTCTCGACAGTGGAATGA | ATTTTCCAGCGGATAACATGCC | 8.42 | Y | 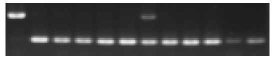 |
| 7 | SRIP-10414 | pig_ctg16:60892238-60892526(-) | CCATTGGACCACCGTAGTTCT | CATTGCATCCCAGCTCATCTC | 11.64 | Y | 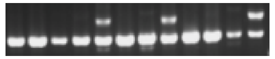 |
| 8 | SRIP-13689 | pig_ctg2:17491189-17491480(+) | GGGGACGTCACTGGATGTTT | TGGGATCTGACCCTGGTAGG | 13.28 | Y | 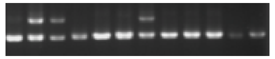 |
| 9 | SRIP-1277 | pig_ctg1:136090876-136091170(+) | ATCACCATTTGACTGCCCGT | CTCATCAACCCCCACTCACC | 13.66 | Y | 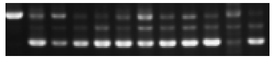 |
| 10 | SRIP-10487 | pig_ctg16:69047171-69047433(-) | ACAGAAAGGCCAGGGAGTTG | CCCAGGCTTTGCCAGTCATA | 16.8 | Y | 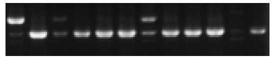 |

| | | | | | | | |
|---|---|---|---|---|---|---|---|
| 11 | SRIP-5428 | pig_ctg13:17797778-17798071(-) | AGCTTAAAAGCCAGGCCTCA | TTGCCTCCCCAGATGTCAAC | 17.08 | Y | 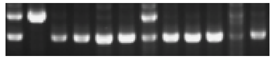 |
| 12 | SRIP-18707 | pig_ctg5:16328241-16328535(-) | ATGGGGATTAGAAACTGCTCTGT | ACCATAATTGGCTTGCCCTGA | 18.91 | Y | 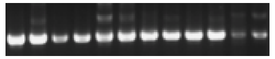 |
| 13 | SRIP-6689 | pig_ctg13:182972753-182973044(-) | CATTAACTTCATAATGGCTGTGGAC | AGAATCATTGGGCACTTCACCT | 28.45 | Y | 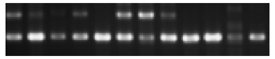 |
| 14 | SRIP-9945 | pig_ctg16:11476501-11476762(-) | AAGACATGAGTGGTATGGACTTT | CTGTCTTCAAAGGCAGTAGGGT | 30.96 | Y | 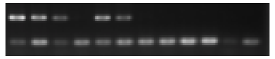 |
| 15 | SRIP-12387 | pig_ctg19:39709431-39709714(-) | GTTACACTCAAACACGGCCCC | AACAATAACTTTCCCCTTCTCCCA | 32.53 | / | 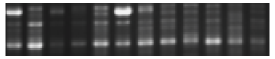 |
| 16 | SRIP-17110 | pig_ctg4:15785371-15785635(+) | AACTTCAGTGAGGAGACCACAT | CCCGTTTTATTCTGGTGAAGC | 40.55 | Y | 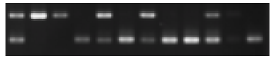 |
| 17 | SRIP-3446 | pig_ctg11:4738964-4739228(-) | CCCTCACGATGTGGAAAGATCA | AAAGGTTATCCTGCTCGCCC | 40.8 | Y | 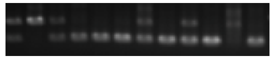 |
| 18 | SRIP-18536 | pig_ctg5:4016102-4016387(+) | ACTTCTCATGCGCGTAGGCT | TCCTCAGGGGTAATCGATGCTAT | 43.68 | Y | 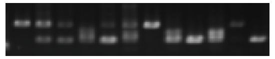 |
| 19 | SRIP-17204 | pig_ctg4:23686890-23687153(+) | ACCTTGAGAGTAAGGCTGAACTG | GCAAAGTGACCCAGTTTTGGATAG | 43.9 | Y | 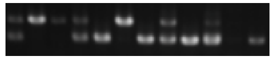 |
| 20 | SRIP-1217 | pig_ctg1:128388377-128388478(+) | GCCTCATGGATCCGGCTTTTA | AGCACACCACTCTTGGAACA | 50.68 | Y | 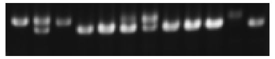 |
| 21 | SRIP-8345 | pig_ctg14:129931400-129931664(-) | ACATACTCCTAGCGTGCTCC | CCTAGGTACCAGGGAAACCC | 56.77 | Y | 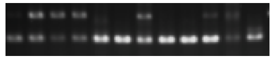 |
| 22 | SRIP-13172 | pig_ctg19:93141829-93142090(-) | TGAGTGAGTTGCGGACTGTG | ATACCAGGGGGAGTAGCTCA | 57.87 | Y | 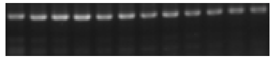 |
| 23 | SRIP-10828 | pig_ctg17:16616288-16616553(+) | ATTACCCGGATAGTGAGGGGG | TTCACAAAACTGCAAGAGGTCTATC | 61.11 | Y | 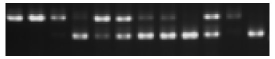 |

| | | | | | | | |
|---|---|---|---|---|---|---|---|
| 24 | SRIP-17900 | pig_ctg4:98833884-98834167(-) | CATGCTTCACTGTCTCCGGG | ATGTCAGTGAAAGCACAGTAAGA | 62.7 | Y | 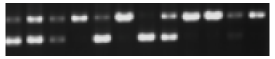 |
| 25 | SRIP-17498 | pig_ctg4:58621879-58622176(+) | CTCTAGTGTGCTAAGTCAGGGC | TGCATCTCCACGTGGTCTTTG | 64.33 | N | 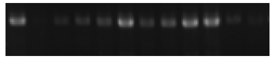 |
| 26 | SRIP-11730 | pig_ctg18:34002328-34002607(+) | TGATGTGTGGAGTCCCATTCG | ATGAGGTGCATCTTTTACATGGAT | 69.07 | Y | 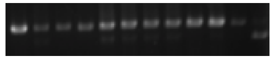 |
| 27 | SRIP-856 | pig_ctg1:85130205-85130498(+) | TCTGATGGCCCCAATTTTGTTTC | GACTAGGACTGTACTAACCTGCT | 69.89 | Y | 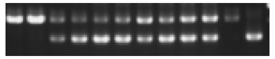 |
| 28 | SRIP-21541 | pig_ctg6:165807151-165807435(+) | CATGGTGTCTGCCATTGAGGA | TGGACACCTAGTGTGCGTCA | 74.51 | Y | 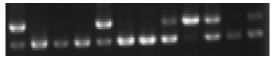 |
| 29 | SRIP-24700 | pig_ctg9:18584097-18584360(-) | CCAGGGTTGGCAATTCAGTCT | GACTTGGGCTTCAGCTCCTTAT | 75.47 | Y | 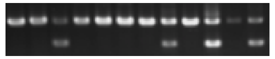 |
| 30 | SRIP-13547 | pig_ctg2:8262861-8263145(+) | AGCCCTAGCAGAGCAGTTTG | AGTCTATGATCTGGGGCTCTGT | 76.63 | Y | 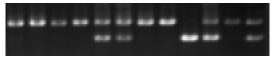 |
| 31 | SRIP-11114 | pig_ctg17:41732463-41732724(+) | GGAAGGACAGCATAGGAGAACC | CATTCCGCTACATGACCAACA | 79.84 | Y | 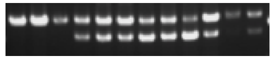 |
| 32 | SRIP-7668 | pig_ctg14:54976620-54976951(+) | CTGAGTGCTCCGAACGTGAC | AGGTGTGGTCTGCTACGAAA | 82.34 | Y | 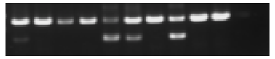 |
| 33 | SRIP-20793 | pig_ctg6:95664340-95664633(-) | ACACCTGAAACCAGCTATGTCC | GCTGATGGAACAGTCCAGTTG | 84.34 | Y | 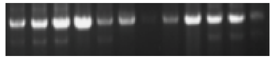 |
| 34 | SRIP-18596 | pig_ctg5:8232134-8232391(-) | TCCCCAGAAGTTTTCCTCGC | TGCTTCATAGACATCAAGAGGTAA | 84.75 | Y | 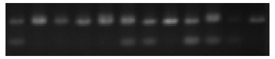 |
| 35 | SRIP-9874 | pig_ctg16:6489451-6489743(-) | TCACTCCACATACACTTGGCTC | CAAGACCACCCACGCATAAA | 85.33 | Y | 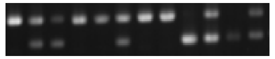 |
| 36 | SRIP-17164 | pig_ctg4:19195155-19195418(+) | GTGACTAGAACTCTCCAGTGC | CTTCCCACCAACAAGGGTAGT | 87.4 | Y | 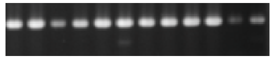 |

| | | | | | | | |
|---|---|---|---|---|---|---|---|
| 37 | SRIP-6734 | pig_ctg13:188726289-188726549(-) | TTTGCCTGTCAGGGTCACCAA | TGCTTTCCCTGACGACTCTATG | 88.48 | Y | 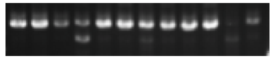 |
| 38 | SRIP-23112 | pig_ctg77:2019133-2019429(+) | CCTCCATGACACCGAAAAGC | CTCGTGCCGAGGTACAGAC | 91.94 | Y | 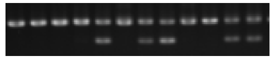 |
| 39 | SRIP-2439 | pig_ctg1:270816767-270817060(+) | TGGTGGAGCGGTAGGTTACA | ACAGCTGGGCAGCAGTACAT | 91.94 | Y | 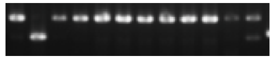 |
| 40 | SRIP-20184 | pig_ctg6:22623303-22623592(+) | CCCAAAAGGAATTTTGTTGCAC | GGCCAGCTAGTAAAAATGGTAGC | 92.35 | Y | 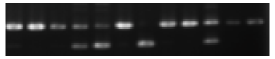 |
| 41 | SRIP-18831 | pig_ctg5:25670047-25670334(-) | CCATCATGACGTTAACCAAAAATG | TAAAACTGGGGCAACTTTGGGA | 92.76 | Y | 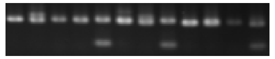 |
| 42 | SRIP-10010 | pig_ctg16:18439160-18439440(+) | GACCTCAAATGGCATTGCTCAG | TCACAATTCTGGTTTCATCCTGAC | 92.84 | Y | 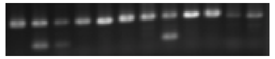 |
| 43 | SRIP-126319 | pig_ctg6:14860453-14860702(+) | GCAGACTTTATTCCTCAGGGCT | CTGGGTAGCCAGATTTAGGGATTA | 93.24 | N | 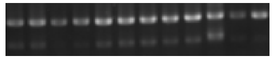 |
| 44 | SRIP-143034 | pig_ctg9:65521351-65521643(-) | CCCAATTCCCAGAGGGACAC | GGAGAAACCTGAGAAGGGGC | 93.48 | Y | 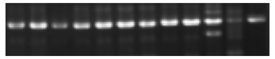 |
| 45 | SRIP-78983 | pig_ctg11:67891743-67892034(-) | TTTTGGTCTCCCAGGGTGTTT | TGCCCAGCTTTCCAGTAGTTA | 93.63 | N | 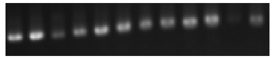 |
| 46 | SRIP-100147 | pig_ctg17:18225397-18225687(+) | CCTTCACGGAAGCAGTTCCT | AGGGGTACGTGACAATCTGC | 93.77 | Y | 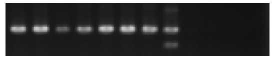 |
| 47 | SRIP-104215 | pig_ctg19:22630416-22630709(-) | GGCTTTTAAAGGCAAGACAAGGA | CCACCAGCAAATGCAGAGGT | 94.23 | N | 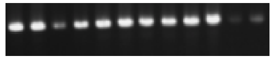 |
| 48 | SRIP-118189 | pig_ctg4:18561590-18561900(+) | TGTGCCATGTTTTTGGGGGAA | CTCTTTGTTTGCCCTAACTTGTGAT | 94.44 | Y | 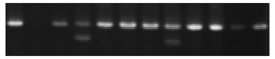 |
| 49 | SRIP-10902 | pig_ctg17:23253556-23253848(-) | AGCCAAATAGGAGCCCTGTA | GCTCAGAAAGTGCCTGCTGATA | 83.47 | Y | 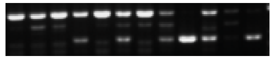 |

| | | | | | | | |
|---|---|---|---|---|---|---|---|
| 50 | SRIP-4832 | pig_ctg12:39871289-39871549(+) | TGGACTCCATCAAGGACACTC | CAGCAATATTTGGCCCTCCA | 93.24 | Y | 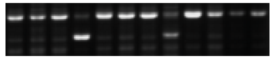 |
| 51 | SRIP-24829 | pig_ctg9:27050166-27050436(+) | ACCTAACAATGGCAAGGTGGT | AGCAAAAATAGCTGATTGGGTGA | 85.84 | Y | 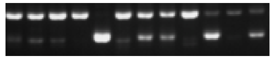 |
| 52 | SRIP-21536 | pig_ctg6:165716412-165716684(-) | TCGTAAGTGTCAGAGCCAGC | TCACCTGACACATTCTGACACA | 91.37 | Y | 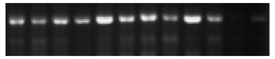 |
| 53 | SRIP-8862 | pig_ctg15:34869485-34869748(-) | CCACAGAAAGCCGTGTAGGT | TGGGATTTGATCCCCACAGC | 94.4 | Y | 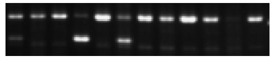 |
| 54 | SRIP-3218 | pig_ctg10:53425919-53426207(-) | ATTTTCGCAATAAAGCCCCC | GCAGACACCTCCCAGAAACT | 92.16 | Y | 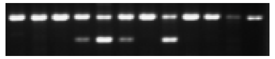 |
| 55 | SRIP-19698 | pig_ctg5:102041822-102042107(-) | CAGGCCGTCAGAGAGGATATG | GTCTCTGTGTGGCCGTAGTTA | 87.53 | Y | 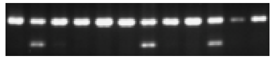 |
| 56 | SRIP-6703 | pig_ctg13:185304446-185304702(+) | TGCCAAAGACATACGTGACTATT | ATGCCCAAGGGTGAGCAATAA | 93.73 | Y | 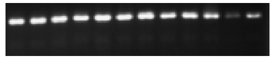 |
| 57 | SRIP-8732 | pig_ctg15:22083357-22083647(-) | GATCCATAGTGGGAGCTGGC | AGCAGGCTGTCTTGCTACAG | 80.83 | / | 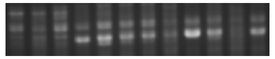 |
| 58 | SRIP-16570 | pig_ctg3:114498289-114498549(+) | TCGAATTGGACGGTCTGGTG | TGGTGCTCCAGGACAATCAC | 85.05 | ? | 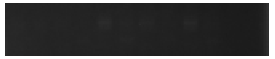 |
| 59 | SRIP-18551 | pig_ctg5:5260882-5261142(-) | CTGCGAATGTTCATAAGGGCG | AGAAGACGACGAGCTGTCAA | 82.15 | Y | 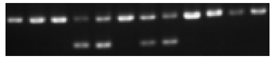 |
| 60 | SRIP-107765 | pig_ctg2:20976644-20976928(-) | ATCTCACCATGAGAAACTGTCCC | CCCAACCTTTGATCGCTCTG | 94.66 | Y | 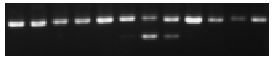 |
| 61 | SRIP-2052 | pig_ctg1:238672693-238672991(+) | CAGTGCAGGGTCTCATCAGG | CCTCTGGGGTTACTGCCAAG | 92.43 | Y | 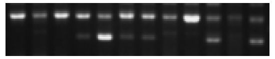 |
| 62 | SRIP-17898 | pig_ctg4:98748947-98749243(-) | TAAGTGATACTGCCCAGACTTCC | GGAGCAGTGGAAAGGTAAAATCG | 80.52 | N | 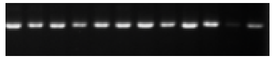 |

| | | | | | | | |
|---|---|---|---|---|---|---|---|
| 63 | SRIP-76253 | pig_ctg10:43141864-43142159(-) | TAGGGTAGAGAAATGTTTTGCGT | TTTACGGAAGAAGACCAGGCT | 94.57 | N | 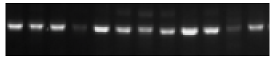 |
| 64 | SRIP-106 | pig_ctg1:10238054-10238342(-) | GGCTTTGCTCCCCATCAAAATA | GAAGATCCACAAGCCCGAGT | 80.14 | Y | 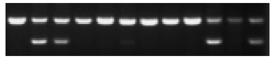 |
| 65 | SRIP-3728 | pig_ctg11:22279111-22279373(-) | GGCAGTCAGGGTACTGTCAT | CCCACTTGTGTGTAACAAAACAGA | 90.98 | ? | 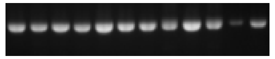 |
| 66 | SRIP-18584 | pig_ctg5:7696431-7696694(-) | TGAGTGAGAAACCCGAAGCC | AGGCCACTGTCATCCGAAAG | 85.04 | Y | 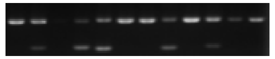 |
| 67 | SRIP-24287 | pig_ctg8:129813563-129813859(-) | CACAGAGGGTAGCAAGGGTG | AGGGAACAGAGCTAGCTGGA | 92.14 | Y | 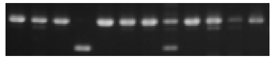 |
| 68 | SRIP-6921 | pig_ctg13:201004627-201004889(-) | TATCTCCGGGCGACATCTCT | AGTATGCTGTATGAACCCATGT | 82.44 | Y | 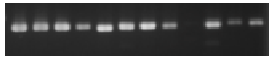 |
| 69 | SRIP-114 | pig_ctg1:10490266-10490558(-) | CTTTGTCTGTAGCTAATAAGCTGGT | ACCATCCAGGCAAATTCCAAAAA | 87.16 | Y | 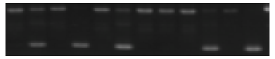 |
| 70 | SRIP-20488 | pig_ctg6:56443575-56443865(+) | AGCATTCTGCAGGGTACGAG | TGACATTCCGAGGGCACTTC | 88.51 | Y | 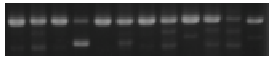 |
| 71 | SRIP-13223 | pig_ctg19:98609572-98609867(+) | ATCCCTTCAACACTTGGGTGC | AGTGCGCAGTAACTCTGTACC | 92.27 | Y | 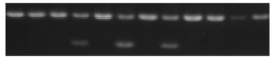 |
| 72 | SRIP-10351 | pig_ctg16:54432238-54432542(-) | TCACTTGGGGCTAACCTTGA | AACTGCAGTATTAAACACCCCC | 85.54 | Y | 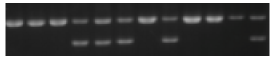 |
| 73 | SRIP-9172 | pig_ctg15:77631418-77631681(+) | CAGCAACAGGTAGGTGCGA | CACCCCTCGATGATACTGCAA | 92.16 | Y | 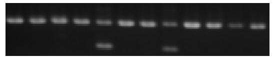 |
| 74 | SRIP-16427 | pig_ctg3:101562412-101562651(-) | TCCCCTATTTGCGAGCTGAAG | AGTCTATCAGGGGTCAGAGTGT | 90.65 | Y | 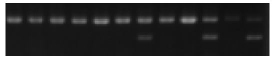 |
| 75 | SRIP-11388 | pig_ctg18:4316588-4316857(+) | GCCTGAAGCCGCCGTATATT | TGAAGCAACGACGAAGGAGA | 92.22 | Y | 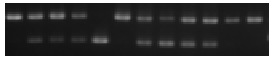 |

| | | | | | | | |
|---|---|---|---|---|---|---|---|
| 76 | SRIP-24029 | pig_ctg8:108516028-108516296(-) | GATGTGATCGGTCCCCTCAG | TGTCCAGATTTTGCTGGTACAGA | 87.91 | / | 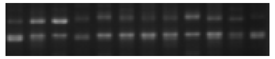 |
| 77 | SRIP-89612 | pig_ctg14:34864101-34864391(+) | TCAGCATCTGAGGGGCATTG | TGGGAAGTTGTGCAAAATGATGT | 94.7 | N | 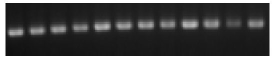 |
| 78 | SRIP-134548 | pig_ctg7:55226948-55227226(-) | TCTAAAGCAGTTGGGGTCCTC | GGATCCTGGGTATGCCTATCTG | 93.85 | / | 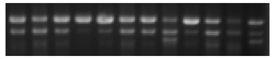 |
| 79 | SRIP-1714 | pig_ctg1:193258735-193259037(+) | AGGAACAACTGAAACGGTGGA | GTCCTGGACCTTTCAATACGG | 87.94 | Y | 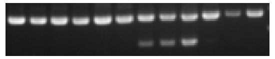 |
| 80 | SRIP-23039 | pig_ctg7:116738667-116738947(+) | TGGGTCAGGACAAACGTAGG | CCCGACAAGTCCCCTCAGAT | 80.54 | Y | 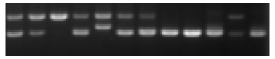 |
| 81 | SRIP-115926 | pig_ctg3:86039747-86040041(-) | TGGAACAGAATCTGGCAGAGTAG | AGTGTACTGCTCTACTGGAGGAAG | 94.46 | N | 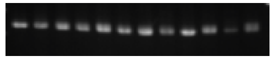 |
| 82 | SRIP-11386 | pig_ctg18:4162335-4162619(+) | AAACATTGAACCGTGACGCC | GAGGTCACTGTCACTAGGCTG | 93.19 | Y | 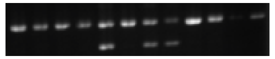 |
| 83 | SRIP-1714 | pig_ctg1:193258735-193259037(+) | AGGAACAACTGAAACGGTGGA | GTCCTGGACCTTTCAATACGG | 87.94 | 重复 | 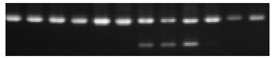 |
| 84 | SRIP-25471 | pig_ctg9:102350764-102351046(+) | CATCCTAGTGGAGGGCAGTT | GCTGAAGAACCTGGCTAGATG | 85.83 | Y | 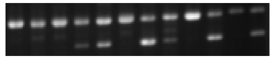 |
| 85 | SRIP-4040 | pig_ctg11:56755421-56755709(+) | ATGCTGTGACCGATCGCATT | ACCTGTAGATCATTTGAAGTGGG | 91.87 | Y | 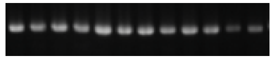 |
| 86 | SRIP-10188 | pig_ctg16:33552465-33552728(+) | GGGGGACAAGTCCCAATATGT | AAGATGACAGACCTTCCCCACT | 90.8 | Y | 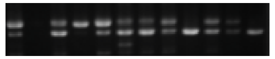 |
| 87 | SRIP-83211 | pig_ctg13:35245632-35245921(+) | TAGCGCAGGTGTAAATGGCA | CCTTGGGTTTACAGGTGATTGC | 93.9 | Y | 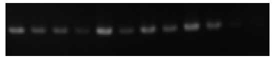 |
| 88 | SRIP-24099 | pig_ctg8:113353508-113353771(-) | AACAACTATGGTGCCGTGTG | CTCTGCACACCTGACGTTTCT | 81.84 | Y | 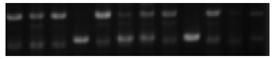 |

| | | | | | | | |
|---|---|---|---|---|---|---|---|
| 89 | SRIP-15053 | pig_ctg20:2794715-2794978(+) | CCAGAGTGAGGCAAGTGTGATA | CCTTCGGGGAACCTCTACG | 92.88 | Y | 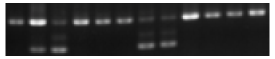 |
| 90 | SRIP-2566 | pig_ctg10:7621911-7622201(-) | AGCTGCTGATCAAGTATGCCA | CCACAGTTTAGGAGTTTGTGCAG | 94.02 | Y | 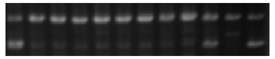 |
| 91 | SRIP-23120 | pig_ctg8:549436-549696(-) | CAGAAACCTCCTTCGCGCTT | ATGGGCTATTCAGCCATCGTT | 83.56 | / | 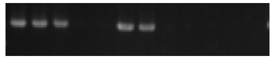 |
| 92 | SRIP-19933 | pig_ctg6:5603297-5603587(+) | ACCCCATCGCCTCATATGTT | GGTCAAGCTCACTCGGGAC | 94.32 | ? | 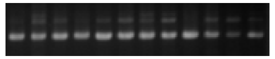 |
| 93 | SRIP-4403 | pig_ctg12:11481790-11482079(+) | CCACGAGCTTCACCTGACAT | ATCTCCTTTCCAACCGTCCG | 94.64 | Y | 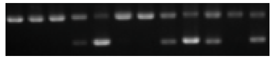 |
| 94 | SRIP-12728 | pig_ctg19:61931130-61931391(-) | TGCCACTGTTTGGCATAAGAA | GGGAAGGTACTTCAGGCTTGTC | 92.84 | Y | 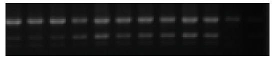 |
| 95 | SRIP-6004 | pig_ctg13:88378035-88378296(+) | TCTGCACAATGTATTCCCGTCA | CAGACTCATATGTTTAGCAGAGTGG | 91.87 | Y | 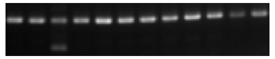 |
| 96 | SRIP-8142 | pig_ctg14:111528063-111528319(-) | ACTGATTAAACTCCCAAGTCCCT | AGATATGGGGATATGGGCAAGTG | 91.98 | / | 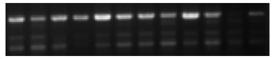 |
| 97 | SRIP-16313 | pig_ctg3:89054380-89054674(-) | AATCCTGACACACCCCAAGT | CGTTCTGTTAGAGCAGAGCAGG | 81.39 | Y | 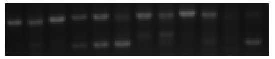 |
| 98 | SRIP-6880 | pig_ctg13:198096716-198097002(-) | ACATGCCCGATGGCTTCATA | CCGAAGCGTCATCACGAGAG | 85.19 | Y | 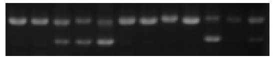 |
| 99 | SRIP-66820 | pig_ctg1:17112019-17112312(-) | ACCCTCTCTGATTGGGTGACATTA | GGAGTGGCTATGGGTGGATAATA | 93.96 | N | 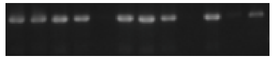 |
| 100 | SRIP-11894 | pig_ctg18:46135692-46135956(-) | TGCGCATGCGTAGTCATTCTA | TTGTATGCAGCAAGGACGTT | 91.21 | Y | 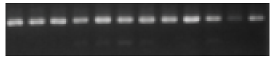 |
| 101 | SRIP-10003 | pig_ctg16:17589938-17590227(-) | ACAGGTTGTGGCTCCTTAGAA | AAACAATGGGGCACCCGAAC | 81.49 | Y | 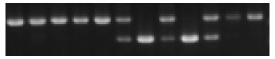 |

| | | | | | | | |
|---|---|---|---|---|---|---|---|
| 102 | SRIP-23161 | pig_ctg8:5442008-5442271(-) | GGATCAGAGGAAAATGACGCA | TCCAGACAAATCACGCCGAC | 87.91 | Y | 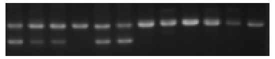 |
| 103 | SRIP-11815 | pig_ctg18:40858291-40858580(+) | TGGTGGTATGAACGGAGTGC | TCCTCTACGGCAGCTCATCT | 94.8 | Y | 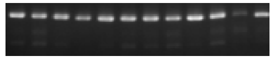 |
| 104 | SRIP-7862 | pig_ctg14:83757720-83758005(+) | TGGTTGAGCCTAGGATGGTG | TGAGTCACAGGAATCCATGCT | 87.57 | Y | 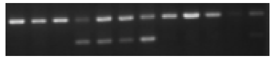 |
| 105 | SRIP-3153 | pig_ctg10:49712665-49712924(+) | TTCCATAGTGCGAAACCCCC | TGGCTGGTGGGATTTACAAAGA | 92.84 | Y | 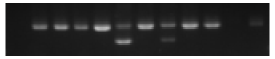 |
| 106 | SRIP-44 | pig_ctg1:6375871-6376132(-) | CCTCAGCTCTATGTCAGCGT | GCTCTCGCTGTGTTTGTGAG | 86.72 | Y | 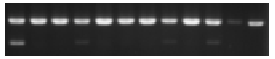 |
| 107 | SRIP-2043 | pig_ctg1:238353725-238354012(-) | TTAAATGTGTGCCGATGCCTTC | ATGAGGATGGACTAAAAGGGGC | 84.15 | Y | 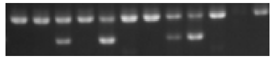 |
| 108 | SRIP-22562 | pig_ctg7:76863858-76864138(+) | TTGCGGGCAACAAATCAAGG | TGAGCACTGAAAGGGCTCTG | 82.19 | Y | 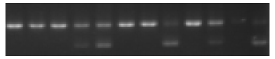 |
| 109 | SRIP-14043 | pig_ctg2:51412918-51413181(+) | ACTGGGCTAGAACCTACCCT | TGGTCTTTAATGTTCTGTTTCTGTG | 80.65 | Y | 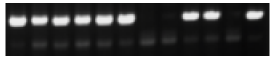 |
| 110 | SRIP-17855 | pig_ctg4:95322037-95322314(-) | TTAAGGACAAACTCACCCCCAAA | AGGTAATGATGCTTTGGCACC | 94.16 | Y | 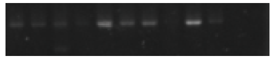 |
| 111 | SRIP-8120 | pig_ctg14:109307025-109307288(+) | TCACACCACATGGTCCACGA | GCTTTCCAGTTTGCTGCTACTC | 92.62 | Y | 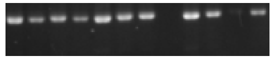 |
| 112 | SRIP-23713 | pig_ctg8:61258918-61259181(-) | AGAATGCCTTAGCATGAGCTG | GTTCTGACCCTACTACTCCCTG | 86.28 | Y | 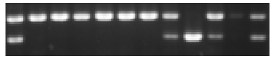 |
| 113 | SRIP-9848 | pig_ctg16:5355053-5355304(-) | AGTGGCTGGCTAATGTCCAAA | GCTTACCTCACTGTTAGCTGGT | 93.44 | Y | 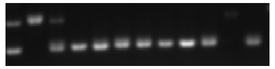 |
| 114 | SRIP-3683 | pig_ctg11:20834591-20834854(-) | TAGAGGAACACCCACCCACA | TCTGTTTCCTTTCCATTATCAGTCA | 91.87 | Y | 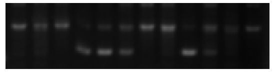 |

| | | | | | | | |
|---|---|---|---|---|---|---|---|
| 115 | SRIP-20672 | pig_ctg6:83455823-83456084(+) | AGGGCTAAAAGCTCAGCCAG | AGGATTTTTGTCCAATAGTGATGC | 92.76 | Y | 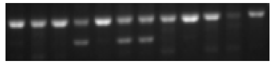 |
| 116 | SRIP-20199 | pig_ctg6:24699171-24699413(+) | TCCGAAGGTAACAACACAGAGA | AGGGCATTGCAGGACACAAT | 93.47 | N | 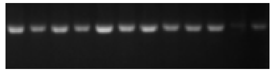 |
| 117 | SRIP-21215 | pig_ctg6:145169409-145169701(+) | TTGTGAGGGCGAATTAGGTCA | CCCCATAAGTAGGTGCCCTG | 89.05 | Y | 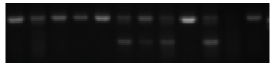 |
| 118 | SRIP-22664 | pig_ctg7:85184285-85184589(+) | TTGGTTGGTGTGGAAGGTTT | GTTGGGGATACTGGACTTGGG | 82.25 | Y | 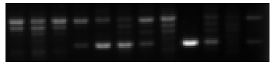 |
| 119 | SRIP-6332 | pig_ctg13:135360493-135360750(-) | CTGTACTGTGTCAACATGCAGTC | CTTGGTTCCTACCTGGAGAGG | 92.64 | N | 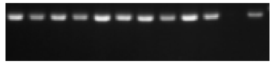 |
| 120 | SRIP-384 | pig_ctg1:30431302-30431565(+) | AGCCTTCAGAAGAGCAACTGG | TCTCTGATTGGCCAAGTTTTCAG | 81.92 | Y | 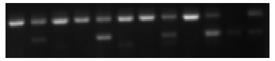 |
| 121 | SRIP-22065 | pig_ctg7:31357373-31357654(-) | TGCACATTCAGGGTTACCTGC | TGACATCCTTCAGGGCTACG | 89.32 | Y | 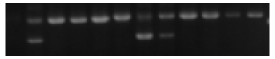 |
| 122 | SRIP-11561 | pig_ctg18:15746192-15746453(-) | AGCTTGCCCTCATCATCACC | AGCACTGGCATAGAGGAGGA | 90.11 | / | |
| 123 | SRIP-23612 | pig_ctg8:45651851-45652151(+) | AACTGACATGGCACTATTTGCT | TCCAACCGTATATATGCCCCAC | 81.52 | Y | 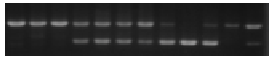 |
| 124 | SRIP-15204 | pig_ctg223:1060060-1060352(-) | TACACTCCCTCCTCGCTACG | TAGCCAAACCGATGGTGGTG | 81.2 | Y | 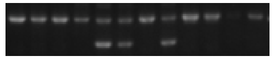 |
| 125 | SRIP-1932 | pig_ctg1:223524733-223525037(-) | CCTCTAGTGGCCTAACGTGG | TTCCCGTCAACCGGAAACAT | 83.11 | Y | 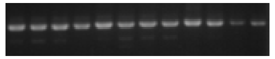 |
| 126 | SRIP-10783 | pig_ctg17:13934088-13934389(-) | CTCCCAAACATGAGCACAACC | CCAACAGCGGTAGTTAATGGC | 85.07 | N | 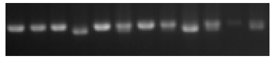 |
| 127 | SRIP-97589 | pig_ctg16:16460581-16460875(-) | TATCCCTCTGACTGTGACACAAC | GACAACTCGACAGACACTTGGT | 93.63 | Y | 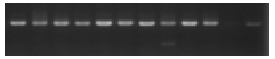 |

| | | | | | | | |
|---|---|---|---|---|---|---|---|
| 128 | SRIP-20651 | pig_ctg6:81594174-81594438(-) | TCCACTTACTTGGGAGATGGATTG | TGCTTAGAACTGAGGGCTATGT | 90.54 | Y | 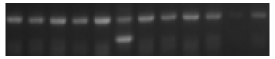 |
| 129 | SRIP-11397 | pig_ctg18:4594743-4595006(-) | ACTCCGTTACCAGTGGGACT | TCCAGCCAATGAACTGTCGG | 92.31 | Y | 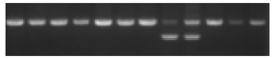 |
| 130 | SRIP-17125 | pig_ctg4:16567426-16567694(-) | CCATGGGATTATCATGGACAATTT | AGGACATGCAGCAAGAGCAT | 81.73 | Y | 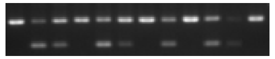 |
| 131 | SRIP-19218 | pig_ctg5:63964547-63964837(-) | ACCCCATCAGACAGCTTTGTA | AGCTAGCATCGCGAATGGA | 85.19 | Y | 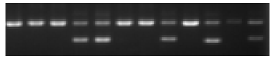 |
| 132 | SRIP-12356 | pig_ctg19:34513594-34513856(+) | TAACAGTATCCGGCAGGCAA | GGAGTCTTGACAGCTTTCAGTTTT | 86.48 | Y | 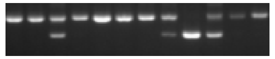 |
| 133 | SRIP-25631 | pig_ctg9:116337463-116337757(+) | TCCAGGGATTCAGGGTGGTA | ATCTGGAAAAGCATTCAATATAGCC | 88.99 | N | 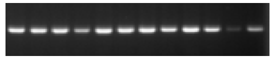 |
| 134 | SRIP-130547 | pig_ctg6:111557744-111558046(+) | GGCATGGGAAGGGTATGCTAT | GGTGACAGTTTGAGCCGAAG | 94.99 | Y | 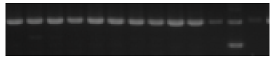 |
| 135 | SRIP-24349 | pig_ctg8:132369527-132369811(+) | GCAGGAGAGCTTTAAACGGG | TGCTACAAGCATTGCGTTCA | 88.96 | Y | 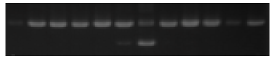 |
| 136 | SRIP-21195 | pig_ctg6:144162392-144162685(+) | CTTCCACGAAGGTATCCAGAAGT | GCTTTAGTTTTGCAACTGCCCT | 84.65 | Y | 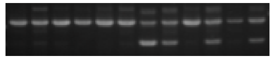 |
| 137 | SRIP-1209 | pig_ctg1:127349444-127349714(-) | GCTACCCCAGTTCAACCTTTCA | GGTTGACCTCTGCACTTCTACT | 81.03 | Y | 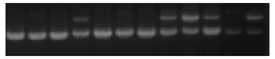 |
| 138 | SRIP-53184 | pig_ctg4:71568494-71568495(-) | GGACACTCCCTAGGCTAGAAAAG | TCACAAGACTTCTCCCACCAA | 5.74 | Y | 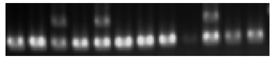 |
| 139 | SRIP-28861 | pig_ctg1:235950327-235950328(+) | TATCTTTGTGGCTAGAATTGTTCCT | TCACCTGACACCCTGTCAAA | 5.18 | Y | 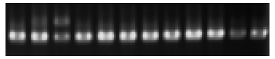 |
| 140 | SRIP-34703 | pig_ctg13:42215471-42215472(+) | CAAGCAATTCATTGGCTGGTT | GTCGTGGGAGCTAGTGTACG | 5.35 | Y | 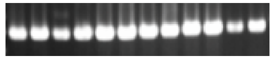 |

| | | | | | | | |
|---|---|---|---|---|---|---|---|
| 141 | SRIP-33913 | pig_ctg13:3375142-3375143(+) | CAAATTCACCCTCCAACACCAG | TCAGGTGGATCCGGTAAGACT | 6 | Y | 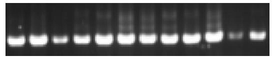 |
| 142 | SRIP-40064 | pig_ctg15:67544045-67544046(-) | CGTCAGAGATGAGGTACTGGC | CAAGGAGGCAGAGGAACGTA | 6.66 | Y | 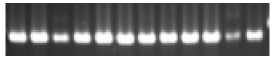 |
| 143 | SRIP-56257 | pig_ctg5:99808036-99808037(-) | TGCTAGGCATTCTACTTGTCCC | ATAAACACGTGGGCATGGTGA | 7.3 | Y | 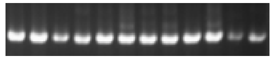 |
| 144 | SRIP-45005 | pig_ctg19:4426093-4426094(-) | ACTTTAGACATTCTTGCACGCTT | TGAGATAGGCGAGTACACCAA | 7.66 | Y | 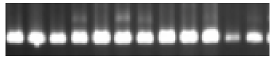 |
| 145 | SRIP-48343 | pig_ctg2:105477729-105477730(-) | CCTTAATCCCTGGTGTCAGAG | CACAGGTGCTAATAATGACAGGC | 8.36 | Y | 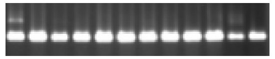 |
| 146 | SRIP-28231 | pig_ctg1:178494516-178494517(+) | AATGCGCAACTGGAAAATTAGG | GCAAGGTTGGAGCTGATGAAT | 9.51 | Y | 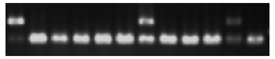 |
| 147 | SRIP-63997 | pig_ctg9:21712497-21712498(+) | ACATGAAGGCGTTGTTCTGGT | ACAACGCAGGTCTTGTGGTAA | 9.62 | Y | 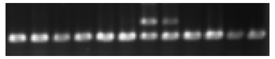 |
| 148 | SRIP-38285 | pig_ctg14:97472775-97472776(-) | CCCAGGAACATGTGCGCTAT | TGAGAATTCTGGGCTGAGCTATT | 10.66 | Y | 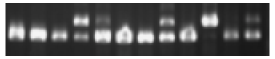 |
| 149 | SRIP-55762 | pig_ctg5:76157128-76157129(+) | ACAGGGCTGTGGAGTGTATAG | TGAAAACGTATGGTGGCAGC | 10.98 | Y | 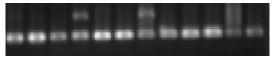 |
| 150 | SRIP-57820 | pig_ctg6:86264007-86264008(+) | CACCGGAACGACACACACAA | GCTCGACAACGGGGAGAAAA | 11.92 | Y | 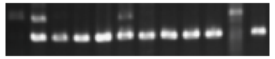 |
| 151 | SRIP-53367 | pig_ctg4:86042773-86042774(+) | CAGGTAAGCCTTCGGACCAG | TTTGATCCCCATCAGGGTGC | 11.92 | Y | 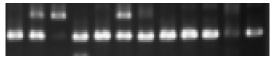 |
| 152 | SRIP-37702 | pig_ctg14:49357131-49357132(+) | CTCACAAGGTTTCAAGCCGC | CATGACAGAATTCTTCCTTGGCA | 12.33 | Y | 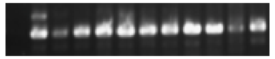 |
| 153 | SRIP-32866 | pig_ctg12:17137617-17137618(+) | ACCTGCGGCAGATAGAATGT | CTGTCTCCATGCCCCTAAAACA | 12.47 | N | 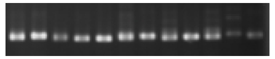 |

| | | | | | | | |
|---|---|---|---|---|---|---|---|
| 154 | SRIP-38328 | pig_ctg14:99712774-99712775(+) | ACACTTTCTCAGTTACGTTGAGAG | AAGCTCAGTGAAAATGACTTGC | 12.5 | Y | 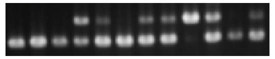 |
| 155 | SRIP-29316 | pig_ctg1:261115585-261115586(-) | AGTTTTCAGCTCATGTGACCAA | ATTCTATGTGGGAGCCTCTGAA | 12.97 | Y | 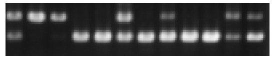 |
| 156 | SRIP-27198 | pig_ctg1:81430432-81430433(-) | ATCTCAAAGACCACCAGAACCAG | TGGCATGTGCACTCAACGA | 13.7 | Y | 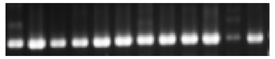 |
| 157 | SRIP-39454 | pig_ctg15:22570249-22570250(+) | GAGGAATTCGTGCTCCGAAA | AGGGCATCTAGGTCAACTTGC | 14.79 | Y | 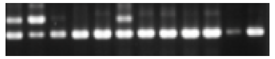 |
| 158 | SRIP-33721 | pig_ctg12:55832676-55832677(+) | CTAGCTGCAGGCGAGGTTAG | GCCATATAGACAAATCTCCCATGAT | 15.4 | Y | 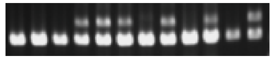 |
| 159 | SRIP-44209 | pig_ctg18:20611607-20611608(+) | GCTCGATGATAGCTGCGTGA | ATGGCTACCTCCCATACCGT | 15.85 | Y | 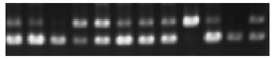 |
| 160 | SRIP-51471 | pig_ctg3:112783601-112783602(+) | TCCAGTCCAAAATGCCCGTG | TGCGAGACTGTCAATACTGGG | 16.67 | Y | 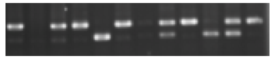 |
| 161 | SRIP-62793 | pig_ctg8:105867773-105867774(-) | TATTTGACTGCAGTTTCCACAGC | ATGGTATCTGGTATCCAGTATGCAG | 17.19 | Y | 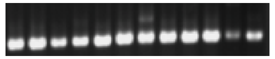 |
| 162 | SRIP-47320 | pig_ctg2:25664774-25664775(+) | ATCATCCATGACCGCAGCTT | CCTACACGCTTTGTTGCGAG | 17.26 | Y | 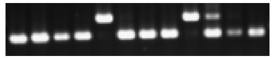 |
| 163 | SRIP-30960 | pig_ctg10:59819273-59819274(-) | AATTTAGACTGGGCTTGCGTG | TCCATGCACATACCATAAGCAAA | 17.93 | Y | 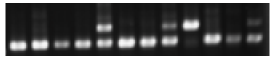 |
| 164 | SRIP-31684 | pig_ctg11:24790429-24790430(-) | GGGCAGATGCTACCTAGAGA | AGTCCCAAATCCAGAGCATAGC | 18.13 | Y | 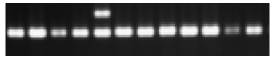 |
| 165 | SRIP-54554 | pig_ctg5:6027170-6027171(+) | TGTCGTTCTTACGAGGACGC | GTCTCATCGGTTTGCTTCGG | 18.99 | Y | 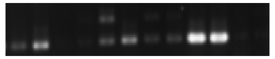 |
| 166 | SRIP-34928 | pig_ctg13:62153700-62153701(-) | GGCACTAGGCTATCTAGGCAC | CCTCGCTGTAGCATGACCTA | 19.26 | Y | 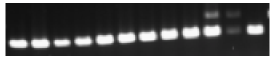 |

| | | | | | | | |
|---|---|---|---|---|---|---|---|
| 167 | SRIP-51692 | pig_ctg3:124760143-124760144(-) | CTTCCTACGTGTGGGTGTCA | GCCACCATAACTGGTGCAGA | 21.51 | Y | 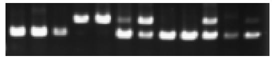 |
| 168 | SRIP-45534 | pig_ctg19:43937933-43937934(+) | GAGAGTGCGGAGGCAACTT | CAGCAAGAAACTGAGTAGGGC | 21.64 | Y | 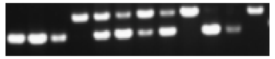 |
| 169 | SRIP-41516 | pig_ctg16:15762652-15762653(-) | TCTGTGTATGTCTACTGGTCCTTG | TGCTGCTTCTTGCTTCTTTGGG | 21.92 | Y | 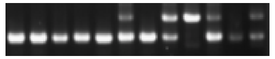 |
| 170 | SRIP-26945 | pig_ctg1:57480575-57480576(-) | GTGAGCAATGGGTCCCTACAA | TTAAGGCTCTGGGAAGTTACTGA | 26.1 | Y | 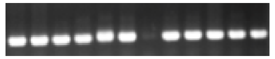 |
| 171 | SRIP-46791 | pig_ctg19:124855822-124855823(+) | TCCGTAAGTGCAGTTCAATCTCA | CGAGTAGCTACCTGCAAGTCAA | 31.67 | Y | 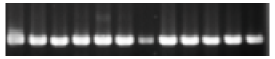 |
| 172 | SRIP-64736 | pig_ctg9:64701476-64701477(-) | TTAGGCTCTGGGCTCTCTCTATC | AGATGCATTGTCGACTCTCTC | 32.61 | Y | 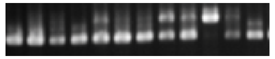 |
| 173 | SRIP-44765 | pig_ctg18:48683505-48683506(-) | GGCTTGAGGGCCGAAACATC | TAGAAAAGTGCTAACGTCGCCT | 34.87 | Y | 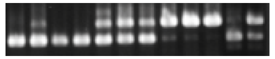 |
| 174 | SRIP-27334 | pig_ctg1:92964314-92964315(+) | GACCCAGTGGCAGAAGAAGT | TCTTGCTTTCCCATCACCAA | 35.25 | N | 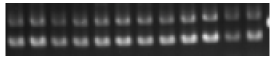 |
| 175 | SRIP-35146 | pig_ctg13:83100035-83100036(-) | CCCTAACCCCTAGGCACCTTA | AGAAGTTGAGGAAGGACTGCT | 36.2 | Y | 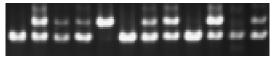 |
| 176 | SRIP-65534 | pig_ctg9:124814721-124814722(-) | ACGAAGACTGGGATCTACGC | GTTACAAGCACAAGTTGCCA | 36.94 | Y | 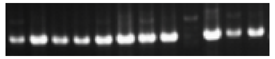 |
| 177 | SRIP-35892 | pig_ctg13:152145292-152145293(-) | GTAAGCTGTCTTGCGATTTTCAA | TCCTTTTTCCCCCACAAAGCTA | 38.5 | Y | 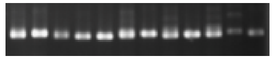 |
| 178 | SRIP-36773 | pig_ctg14:621524-621525(-) | CGGGACAGGATGTTGGTTGA | CATCGGTGTGACGACATCCA | 39.84 | Y | 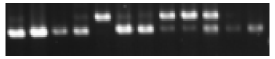 |
| 179 | SRIP-28500 | pig_ctg1:206178956-206178957(-) | GTGATGGAAACTTGGGGTACT | TCCTTACACAGATGGCGGAC | 41.96 | Y | 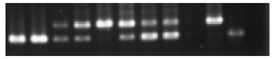 |

| | | | | | | | |
|---|---|---|---|---|---|---|---|
| 180 | SRIP-44208 | pig_ctg18:20432496-20432497(-) | CCTTCCCATGATTTCCCCGT | ACAGCTTAGAATTGCACGCC | 47.15 | Y | 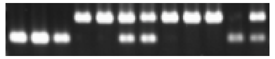 |
| 181 | SRIP-28795 | pig_ctg1:229742661-229742662(-) | TGGCTTGTCAACGATGCACTG | GAGGCCTCCCCAGTAATAAGG | 50.14 | Y | 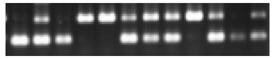 |
| 182 | SRIP-63586 | pig_ctg9:5290244-5290245(+) | CCTGTTCCTCCTTTCTCTTAGCTTT | AGCATATTTGGCAGTTTCAGTTCT | 51.23 | Y | 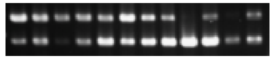 |
| 183 | SRIP-56637 | pig_ctg6:7233485-7233486(-) | AGTGGATACTGGCGGACTCT | GGCTCTAGAACTGAGCCGTC | 85.91 | Y | 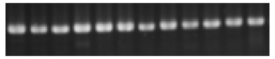 |
| 184 | SRIP-58455 | pig_ctg6:135213452-135213453(-) | CAGCTTCTGGACATGTAACGTA | GCTGGGAGGACAATTTACAGAAAG | 90.17 | Y | 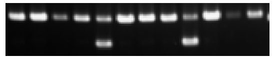 |
| 185 | SRIP-47434 | pig_ctg2:29859264-29859265(-) | AATCCTACCTCCACCCAGTTGA | AAGGCTCTGAATGTGCTACG | 90.41 | Y | 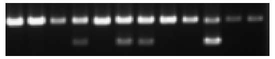 |
| 186 | SRIP-45438 | pig_ctg19:39007432-39007433(-) | ATGGTGCTCACCTCTTGTCTT | GAAAATGTGCGCAAGGAGGTC | 17.35 | Y | 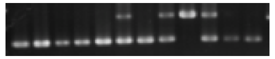 |
| 187 | SRIP-43699 | pig_ctg17:53084836-53084837(+) | TTTCCACCGAAACAGACGGG | ACTGGTCTTAGAGATTTGCCCAT | 17.9 | N | 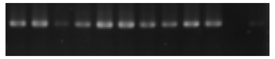 |
| 188 | SRIP-28269 | pig_ctg1:183870023-183870024(+) | AGTTCCCTCAGGCGGTACTA | ATCTCTGCCCCATTGAAGAGC | 19.32 | Y | 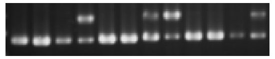 |
| 189 | SRIP-46771 | pig_ctg19:122352677-122352678(-) | CTGGGTCTTCACGATGGGTC | CAGAATAGTGTGCCTTCATTTATGT | 8.82 | / | 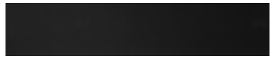 |
| 190 | SRIP-31715 | pig_ctg11:25429691-25429692(+) | TGATGGGGCAGAAAGATGATGA | TACAGTTAGGTGGATTGCCTGC | 6.67 | Y | 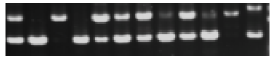 |
| 191 | SRIP-62454 | pig_ctg8:77322953-77322954(+) | GTGGGTCGTAAGTGGTCCTG | ATCTTCATCCTGCTGCCCAC | 14.44 | / | 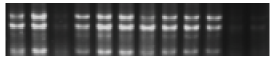 |
| 192 | SRIP-32328 | pig_ctg11:65681431-65681432(-) | GCTATTTGTGGTCAACCCGC | CGTCCATGAATCACGTCCCT | 14.9 | Y | 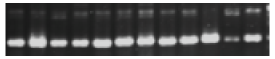 |

| | | | | | | | |
|---|---|---|---|---|---|---|---|
| 193 | SRIP-34194 | pig_ctg13:13387267-13387268(-) | CACTCTATGCCAGCCCATGT | GAGGAGGAGGACCCAGGAAA | 14.82 | / | 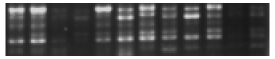 |
| 194 | SRIP-60771 | pig_ctg7:93211929-93211930(+) | GCTGTAAAAGGCTTATCACTCTCAG | AAGAGCCTGTAGCCAAACACC | 7.61 | Y | 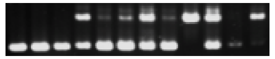 |
| 195 | SRIP-415618 | pig_ctg17:29913068-29913069(+) | AGAGTCATCCCAGGAGTGGTAG | AGATTAGTGACGCAAGGGGT | 5.16 | Y | 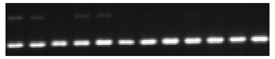 |
| 196 | SRIP-65422 | pig_ctg9:120090000-120090001(-) | TCATGCTTCCTTCTGGAGTGG | TAAGTGTTACCCCGCCCCAT | 15.22 | Y | 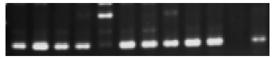 |
| 197 | SRIP-58877 | pig_ctg6:155349902-155349903(-) | AGGCCTACCGTGAGCAAAAA | TCAGGCGCTTCACCTCATC | 18.39 | Y | 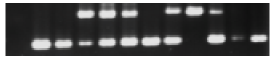 |
| 198 | SRIP-50615 | pig_ctg3:55155945-55155946(+) | ATGGTACCAGTGAGGTACGG | TGCTCACCTGGCACTTACG | 13.04 | Y | 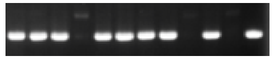 |
| 199 | SRIP-30835 | pig_ctg10:54166236-54166237(-) | ATTGGCCTCATAGCCGTCAC | CCAGGGTTATGGAGCAGTTCA | 18.61 | Y | 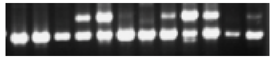 |
| 200 | SRIP-33496 | pig_ctg12:46976540-46976541(-) | ATTTGCTCGTCTCGCCTGAT | ACCAATAAAAACCATGCCCTGC | 7.46 | Y | 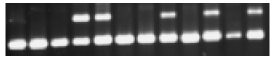 |
| 201 | SRIP-57133 | pig_ctg6:31353975-31353976(-) | GGTCCCTCCGCTTAGTTCAC | CCGGGATCGACCTCTCTTTG | 19.24 | Y | 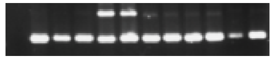 |
| 202 | SRIP-33285 | pig_ctg12:39096752-39096753(-) | TCATGTCAGGCCAGCATTTAGT | GGCCGAATGGAAGACCAACA | 11.92 | Y | 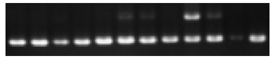 |
| 203 | SRIP-56989 | pig_ctg6:20901703-20901704(+) | CTTAAGACGAGTTTCCCAGGATGA | TGGGTTCCTACACCCTTTGGC | 7.32 | Y | 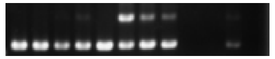 |
| 204 | SRIP-55271 | pig_ctg5:48678062-48678063(-) | TCAGGATTGGACCAGCTAAAGT | ATAGGTTTCTGGCAGGCATGG | 17.08 | Y | 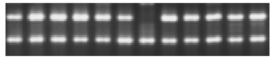 |
| 205 | SRIP-28099 | pig_ctg1:165082020-165082022(-) | CGGGTCACGACCTACTTGTG | AAGGCCTAGTACCACCGAGT | 8.64 | N | 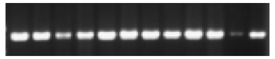 |

| | | | | | | | |
|---|---|---|---|---|---|---|---|
| 206 | SRIP-29967 | pig_ctg10:14230819-14230820(+) | TCTTCATTCCCCTGTAGCCC | TAACTGCTGAGCAACGACTGTT | 14.36 | Y | 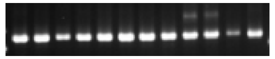 |
| 207 | SRIP-60961 | pig_ctg7:102638137-102638139(+) | TTTCAAGGCAAGGGAAGCCA | CTGGTGGGCCTGGAAAGAAT | 9.72 | / | 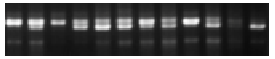 |
| 208 | SRIP-26006 | pig_ctg1:6755113-6755125(+) | GGTAGCAAATGGATGGACAGTG | ACGTATCAACTACGCCATCCG | 5.1 | / | 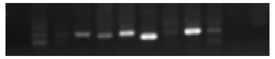 |
| 209 | SRIP-63236 | pig_ctg8:128099053-128099054(+) | AGTTTGCCTGGCTATCACTTCT | GGATGCTGGATGAAGGCAGTTA | 9.67 | Y | 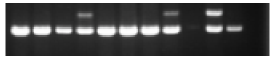 |
| 210 | SRIP-31799 | pig_ctg11:29574524-29574526(-) | GTTAAAAGCACACCCAAAGGTCA | CAATTTCATAAGTTAATGCGGGGAG | 6.39 | Y | 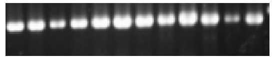 |
| 211 | SRIP-41341 | pig_ctg16:7338049-7338050(-) | ACAACGGACTACAGATTGATCC | TGTTTACTAATTCTCCCCAAGGCT | 18.8 | Y | 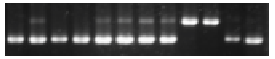 |
| 212 | SRIP-39339 | pig_ctg15:16951697-16951698(+) | TGCAGCAGTGTTACTCAAAATGG | CTCTGCCCCTCACACTTCAG | 7.47 | N | 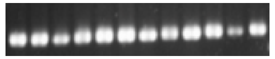 |
| 213 | SRIP-28975 | pig_ctg1:243211075-243211076(-) | AGGAAAGCTGAAGGTACTCAGTC | TGACTGTGAGGTTGTTGACG | 6.09 | N | 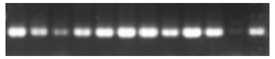 |
| 214 | SRIP-58387 | pig_ctg6:131017314-131017315(+) | ATGACCTCTTCCTCAGTCCGT | GTAGATGAAGTCATGGATTGTCAAG | 6.5 | N | 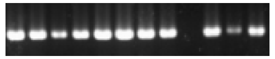 |
| 215 | SRIP-30095 | pig_ctg10:18763345-18763346(+) | GTTCCCCCGGGTAGTCAAAT | TGCGAGTAGCTTAGTGTCGTC | 13.97 | Y | 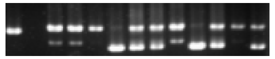 |
| 216 | SRIP-44682 | pig_ctg18:46330724-46330725(+) | CCAAATATGAGAGCGACTGGC | CCCCAAACTCTGAAGCGTG | 5.87 | Y | 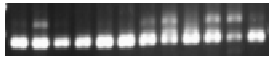 |
| 217 | SRIP-46650 | pig_ctg19:112619620-112619621(+) | CCTCATTCCCCGGCTTATGT | CCGTCCCTAAGATGTGTGCG | 16.76 | Y | 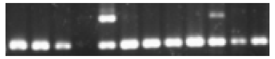 |
| 218 | SRIP-55353 | pig_ctg5:57477015-57477016(-) | GCACATAAGCTCCAAGCACAG | ACGGTCCATTATAGGTAAGACGG | 14.91 | Y | 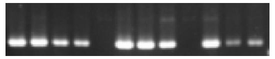 |

| | | | | | | | |
|---|---|---|---|---|---|---|---|
| 219 | SRIP-64491 | pig_ctg9:43766587-43766588(+) | CCAGAGTAAGCTCCCCTCCT | AGGTCAGGGCTGAGACTAGG | 14.01 | Y | 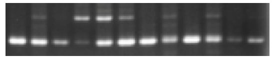 |
| 220 | SRIP-44925 | pig_ctg19:758900-758901(+) | TACAACGTACCACAAAGGCGT | AAGGCTACCTACGCTCCATTT | 9.25 | N | 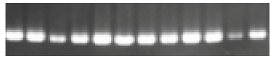 |
| 221 | SRIP-53963 | pig_ctg4:120136141-120136142(-) | GCTCCTAGAGGGCTAACAAGC | CAACGGAAATCTCAACATGGGG | 5.43 | Y | 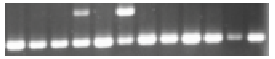 |
| 222 | SRIP-48642 | pig_ctg2:126100587-126100588(-) | GCGTGTACACACCATCAGTCT | GCAAGCCAATATGGTTCCCAC | 14.87 | Y | 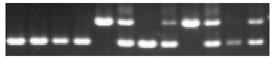 |
| 223 | SRIP-29693 | pig_ctg10:3794726-3794727(+) | TATAACAGTTGCTGTGGCGGAA | GCCGAATATGAGGACATAACAGC | 6.37 | Y | 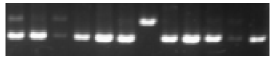 |
| 224 | SRIP-50830 | pig_ctg3:68487183-68487184(+) | GTTTCCCCATGTACCCGGAAG | AGCGGGCATAAGGATGAAGT | 8.33 | N | 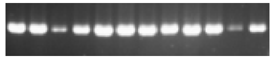 |
| 225 | SRIP-48910 | pig_ctg2:137739128-137739129(-) | GCCCAGAATAGTACCTGCCA | CCTCTCAGCAGTCCCAAAGG | 9.1 | Y | 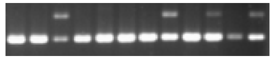 |
| 226 | SRIP-40531 | pig_ctg15:105764199-105764200(-) | CAAGCATTAGGAAGGCATTGAAA | ACCTCATGCCTTTCAGCTCG | 18.85 | Y | 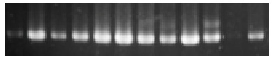 |
| 227 | SRIP-32998 | pig_ctg12:22795981-22795982(+) | CTTCCTACTCCCAGTTTTGATTGC | TGGGTAAGGCTGTTTTCACCT | 8.71 | Y | 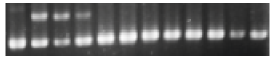 |
| 228 | SRIP-37871 | pig_ctg14:63438739-63438740(-) | TGCAGAGATACTGGCTGCTAAG | TAAGATTGGCAGACTCCAGGG | 5.3 | Y | 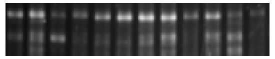 |
| 229 | SRIP-50327 | pig_ctg3:26773381-26773382(+) | TGTTCCCACAGCACTTCGTATT | TGATACTACCAACACGCGAAG | 8.29 | Y | 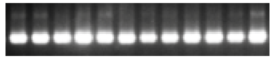 |
| 230 | SRIP-62029 | pig_ctg8:37947682-37947683(+) | GTTGTCTCTGGGGAGTTCGC | CATCCTTTCTCCAGGGGATTGAA | 8.29 | Y | 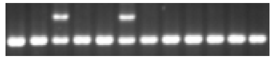 |
| 231 | SRIP-47001 | pig_ctg2:9564059-9564060(-) | CACACGTTTCCTGGTGCAAAT | TGAAAATGCGTGAATCTCAGAAGG | 16.67 | Y | 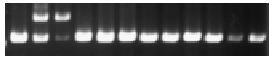 |

| | | | | | | | |
|---|---|---|---|---|---|---|---|
| 232 | SRIP-56075 | pig_ctg5:89866370-89866371(+) | TCTGGGGACAATCACTCGGT | AGCATGCCATTTGTAGAAGCTA | 17.8 | Y | 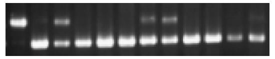 |
| 233 | SRIP-58623 | pig_ctg6:145215338-145215339(+) | CCCTCTAGCATAGGGGGCAT | TAACGTTGAAGAGCTGGCAAT | 14.58 | Y | 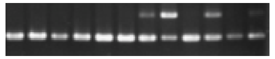 |
| 234 | SRIP-64074 | pig_ctg9:24416215-24416216(-) | CCTACATGGGTACAAGCTCCTC | GCGTATGCGAGAGGAATAAGC | 5.83 | Y | 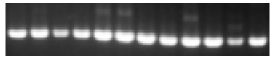 |
| 235 | SRIP-26200 | pig_ctg1:13852846-13852847(+) | TACCATTCTCTCTAGGACGTTGTAT | TTATGCTCCCCTGACCTCGT | 9.32 | Y | 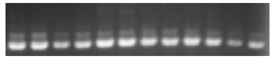 |
| 236 | SRIP-62309 | pig_ctg8:67451725-67451728(-) | TTCGTTTGCCCCAGAGTTCA | GAGATCCTGGCAGTGATGGG | 5.83 | Y | 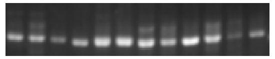 |
| 237 | SRIP-57588 | pig_ctg6:70424994-70424995(+) | CTGCAAGAATCAAGACCACGG | CAGGGGATCCCGAGGTACAG | 17.93 | Y | 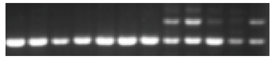 |
| 238 | SRIP-62371 | pig_ctg8:72639566-72639567(+) | CAGAGCAGGGCCATTCTTCA | TGCACACCACATACCGAGTC | 9.1 | N | 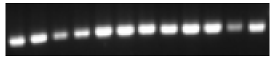 |
| 239 | SRIP-46551 | pig_ctg19:105918805-105918806(-) | AGGAGATCTTCCGTCCATACTCA | TCTGGGATAGTTGACTCTTGAACTG | 7.12 | Y | 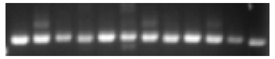 |
| 240 | SRIP-64914 | pig_ctg9:80194867-80194869(+) | TAAGAGACAGAGCAAGGAACCAC | ACTTCCCACATACTACCCTGG | 9.06 | Y | 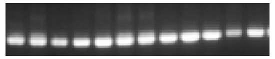 |
| 241 | SRIP-28648 | pig_ctg1:221143088-221143089(+) | CGCGAAGTGTTAGCGACCT | GGCATCTTCCTTACCACGGC | 8.93 | Y | 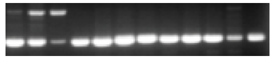 |
| 242 | SRIP-42865 | pig_ctg17:13070152-13070153(+) | CACCGTGATCCTTATCGGGAC | GGGCAACCCTGTCTCAGAA | 14.71 | Y | 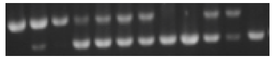 |
| 243 | SRIP-36382 | pig_ctg13:190132364-190132365(-) | TAGGAAAATGACATTGTGCAGC | ACAGAAATATAAACTCCTGCCTCA | 6.73 | Y | 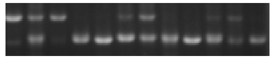 |
| 244 | SRIP-59838 | pig_ctg7:28620484-28620485(-) | CTTTGCCTGCTAGCTTGAACC | CTTCTAACATAGGGTCGTCATGGAT | 15.48 | Y | 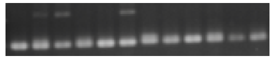 |

| | | | | | | | |
|---|---|---|---|---|---|---|---|
| 245 | SRIP-29626 | pig_ctg10:545522-545523(+) | ACCGTTTGCAGAGAGACGAT | ACGGCAAGCGACAGAACATT | 7.4 | Y | 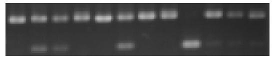 |
| 246 | SRIP-51144 | pig_ctg3:93211666-93211667(-) | TACACACTGGTGAAATGGCA | AGATGCACTTTTACTGACACAGTC | 19.84 | Y | 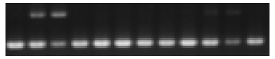 |
| 247 | SRIP-33016 | pig_ctg12:23279015-23279016(+) | AGCGGTGTACAATGACCTAAAGT | GTCTTTCCCATTCTTCAAAGGAGC | 5.18 | Y | 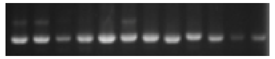 |
| 248 | SRIP-62533 | pig_ctg8:83070065-83070066(+) | AAGAACACCAACGACCTCCA | GGCTGAATTGGTGGGAACTC | 5.74 | Y | 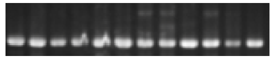 |
| 249 | SRIP-43679 | pig_ctg17:51409766-51409767(+) | GGACTCAAGCTCAACTCTGC | TGCCGACATGAAGTATTTTGCC | 7.18 | Y | 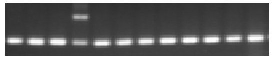 |
| 250 | SRIP-32425 | pig_ctg11:69395647-69395648(-) | TAGTGGGTCACAGAGGACCG | CGGGTGATGTTTGAGAAGCAC | 6.52 | Y | 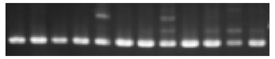 |
| 251 | SRIP-33147 | pig_ctg12:30829249-30829250(+) | CAGGCCTGGAACCTAAAAAGAG | CCACCTGCATTAGTATTTTGAGCA | 5.31 | Y | 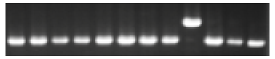 |
| 252 | SRIP-44408 | pig_ctg18:33456727-33456728(-) | AGCAACTGGGGACGAACTAT | TAAATGCCCACACCCCATACC | 6.3 | / | 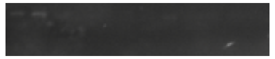 |
| 253 | SRIP-32095 | pig_ctg11:53427413-53427414(-) | GTTCTGGTCTAGCTGGCTTTACT | GGCTCCAAGGAAAAGTGACCA | 7.59 | Y | 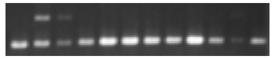 |
| 254 | SRIP-41552 | pig_ctg16:18316725-18316726(-) | GTTGTGGATGACAATCCGTGG | TGGACTGTCCTGGCTAACTTC | 17.51 | Y | 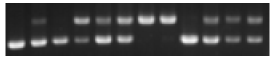 |
| 255 | SRIP-39089 | pig_ctg15:1776271-1776272(-) | AAGGACCCCTAGACCCTTAG | TGCAACCAATAAGGTGGAGGA | 16.57 | Y | 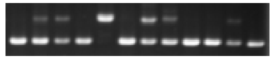 |
| 256 | SRIP-34105 | pig_ctg13:10479847-10479848(-) | AAACATCACGCAGGACTGGAA | GTTTTAGGTGCTCATCAGCCTTT | 47.78 | Y | 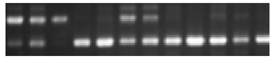 |
| 257 | SRIP-64709 | pig_ctg9:61823417-61823418(+) | CAGCACAAACCATTTCATGGC | GCTTCCAATCCAAGACAAATACA | 25.9 | N | 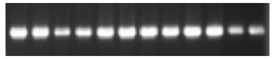 |

| | | | | | | | |
|---|---|---|---|---|---|---|---|
| 258 | SRIP-39268 | pig_ctg15:13757493-13757494(-) | TGCCTAGAGTGACAGGTAGC | GCAGTGCAGTCCAACAAACT | 34.17 | Y | 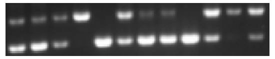 |
| 259 | SRIP-50016 | pig_ctg3:14487972-14487973(+) | CCTTCCTCCTCCGCTTGTTG | ATGCCAGCCTTCATACGGTT | 21.31 | Y | 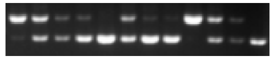 |
| 260 | SRIP-54457 | pig_ctg498:35049-35050(-) | GCTTTCCAATGCTCCTCGTTTT | CAGCTGATTTGAATACCCTTACCG | 45.81 | N | 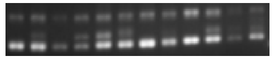 |
| 261 | SRIP-33736 | pig_ctg12:56582475-56582476(-) | CTGGGGACTTGGCTGTTTCT | AACGTGCGACAGCCTGATAA | 24.66 | Y | 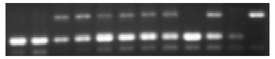 |
| 262 | SRIP-28836 | pig_ctg1:232929968-232929969(-) | CCCTGCTACAGAGTCCCTTG | AGCACCCAGTAAAAGCCAACT | 49.73 | Y | 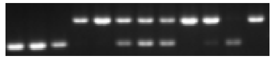 |
| 263 | SRIP-35209 | pig_ctg13:89191691-89191692(-) | TTTGCTGTGAGAACCTAGCCAA | ACAACTCGGGAGACCAGGTG | 28.85 | Y | 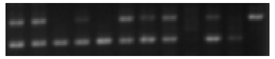 |
| 264 | SRIP-65451 | pig_ctg9:121135231-121135232(+) | CCGTTCGTCACAGTCACACG | ACACTCCAAACCTGAAACGCC | 36.85 | Y | 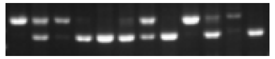 |
| 265 | SRIP-40104 | pig_ctg15:72003234-72003235(+) | TGTTGTACCCTTGTCTGGAACA | CACTCAAAAGTTCCAGATGGAAAGA | 33.62 | Y | 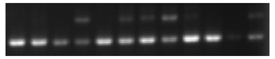 |
| 266 | SRIP-40795 | pig_ctg15:120879105-120879106(+) | GTTTGTTAGGCACGCTGTGA | GCCCAGCGTGATAGAGTTGA | 40.36 | Y | 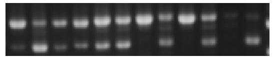 |
| 267 | SRIP-40939 | pig_ctg15:127570537-127570538(+) | GAGCGATGATAGCTGCTGAGT | CCCAGCCCTACCTTGAGTTT | 40.98 | Y | 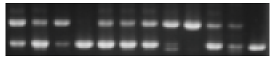 |
| 268 | SRIP-41755 | pig_ctg16:25929147-25929148(-) | TCACAGACAACCTCGGTGC | CATCTCCGGGACGTGTTGAT | 46.33 | Y | 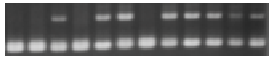 |
| 269 | SRIP-60517 | pig_ctg7:77247974-77247975(-) | ACCTTATAGAGGGCGAGCTG | GAGAGGCTACTTCATTCTTTTTGAA | 28.73 | Y | 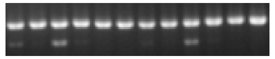 |
| 270 | SRIP-40922 | pig_ctg15:126853433-126853434(+) | CCAGCACCATGTGATGAGCA | TCCTAAACCCTCACCAATAAACAGT | 58.17 | Y | 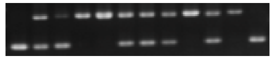 |

| | | | | | | | |
|---|---|---|---|---|---|---|---|
| 271 | SRIP-28364 | pig_ctg1:191281960-191281961(+) | AGGTTTGGCTTCCCTCTTACT | CACCTTACCACACAGTACCCA | 28.55 | Y | 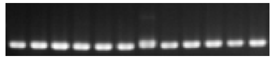 |
| 272 | SRIP-49503 | pig_ctg223:372367-372368(+) | AGGCGTAGAACACATCCGTG | TCGATGCTCGTTATTAGCAGGT | 26.83 | Y | 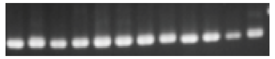 |
| 273 | SRIP-49753 | pig_ctg3:3538279-3538280(-) | AGCTTTGAGATGGGTTCAGGAAT | GCAAGTGTTTTCCCATGAGGC | 27.2 | Y | 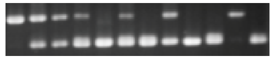 |
| 274 | SRIP-55256 | pig_ctg5:47228292-47228293(-) | GGCGAACGTTCAGAGTTTTGAT | TCCTGCCAACCAATAACGACT | 30.57 | N | 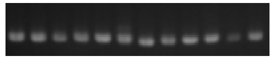 |
| 275 | SRIP-29017 | pig_ctg1:246292267-246292268(+) | ACCTTCAAGCTTTGTGTGGG | CATCTGTGGGCATTCGATTTCA | 54.33 | N | 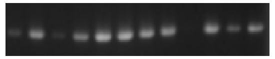 |
| 276 | SRIP-58203 | pig_ctg6:115560644-115560645(+) | AACTATTATTGGGGGAGGCACC | GCTTTAGCCTCTGGGGTGTA | 21.6 | Y | 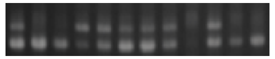 |
| 277 | SRIP-53261 | pig_ctg4:77036711-77036712(+) | CCTCCGCCTCAAAGAAAGAC | AAGGCTGGGGATCATTACGG | 27.38 | Y | 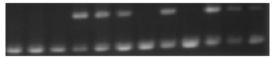 |
| 278 | SRIP-37751 | pig_ctg14:55339779-55339780(-) | ACTCCCACATCTCAAAGTGCAA | CCAGCTAGGCTCCTTCATCC | 28.1 | Y | 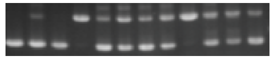 |
| 279 | SRIP-50047 | pig_ctg3:15749652-15749653(+) | TCTGAAAGCCTCCTTGGTGTC | GGATGGGGAGAGTTACAGTACA | 34.45 | Y | 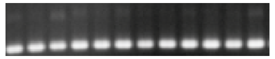 |
| 280 | SRIP-58689 | pig_ctg6:147386276-147386278(-) | GTGAACCACTCACAGAGCCTT | ACACTGGAATCTTTGAATGCGG | 35.83 | / | 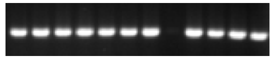 |

Fig S1
